# In silico unwinding of *Caenorhabditis elegans* microRNA duplexes to evaluate thermodynamic end stabilities improves predictions of microRNA strand selection

**DOI:** 10.1101/2025.06.26.661848

**Authors:** Jeffrey C. Medley, Anna Zinovyeva

## Abstract

microRNAs (miRNAs) are endogenous ∼22 nucleotide long, non-coding RNAs that post-transcriptionally regulate gene expression. During miRNA biogenesis, stem-loop-containing miRNA precursors are enzymatically cleaved to form a small RNA duplex. Cleavage positions are determined based on the position of structural motifs and junctions on the stem-loop precursor. The duplex end containing a favorable 5’ nucleotide and lower thermodynamic stability is subsequently loading into an Argonaute protein. Typically, one duplex (guide) strand is retained in Argonaute and becomes functional whereas the other (passenger) strand is degraded. Therefore, accurate structural predictions of miRNA intermediates and quantification of duplex end stabilities are important towards understanding miRNA biogenesis. Here, we compiled predicted secondary structures for all *Caenorhabditis elegans* miRNA hairpins and duplexes at physiologically relevant temperatures. We developed a new approach to calculate the thermodynamic stability of miRNA duplex ends, which resulted in improved predictions of miRNA strand selection. Our approach introduces hard constraints to folding algorithms to restrict base-paring of terminal nucleotides, which improves modeling of *in vivo* duplex end unwinding. We propose that constrained RNA folding can be used to evaluate local stabilities within an RNA secondary structure.

## Introduction

While approximately two-thirds of the human genome is actively transcribed into RNA, less than 3% of the human genome appears to encode for proteins (Djebali et al. 2012; Dunham et al. 2012). A large fraction of transcribed loci corresponds to non-coding RNAs, which can be broadly categorized into two major classes according to their length: long non-coding RNAs (typically defined as ≥ 200 nucleotides long) and small non-coding RNAs (≤ 200 nucleotides) (Chen and Kim 2024). The secondary structure of non-coding RNA molecules often defines their biological activities (Mortimer et al. 2014). Dysregulation of non-coding RNAs, including mutations expected to impact secondary structure, have been reported in several human diseases including cancer (French and Edwards 2020; Nemeth et al. 2024; Wan et al. 2014; Zhou et al. 2023). Long non-coding RNAs can have extensive secondary or tertiary structures that contribute to diverse biological functions including regulation of gene expression, control of protein activity, or serving as scaffolds for the assembly of biomolecular complexes (Kopp and Mendell 2018; Wang and Chang 2011). Some small non-coding RNAs, such as ribosomal RNAs (rRNAs) or transfer RNAs (tRNAs), have a secondary structure that is critical for their biological activities. Other classes of small non-coding RNAs, including microRNAs (miRNAs), play repressive roles in gene expression by guiding the RNA-induced silencing complex (RISC) to specific mRNA targets (Jouravleva and Zamore 2025; Shi et al. 2022).

As miRNAs rely on intermolecular base pairing to identify mRNA targets, an extensive secondary structure would likely be prohibitive towards target recognition. However, during miRNA biogenesis, the secondary structure of miRNA precursors plays critical roles in generating the correct mature miRNA sequence. Primary miRNAs (pri-miRNAs) are genomically encoded as single-stranded RNA species that contain a self-complementary stem-loop segment (Cai et al. 2004; Lee et al. 2004). During canonical miRNA biogenesis, the microprocessor complex cleaves pri-miRNAs at a defined position by measuring the length of the base-paired stem relative to unpaired structural junctions on the pri-miRNA to produce an ∼70 nucleotide hairpin precursor miRNA (pre-miRNA) (Denli et al. 2004; Gregory et al. 2004; Han et al. 2004; Kwon et al. 2019; Landthaler et al. 2004; Lee et al. 2003; Nguyen et al. 2023, 2015; Partin et al. 2020). Additional sequence features and motifs have also been identified that influence the accuracy of microprocessor cleavage (Auyeung et al. 2013; Fang and Bartel 2015; Li et al. 2020a, 2020b). Dicer subsequently cleaves the pre-miRNA to remove the terminal loop from the hairpin and generate an ∼22 nucleotide miRNA duplex (Bernstein et al. 2001; Grishok et al. 2001; Hutvágner et al. 2001; Ketting et al. 2001; Knight and Bass 2001). In addition to structural elements of pre-miRNAs, Dicer recognizes sequence features including the GYM or YCR motifs to identify the correct cleavage position (Gu et al. 2012; Feng et al. 2012; Le et al. 2024a, 2024b; Lee et al. 2023a, 2023b; MacRae et al. 2006; Park et al. 2011; Tsutsumi et al. 2011). The miRNA duplex is then loaded into an Argonaute protein, and a single, mature strand (guide/miR strand) is retained in the Argonaute, while the other strand (passenger/miR* strand) is degraded (Elbashir et al. 2001; Grishok et al. 2001; Hammond et al. 2001; Martinez et al. 2002; Mourelatos et al. 2002; Nykänen et al. 2001). The MID domain of Argonaute prefers to load the duplex end containing a more favorable 5’ nucleotide (U>A>G>C) and lower thermodynamic end stability (Frank et al. 2010; Khvorova et al. 2003; Schwarz et al. 2003; Suzuki et al. 2015).

However, these guidelines are not sufficient to describe the strand preference of ∼25% of miRNA duplexes (Medley et al. 2021), suggesting that additional cis (structural, thermodynamic or sequence-based) features or trans-acting factors likely influence miRNA strand choice. Thus, accurate calculations of free energy within miRNA duplexes may facilitate identification of additional features impacting strand selection.

Understanding the various thermodynamic and structural elements influencing miRNA biogenesis relies on accurate predictions of RNA secondary structure and stability. As RNA folding is largely dependent on intramolecular base-pairing, the secondary structure of RNA molecules can be predicted from their primary nucleotide sequence (Fallmann et al. 2017; Lorenz et al. 2016). Prediction of RNA secondary structure relies on identifying the minimal free energy (MFE) structure of the folded RNA, which can be calculated using experimentally derived parameters for neighboring nucleobase-stacking effects on the thermodynamic stability of different RNA helices (Andronescu et al. 2014; Fallmann et al. 2017; Mathews et al. 1999; Schroeder and Turner 2000; Xia et al. 1998; Zuber et al. 2022). The nearest neighbor energy model has been used for algorithmic prediction of RNA secondary structure, including by the widely used ViennaRNA package (Lorenz et al. 2011). The nearest neighbor model uses an additive sliding window approach to calculate free energy, where the thermodynamic stability of dinucleotide pairs is summed across the length of an RNA structure (Lorenz et al. 2011; Mathews et al. 1999; Schroeder and Turner 2000). The free energy of RNA is also directly linked to temperature, and temperature-sensitive differences in RNA folding allow for alternative functions of ’RNA thermometers’ in bacteria, yeast and plants (Kortmann and Narberhaus 2012; Righetti et al. 2016; Su et al. 2018; Thomas et al. 2022; Wan et al. 2012). The default parameters of RNA folding algorithms typically assume the RNA should be folded at 37°C but allow for user-defined changes in folding temperature (Lorenz et al. 2011). As 37°C is outside the physiological range of many organisms including the nematode *Caenorhabditis elegans*, using default parameters for these algorithms may lead to inaccurate predictions of RNA secondary structure. Notably, the current versions of the miRNA repositories miRBase v22.1 (Kozomara et al. 2018) and MirGeneDB v3.0 (Clarke et al. 2024) often disagree with their predicted structures of *C. elegans* miRNA precursors (Figure S1).

In this study, we used RNAfold and RNAcofold from the ViennaRNA package (Bernhart et al. 2006; Hofacker et al. 1994; Lorenz et al. 2011) to predict the folding of *C. elegans* miRNA hairpins and duplexes at physiologically relevant temperatures. We show that more than 25% of *C. elegans* miRNA hairpins have increased predicted base-pairing at 20°C compared to the default temperature of 37°C. Most of the predicted changes in base-pairing occurred at the terminal ends, although in some cases we observed altered positions of central bulges and mismatches at lower temperatures. We found that the predicted MFE structures of ∼15% of miRNA duplexes also showed increased predicted base-pairing at 20°C. We also observed differences in the predicted folding pattern of miRNA hairpins and miRNA duplexes across the physiological range of *C. elegans* temperatures, including differences that would be expected to impact miRNA processing. Interestingly, the predicted MFE structures for ∼30% of miRNA duplexes were partially unwound relative to the hairpin-derived miRNA duplex structure, suggesting that unwinding of many miRNA duplex ends is energetically favorable once the duplex is liberated from the hairpin structure. As miRNA strand selection is connected to thermodynamic asymmetry of duplex ends (Khvorova et al. 2003; Schwarz et al. 2003; Suzuki et al. 2015), energetically favorable unwinding may contribute to proper miRNA strand choice. To quantify the thermodynamic stability of miRNA duplex ends, we developed a new approach that calculates the stability of partially unwound duplex structures by introducing hard constraints into the folding algorithm (Lorenz et al. 2016). We use these constraints to calculate the difference in the free energy of partially unwound duplexes and the fully wound duplex structures. We refer to the difference in the free energy as unwinding energy, which is proportional to the amount of energy that would be required to unwind that duplex end.

Unwinding energy modeled thermodynamic asymmetry of miRNA duplex ends and outperformed free energy values obtained from the nearest neighbor database (Turner and Mathews 2010) at predicting strand selection of *C. elegans* miRNAs.

Importantly, our analysis provides an updated catalog of *C. elegans* miRNA structures and end stabilities at physiologically relevant temperatures, which may improve discovery of sequence features influencing miRNA biogenesis and strand selection. Furthermore, our *in silico* method of duplex unwinding allows for evaluation of thermodynamic asymmetries within miRNA duplex ends and may have broader use for assessing local stabilities within different RNA structures.

## Results

### Secondary structure predictions of *C. elegans* miRNA hairpins at different temperatures

To improve upon the current publicly available miRNA hairpin folds (Clarke et al. 2024; Kozomara et al. 2018), we used RNAfold to predict structures of miRNA hairpins (Table S1) across the physiological range of *C. elegans* (15°C, 20°C and 25°C) compared to the traditional default temperature parameter (37°C). We acquired sequence information for all *C. elegans* miRNA hairpins from miRBase which included a mixture of pre-miRNA sequences and extended pri-miRNA hairpin sequences when available (Kozomara et al. 2018). We found that more than 25% of *C. elegans* pre-miRNAs were predicted to fold differently at 20°C, a temperature commonly used for laboratory culture than at the non-physiological temperature 37°C (Table S1). Most of the temperature-dependent differences that we observed occurred at the terminal end of the hairpin, although we also observed some examples where the position of central mismatches or bulges were affected (Figure 1A, Table S1). In some cases, the temperature-dependent changes in hairpin folding affected the locations of basal or apical junctions that are important for miRNA processing (Shang et al. 2023) (Figure 1B-C, Table S1). We also found that some hairpins were predicted to fold differently within the physiological range of *C. elegans* (Figure 1D-E, Table S1). Compared to 20°C, we observed alternative predicted folding for 3.5% of miRNA hairpins at 15°C and ∼8% of hairpins at 25°C (Table S1). These findings show that temperature should be considered when performing algorithmic prediction of miRNA hairpin structure, with parameters assessed for physiological relevance.

**Figure 1:**
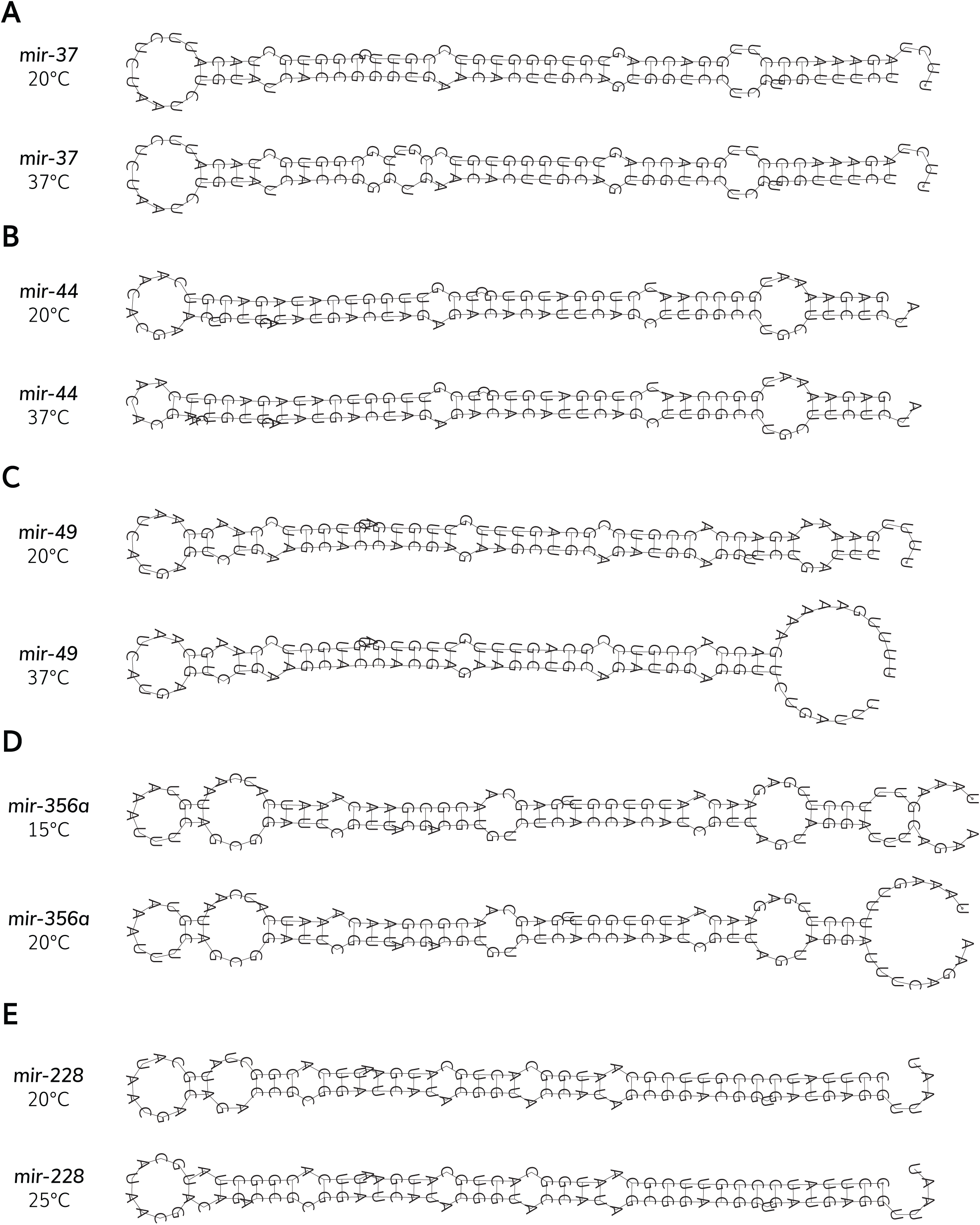
Differences in predicted miRNA hairpin folding across different temperatures. (A) Predicted folding of the *mir-37* hairpin at 20°C (top) and 37°C (bottom) shows altered locations of central mismatches and bulges at different temperatures. (B) The position of the apical junction in *mir-44* is shifted at 20°C (top) compared to 37°C (bottom). (C) The location of the *mir-49* basal junction is altered at 20°C (top) relative to 37°C (bottom). (D) The predicted folding of the *mir-356a* hairpin suggests different folding is favored at the physiological temperatures 15°C (top) and to 20°C (bottom). (E) The *mir-228* hairpin is expected to adopt a different structure at the physiological temperatures 20°C (top) and to 25°C (bottom). (A-E) Minimal free energy (MFE) structures of miRNA hairpins were acquired from RNAfold. miRNA hairpin sequences were obtained from miRBase v22.1 (Kozomara et al. 2018).

### Predicted MFE structures of *C. elegans* miRNA duplexes at different temperatures

We next used RNAcofold to examine how temperature influences the MFE structures of *C. elegans* miRNA duplexes (Table S2). We obtained sequences for available miRNA guide and passenger strands from miRBase and generated MFE structures for 190 miRNA duplexes that have annotated guide and passenger strand sequences (Table S2). Compared to the physiologically relevant temperature 20°C, we found that 13.2% of miRNA duplexes have an altered MFE structure at the non-physiological temperature 37°C. In general, higher temperature was associated with decreased base-pairing of miRNA duplexes, especially at terminal duplex ends (Figure 2, Table S2). In some miRNA duplexes, we observed temperature-dependent changes in the positions of central mismatches/bulges (Figure 2A, Table S2), which play a role in sorting small RNAs to specific Argonautes (Jannot et al. 2008; Seroussi et al. 2023; Steiner et al. 2007). In other duplexes, increased temperature was predicted to result in partial unwinding of one duplex end (Figure 2B-D, Table S2), leading to altered predictions of thermodynamic asymmetries important for small RNA strand selection (Khvorova et al. 2003; Schwarz et al. 2003). We also observed cases where higher temperature affected the overall structure of mismatched nucleotides (Figure 2E, Table S2) or resulted in rearrangements that affected both duplex ends and central mismatches (Figure 2F, Table S2). Similar to miRNA hairpins, alternative folding of miRNA duplexes was also predicted to occur within the physiological temperature range of *C. elegans* (Figure 2G-H, Table S2). Compared to 20°C, temperature-dependent differences in duplex structure were observed for 4.2% of miRNAs at 15°C and 4.7% of miRNAs at 25°C (Figure 2G-H, Table S2). These findings show that, similar to miRNA hairpins, temperature should be considered when performing algorithmic prediction of miRNA duplex structure, with parameters assessed for physiological relevance.

**Figure 2:**
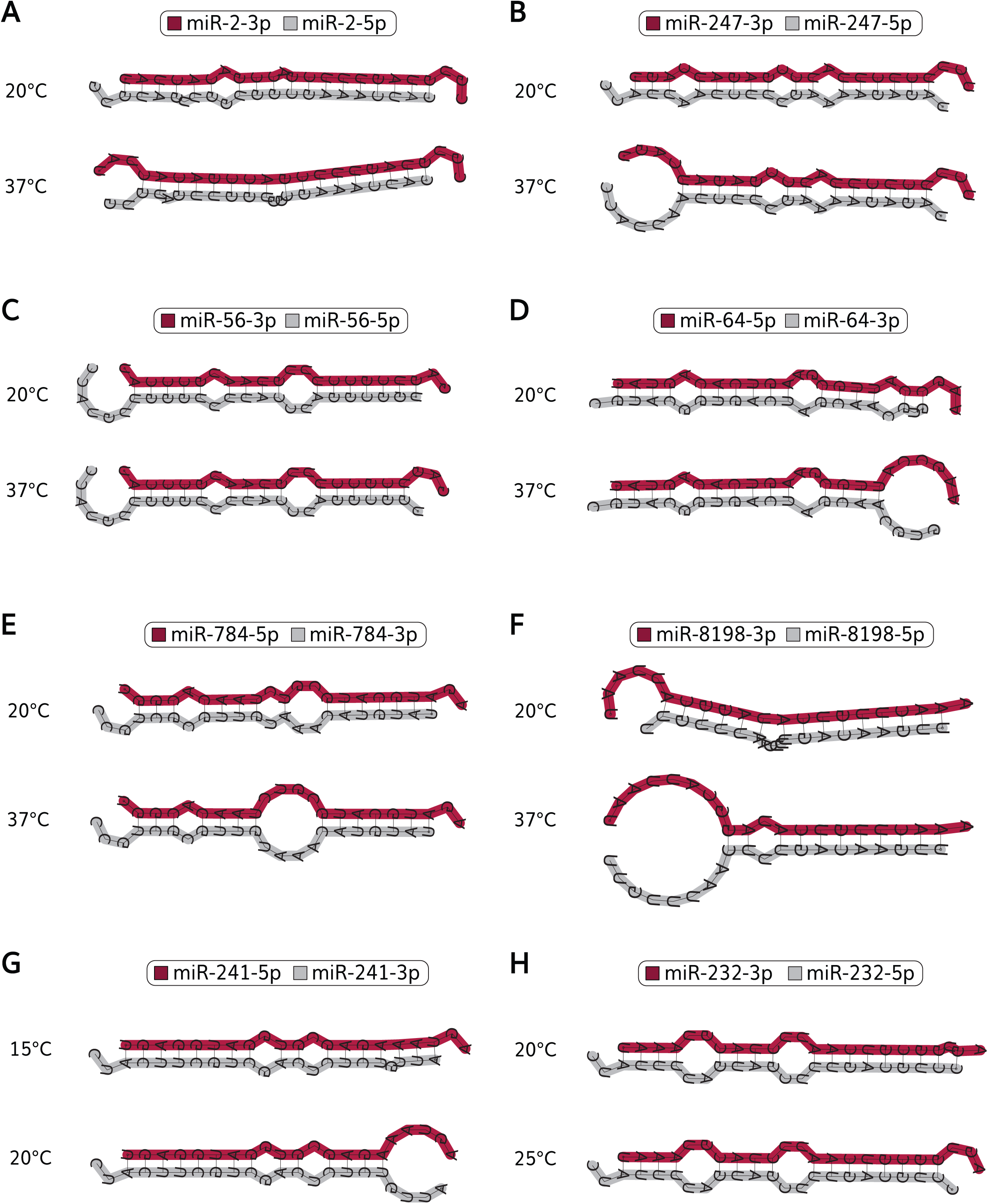
Differences in minimal free energy miRNA duplex structures at different temperatures. (A) Minimal free energy (MFE) structures of the miR-2 duplex at 20°C (top) and 37°C (bottom). The expected positions of central mismatches and bulges are affected by different temperature conditions. (B) MFE structures of the miR-247 duplex at 20°C (top) and 37°C (bottom). The 3p end (guide end) of the miR-247 duplex is predicted to be partially unwound at higher temperature. (C) MFE structures of the miR-56 duplex at 20°C (top) and 37°C (bottom). The 5’ nucleotide of miR-56-5p (passenger strand) is predicted to become unpaired at higher temperature. (D) MFE structures of the miR-64 duplex at 20°C (top) and 37°C (bottom). The 3p end (passenger end) of the miR-64 duplex is predicted to be partially unwound at higher temperature. (E) MFE structures of the miR-784 duplex at 20°C (top) and 37°C (bottom). At lower temperature, a U:A base pair intervenes a centrally located loop structure to create two adjacent loops. At 37°C, the U:A base pair is disrupted to generate a single, larger loop structure. (F) MFE structures of the miR-8198 duplex at 20°C (top) and 37°C (bottom). The 3p end (guide end) of the miR-8198 duplex is predicted to be partially unwound at higher temperature and the location of central mismatches/bulges are shifted. (G) MFE structures of the miR-241 duplex at 15°C (top) and 20°C (bottom). At 20°C, the 3p end (passenger end) of the miR-241 duplex is predicted to be partially unwound while increased base-pairing is predicted at 15°C. (H) MFE structures of the miR-232 duplex at 20°C (top) and 25°C (bottom). The 5’ nucleotide of miR-56-5p (passenger strand) is predicted to be paired at 20°C and become unpaired at 25°C. (A-H) Minimal free energy (MFE) structures of miRNA hairpins were acquired from RNAcofold. Guide strands are shown on the top of each duplex structure and highlighted in maroon. Passenger strands are shown on the bottom of each duplex structure and highlighted in gray. miRNA 5p and 3p sequences and guide strand information were obtained from miRBase v22.1 (Kozomara et al. 2018).

### miRNA hairpin precursors constrain miRNA duplexes into a suboptimal energy state

When we examined the MFE structures of *C. elegans* miRNA duplexes, we noticed that the base-pairing of the MFE structure often differed from the base-pairing of the equivalent nucleotides in the hairpin structure at the same temperature (Table S3). At 20°C, nearly one-third (31.1%) of miRNA MFE duplex structures were different from their predicted structure within the hairpin (Table S3). We observed differences in both the location of central mismatches and terminal nucleotide pairing (Figure 3, Table S3). We quantified the positions of mismatched or bulged nucleotides, which revealed decreased base-pairing of terminal nucleotides in the MFE duplex structure compared to the hairpin-derived duplexes (Figure S2, Table S3). We found that many of the duplex ends that became unpaired in the MFE structure corresponded to miRNA guide strands (Figure 3, Figure S2), which could contribute to thermodynamic asymmetry of duplex ends. In some cases, unpairing of terminal nucleotides appeared to result in more extensive unwinding of that duplex end (Figure 3). Our analysis suggested that miRNA hairpins constrain duplexes into less energetically favorable states and that partial unwinding of some miRNA duplex ends is energetically favorable once the duplex is liberated from the hairpin precursor. This difference in stability between the hairpin-constrained duplex and MFE structure may contribute to thermodynamic asymmetries of duplex ends.

**Figure 3:**
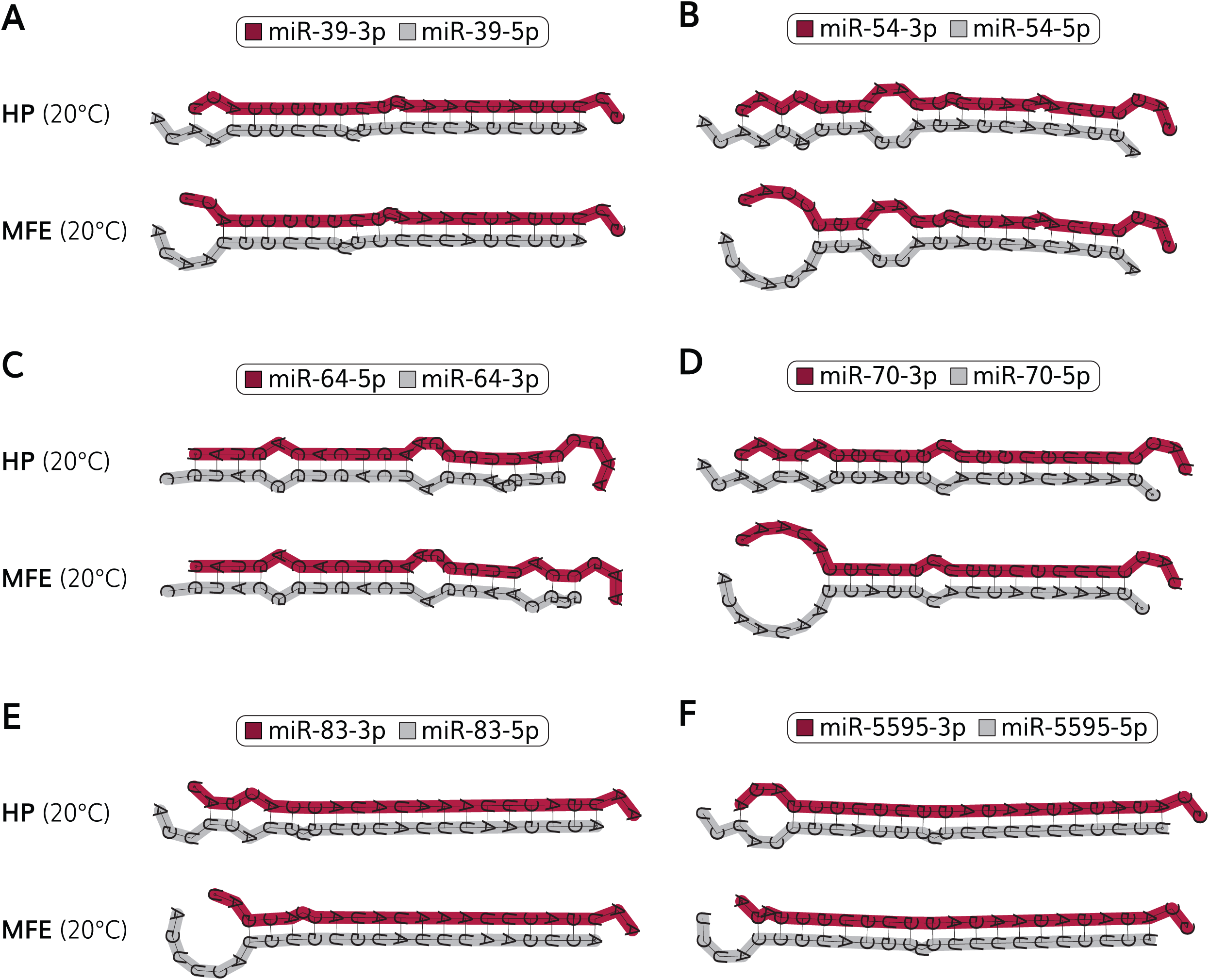
miRNA hairpins constrain duplexes into less energetically favorable states (A) Differences in the miR-39 hairpin-derived duplex (top) and MFE duplex (bottom) structures at 20°C. The 5’ nucleotide of miR-39-3p (guide strand) is predicted to paired in the hairpin, but is energetically favored to unwind in the MFE structure. (B) Differences in the miR-54 hairpin-derived duplex (top) and MFE duplex (bottom) structures at 20°C. Two nucleotides that are paired in the miR-54 hairpin are energetically favored to unwind in the 3p end (guide end) of the MFE duplex leading to unwinding of the 3p duplex end. (C) Differences in the miR-64 hairpin-derived duplex (top) and MFE duplex (bottom) structures at 20°C. The positions of mismatches/bulges in the 3p end (passenger end) of the miR-64 are favored to rearrange in the MFE structure relative to how they are positioned in the hairpin structure. (D) Differences in the miR-70 hairpin-derived duplex (top) and MFE duplex (bottom) structures at 20°C. Three nucleotides that are paired in the miR-70 hairpin are energetically favored to unwind in the 3p end (guide end) of the MFE duplex leading to extensive unwinding of the 3p duplex end. (E) Differences in the miR-83 hairpin-derived duplex (top) and MFE duplex (bottom) structures at 20°C. Two nucleotides that are paired in the miR-83 hairpin are energetically favored to unwind in the 3p end (guide end) of the MFE duplex leading to unwinding of the 3p duplex end and rearrangement of a mismatched nucleotide pair to a bulged nucleotide. (F) Differences in the miR-5595 hairpin-derived duplex (top) and MFE duplex (bottom) structures at 20°C. The 3p duplex end (guide end) of the miR-5595 is favored to rearrange so that the 5’ nucleotide in the MFE structure becomes unpaired. (A-F) Minimal free energy (MFE) structures of miRNA hairpins were acquired from RNAcofold. Hairpin structures were acquired by introducing folding constraints to mirror the duplex structure within the MFE hairpin that was acquired using RNAfold. Guide strands are shown on the top of each duplex structure and highlighted in maroon. Passenger strands are shown on the bottom of each duplex structure and highlighted in gray. miRNA 5p and 3p sequences and guide strand information were obtained from miRBase v22.1 (Kozomara et al. 2018).

### *In silico* unwinding of miRNA duplexes quantifies how unwound terminal nucleotides impact duplex stability

Previous methods to quantify duplex end stability have used the nearest neighbor model that uses energy parameters obtained at 37°C (Khvorova et al. 2003; Medley et al. 2021; Schwarz et al. 2003; Suzuki et al. 2015). As 37°C is a non-physiological temperature for *C. elegans*, we developed an alternative method for quantifying duplex end stability that uses physiological temperature parameters for *C. elegans* miRNA duplexes. Our approach was to calculate the stability of partially unwound duplexes compared to the fully wound MFE or hairpin-derived duplex structures (Figure 4A). We refer to the difference in free energy (ΔΔG) of the partially unwound duplex and the fully wound state as the ’unwinding energy’ for each duplex end, which should be proportional to the amount of energy required to unwind that end of the duplex (Figure 4A). An advantage of this approach is that it considers how unwinding terminal nucleotides impacts overall duplex stability, which is not necessarily considered when using values from nearest neighbor databases. To calculate unwinding energies, we first used RNAcofold to quantify the stability of each *C. elegans* miRNA duplex at 20°C. We then introduced hard folding constraints to RNAcofold (Lorenz et al. 2016) that prevented base-pairing of the terminal 1-4 nucleotides on each duplex end and determined the free energy of the partially unwound duplex structures (Figure 4A). The constraint files also instructed RNAcofold how to enforce base-pairing throughout the rest of the duplex, which allowed us to recapitulate in unwinding of MFE (Figure 4B, Table S4) or hairpin-derived (Figure 4C, Table S5) duplex structures. Using this method, a ΔΔG value of zero was obtained when the corresponding nucleotides were already unpaired in the MFE duplex structure, supporting that additional energy would not be required to unwind those terminal nucleotides (Figure 4B-C, Tables S4-S5). On the other hand, a negative ΔΔG indicated that it was energetically favorable for the duplex to unwind the corresponding nucleotides, which was often observed for the non-MFE, hairpin-derived duplex structures (Figure 4C, Table S5). As expected, the average unwinding energy of duplex ends corresponding to 5’ end of guide strands was lower than the duplex ends corresponding to the 5’ end of passenger strands, indicating that less energy would be required to unwind the 5’ guide end of the duplex (Figure 4B-C, Tables S4-S5). Therefore, our calculations of unwinding energy illustrate the intrinsic thermodynamic asymmetry of miRNA duplex ends.

**Figure 4:**
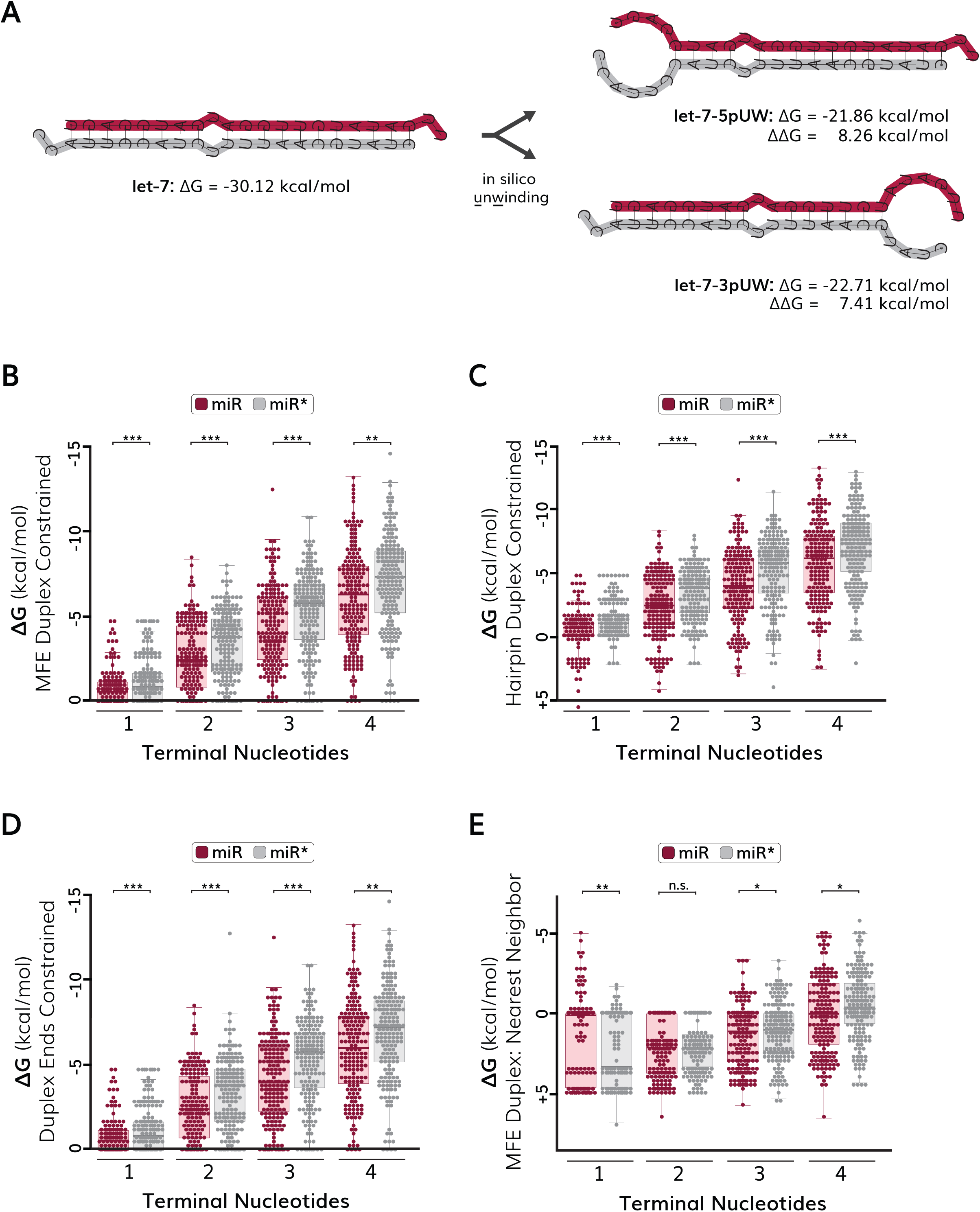
In silico unwinding of *C. elegans* miRNA duplexes to evaluate thermodynamic stability of duplex ends (A) Example of in silico unwinding (UW) method for the let-7 miRNA. The stability of the MFE or hairpin duplex is calculated using RNAcofold (left) and folding constraints are introduced to calculate the stability of the partially unwound duplex (right). The stability of each duplex end is calculated by taking the difference in free energy (ΔΔG) of the fully wound duplex and the partially unwound duplex. The let-7 guide strand (let-7-5p) is shown on the top of duplexes in maroon highlight, and the let-7 passenger strand (let-7-3p) is shown on the bottom of duplexes in gray highlight. (B) Quantification of thermodynamic end stability using in silico unwinding of constrained MFE duplexes. Duplexes were folded according to the MFE structure, or with the first one through four nucleotides of each miRNA duplex end forced to be unpaired. (C) Quantification of thermodynamic end stability using in silico unwinding of hairpin-constrained duplexes. Duplexes were folded according to the structure embedded within the miRNA hairpin, or with the first one through four nucleotides of each miRNA duplex end forced to be unpaired. (D) Quantification of thermodynamic end stability using in silico unwinding of unconstrained duplexes. Duplexes were allowed to refold into the MFE state following unwinding of the first one through four nucleotides of each miRNA duplex end. (E) Quantification of thermodynamic end stability using nearest neighbor values. Duplexes were folded according to the MFE structure, and values for nearest neighbor energy were acquired from the nearest neighbor database (Turner and Mathews 2010). Note that nearest neighbor values are provided at the 37°C, a non-physiological temperature for *C. elegans*. The stability of the rest of the duplex is not considered. (B-E) The guide end of the duplex (miR) is shown in maroon color and the passenger end of the duplex (miR*) is shown in gray color. Each dot represents a miRNA duplex end (n=190). Values for ΔΔG are given as kcal/mol. Boxes range from the first to third quartile of the data. The thick bar represents the statistical median. Lines extend to the minimum and maximum data point excluding outliers that were defined as 1.5 times the interquartile range. Statistical analysis: *p<0.05, **p<0.01, ***p<0.001, n.s. (not significant) p>0.05. (B-D) A value of zero indicates no difference in free energy between the wound and partially unwound duplexes and corresponds to duplex ends that are already unwound in the MFE or hairpin structures. A positive value indicates that the wound structure is less stable than the partially unwound structure.

We next examined whether partial unwinding of duplex ends may lead to unstable intermediate structures that are energetically favored to further unwind or otherwise undergo structural rearrangements. To address this, we introduced new folding constraints that prevented base-pairing of terminal nucleotides but did not instruct RNAcofold to retain the predicted hairpin-derived nucleotide pairing, which allowed the partially unwound duplexes to reconfigure into MFE structures. We refer to these structures as ’end-constrained’, as only the duplex ends are constrained during folding. Our quantification of unwinding energy revealed that the MFE state of some duplexes changes after partial unwinding, suggesting that unwinding of duplex ends could lead to progressive unwinding or structural rearrangements for some miRNAs (Figure 4D, Table S6). When compared to energy values from the nearest neighbor tables (Figure 4E, Table S7) (Turner and Mathews 2010), we found that our unwinding energy models outperformed nearest neighbor values at predicting the expected thermodynamic asymmetry of miRNA duplex ends (Figure 4B-D). In fact, the nearest neighbor model suggested that the terminal nucleotide was more stable for *C. elegans* duplex ends corresponding to the 5’ end of guide strands than the 5’ end of passenger ends (Figure 4E, Table S7), which contradicts previous studies of small RNA asymmetry (Khvorova et al. 2003; Schwarz et al. 2003; Suzuki et al. 2015). It is worth noting that the parameters of the nearest neighbor tables were determined at 37°C (Turner and Mathews 2010), which may lead to inaccurate representation of the thermodynamic asymmetry of *C. elegans* miRNA duplexes at physiological temperatures. We conclude that *in silico* unwinding of miRNA duplexes improves models of thermodynamic asymmetry of miRNA duplex ends and allows duplex end stability to be evaluated at a defined temperature condition. Overall, our annotation of miRNA duplex unwinding energies provides an improved resource quantifying *C. elegans* miRNA end stabilities.

### Prediction of miRNA strand selection using unwinding energies of miRNA duplex ends

To investigate whether different methods of calculating the thermodynamic stability of miRNA duplex end affected predictions of miRNA strand selection, we used the twin-drive model (Suzuki et al. 2015) to calculate the predicted strand choice of *C. elegans* miRNAs. The twin-drive model uses a formula that weighs the relative effects of 5’ nucleotide identity (N-drive) and thermodynamic asymmetry (G-drive) to calculate the expected levels of each strand (Suzuki et al. 2015). The twin-drive formula is given as ln(5p/3p) = kΔΔG_5p-3p_ + N_5p_-N_3p_, where k and N are constants affecting thermodynamic stability and nucleobase identity respectively (Suzuki et al. 2015). We used the twin-drive formula to predict strand selection of *C. elegans* miRNAs using different methods of calculating thermodynamic stability for the first through fourth nucleotides on each end of the duplex (Tables S8-S11). Using the unwinding energy of MFE duplexes, we were able to correctly predict strand selection for 75.3% of miRNA duplexes when considering the thermodynamic stability of only the single terminal nucleotide on each duplex end (Figure 5A-B, Table S8). When we calculated thermodynamic stability using additional nucleotides on each duplex end, we found that thermodynamic asymmetry was less predictive of miRNA strand selection (Figure 5A-B, Figure S3A-B, Table S8). For example, when considering the unwinding energy of the three terminal nucleotides on each duplex end, we were able to correctly predict strand selection for 71.1% of miRNAs (Figure 5A-B). Predictions of miRNA strand selection were similar when using the unwinding energy of hairpin-derived duplexes (Figure 5C, Figure S3C, Table S9), or end-constrained duplexes (Figure 5D, Figure S3D, Table S10), although predictions using the unwinding energy of MFE duplexes slightly outperformed both alternative folding methods (Figure 5). Therefore, different methods of calculating unwinding energy appear to have similar predictive value for determining miRNA strand preference. As a comparison, we next used nearest neighbor energy values to predict strand selection of *C. elegans* miRNAs (Figure 5E, Figure S3E, Table S11). We found that nearest neighbor stabilities predicted strand selection equally as unwinding energies of MFE duplexes when considering the stabilities for three or four terminal nucleotides on each duplex end (Figure 5E, Figure S3E). However, nearest neighbor stabilities were less predictive of strand selection when considering one or two terminal nucleotide stabilities (Figure 5E, Figure S3E). While the unwinding energy of the single terminal nucleotides for MFE duplexes correctly predicted strand selection of 75.3% of miRNAs, the nearest neighbor stabilities of terminal nucleotides were only successful at predicting strand selection of 58.4% of miRNAs (Figure 5, Figure S3). Thus, it appears that unwinding energies better model thermodynamic energies of duplex ends when considering few terminal nucleotides, possibly because our unwinding energies approach considers how unwinding of terminal ends affects the overall stability of the partially unwound duplex. Collectively, our results support that unwinding energies model thermodynamic asymmetry of miRNA duplex ends and are predictive of miRNA strand selection in *C. elegans*.

**Figure 5:**
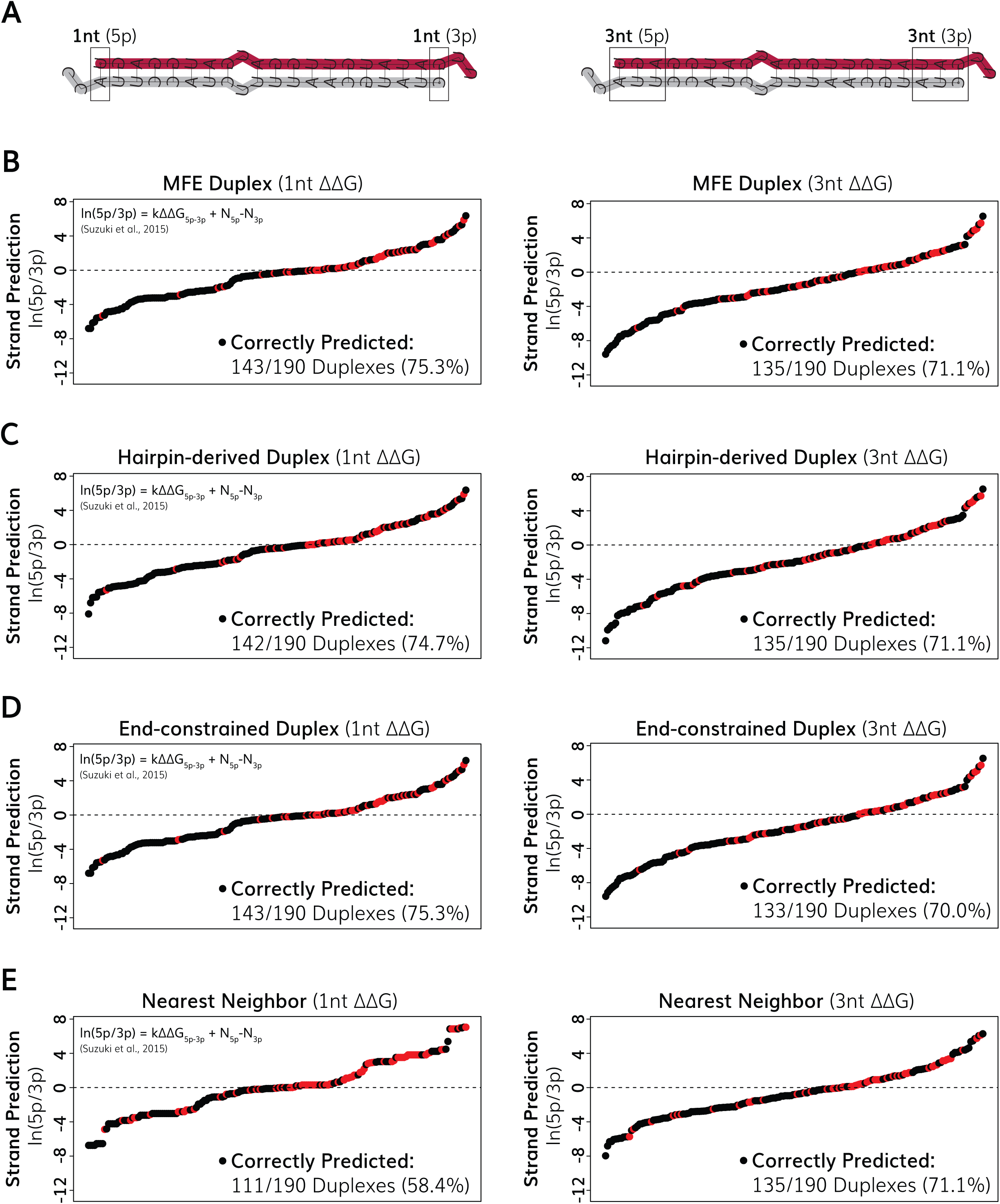
Predictions of miRNA strand selection. (A) Example let-7 duplexes illustrating nucleotides considered for thermodynamic stability calculations. The terminal nucleotides (left) or three terminal nucleotides (right) were considered. Ranked-order plots of predicted *C. elegans* miRNA strand asymmetry for (B) MFE duplex structures using unwinding energy, (C) hairpin-derived duplex structures using unwinding energy, (D) End-constrained duplexes using unwinding energy and (E) nearest neighbor energy values. (B-E) Predicted strand ratios were determined using a formula from the twin-drive model [ln(5p/3p) = kΔΔG_5p-3p_ + N_5p_-N_3p_], using previously described values for constants (Suzuki et al. 2015). miRNAs are plotted in order of lowest to highest predicted ln(5p/3p) values. miRNAs above dashed line are predicted 5p dominant and miRNAs below dashed line are predicted 3p dominant. Correct predictions (based on guide strands reported in miRBase) are indicated as black dots, and incorrect predictions are indicated as red dots.

## Discussion

### Structural predictions of *C. elegans* miRNA intermediates

Structural features of miRNA intermediates are critical for accurate miRNA biogenesis (Shang et al. 2023), yet structural repositories for miRNA structure information are often in disagreement (Figure S1). In this study, we explored different in silico folding methods to predict the structure of *C. elegans* miRNA hairpins and duplexes and calculate duplex end stabilities. Our analysis shows that temperature parameters of RNA folding algorithms have a substantial impact on predicted structures of miRNA intermediates. Previous studies, including our own (Medley et al. 2021), have used default temperature parameters (37°C) to predict the structure of *C. elegans* miRNA intermediates. However, as temperature is directly linked to free energy, folding *C. elegans* miRNAs at non-physiological temperatures may lead to inaccurate predictions of secondary structures. We show that performing RNA folding within the physiological temperature range of *C. elegans* (15-25°C) leads to increased predicted base-pairing of *C. elegans* miRNA intermediates. We also observed structural differences when miRNA intermediates were folded across different temperatures within the physiological range of *C. elegans*, including altered positions of structural junctions that are important for miRNA processing (Shang et al. 2023). It is intriguing to speculate that temperature may regulate miRNA-dependent gene regulation in a manner akin to ’RNA thermometers’ that have been described in other species (Kortmann and Narberhaus 2012; Righetti et al. 2016; Su et al. 2018; Thomas et al. 2022; Wan et al. 2012). However, additional studies will be important to uncover whether temperature-dependent differences in miRNA folding predictions accurately reflect physiological folding and are functionally relevant.

During our analysis, we noticed that the base-paring of MFE miRNA duplex structures was often different than the base-paring of the equivalent nucleotides within the miRNA hairpin precursor. These differences likely arise from additional structural stability that is conferred from neighboring base-pairs that are present in the hairpin precursor but are removed during miRNA processing events. This observation suggests that miRNA hairpins, at least temporarily, constrain some miRNA duplexes into suboptimal energy states. As thermodynamic asymmetries of miRNA duplex ends are associated with miRNA strand selection (Khvorova et al. 2003; Schwarz et al. 2003; Suzuki et al. 2015), it is possible that some miRNA hairpins “prime” miRNA duplexes for asymmetric Argonaute loading and subsequent miRNA strand selection.

Energetically favorable rearrangements of miRNA duplexes to reach their MFE states may promote unwinding of miRNA duplex ends and influence strand choice. Alternatively, miRNA duplexes may have sufficient time to rearrange into their MFE structures before miRNA strand selection occurs, in which case it is unlikely that hairpin-constrained duplex folding contributes towards thermodynamic asymmetries influencing strand choice. Additional work will be needed to clarify whether the base-paring status of miRNA duplexes within their hairpin precursors influences miRNA stand biogenesis.

### Evaluating thermodynamic asymmetry by in silico unwinding improves predictions of miRNA strand choice

We developed an *in silico* method to assess the thermodynamic stability of miRNA duplex ends using physiologically relevant temperature parameters. We introduced hard constraints to the RNA folding algorithm to restrict terminal nucleotide base pairing and calculate the stability of partially unwound duplex structures. By comparing the difference in stability of partially and fully wound MFE structures, we determined the unwinding energy of each duplex end. We reasoned that the unwinding energy should be proportional to the amount of energy required to unwind each duplex end. We used unwinding energy to predict miRNA strand asymmetry using the previously described twin-drive model (Suzuki et al. 2015). We found that unwinding energy outperformed free energy values obtained from the nearest neighbor database (Turner and Mathews 2010), especially when considering the free energy one or two terminal nucleotides. As miRNA strand selection is often disrupted in human disease (reviewed in (Medley et al. 2021)), improved predictions of miRNA strand selection may help better understand the connection between miRNA strand choice and human diseases. Notably, we were able to correctly predict the preferred strand for ∼75% of *C. elegans* miRNAs, suggesting that the twin-drive model is not sufficient to explain miRNA strand selection of all miRNAs. It seems likely that additional features of miRNA duplexes influence strand selection of *C. elegans* miRNAs *in vivo*.

Locally stable RNA structures can have important functions, and algorithms have been developed to identify locally structured regions of RNA molecules (Waldl et al. 2023). We used our in silico unwinding method to assess the local thermodynamic stability of miRNA duplex ends. In principle, this approach could be used to evaluate local stabilities of secondary structures within other RNA molecules. As some RNA binding proteins may recognize locally structured regions of their target RNA molecules (Bravo et al. 2018), unwinding energies may help identify regions that may facilitate interactions with RNA binding proteins. Alternatively, it is possible that intermolecular interactions, between RNA and proteins or other nucleic acids, may locally restrict base-pairing across specific nucleotides of a given RNA molecule. In such case, constrained folding may improve predictions of how RNA secondary structure is altered upon intermolecular interactions and the energy requirement needed to facilitate structural rearrangements.

A current disadvantage of our approach is the time investment required to generate constraint files for folding algorithms based on careful manual curation of predicted structures. Towards that end, user-friendly automation of our in silico unwinding approach will substantially reduce the time required to perform such analyses and facilitate similar analyses across different RNA molecules and organisms. Collectively, our current analysis provides a careful manual curation of the structures and stabilities of *C. elegans* miRNA intermediates, a valuable resource towards understanding the mechanisms of miRNA biogenesis, strand selection, and, ultimately, miRNA-dependent gene regulation.

## Materials and Methods

### Secondary structure predictions of *C. elegans* pre-miRNAs and miRNA duplexes

The sequences of *C. elegans* pre-miRNAs and mature miRNAs were retrieved from miRBase v22.1 (Kozomara et al. 2018). RNAfold was used to predict pre-miRNA hairpin structures and RNAcofold was used to predict miRNA duplex structures (Bernhart et al. 2006; Hofacker et al. 1994; Lorenz et al. 2011). We were unable to predict the duplex structure of miRNAs that lacked annotations for the passenger strand sequence. The parameter "-T 20" was used to predict RNA folding at 20°C, or "-T X" (where X is a specific temperature) was used to predict RNA folding at alternative temperatures. The input files used for *C. elegans* pre-miRNAs (File S1) and miRNA duplexes (File S2) are provided in the supplementary material. miRNA duplex structures were recolored using Adobe Illustrator such that the guide strand is maroon and the passenger strand is gray. Locations of mismatches and bulges were identified using dot-bracket nomenclature (Lorenz et al. 2011). Terminal 3’ nucleotide overhangs were not included when scoring internal mismatch/bulge locations.

### In silico unwinding of miRNA duplexes and predictions of strand selection

To calculate unwinding energies, folding constraints were introduced in RNAcofold to force unpaired nucleotides at the miRNA duplex ends: “-T 20 -C --enforceConstraint", which mimics partial unwinding of duplex ends. The input files containing constraint information are provided in the supplementary material (Files S3-S5). Unwinding energies for each duplex end were calculated by subtracting the energy of the fully wound (maximal free energy state) duplex from the partially unwound duplex (the first 1:4 nucleotides were unwound from each duplex end).

Guide and passenger strand annotations were determined based on which strand had the most reported reads on miRBase (Kozomara et al. 2018). For nearest neighbor stabilities, values were acquired from the nearest-neighbor database (Turner and Mathews 2010). Predictions of miRNA strand selection were determined using the twin-drive model equation [ln(5p/3p) = kΔΔG_5p-3p_ + N_5p_-N_3p_] using constants previously described (Suzuki et al. 2015).

## Statistical analysis

R statistical software (R Core Team, 2021) was used to calculate all statistics. All P-values were calculated using two-tailed t*-*tests assuming equal variance among samples. All statistics are presented as mean ± one standard deviation unless otherwise specified.

## Acknowledgements

We thank members of the Zinovyeva lab for helpful comments and discussions. This work was supported by grant from the National Institutes of Health (R35GM124828 to A.Z.), the Kansas INBRE program (P20 GM103418) and the Johnson Cancer Center at K.S.U. J.C.M was supported by a fellowship from the NIH (F32GM148040).

