## Supplementary Materials for "In silico unwinding of *Caenorhabditis elegans* microRNA duplexes to evaluate thermodynamic end stabilities improves predictions of microRNA strand selection"

**Figure S1:** Comparison of structural information obtained from central miRNA repositories. (A) Comparison of the (A) *let-7*, (B) *lin-4*, (C), *mir-1*, (D) *mir-2* and (E) mir-35 hairpin structures from miRBase v22.1 (left) and MirGeneDB 3.0 (right). (B) Structure of the lin-4 hairpin obtained from miRBase (left) and MirGeneDB (right). For miRBase structures, nucleotides contained within the duplex structure are given as pink, uppercase text. For MirGeneDB structures, guide strands are given as red text and passenger strands are indicated by blue text. Yellow highlighting indicates locations of motifs involved in miRNA processing.

**Figure S2:** Quantification of mismatched or bulged nucleotide frequency within miRNA duplexes based on the hairpin-derived duplex (A-D) or the MFE duplex (E-H) structures. (A-H) Percent of miRNAs containing mismatches (gray) or bulges (purple) at positions (x-axis) relative to the 5' end of the miRNA. 3' terminal nucleotide overhangs were excluded from this analysis. All *C. elegans* miRNA duplexes (n=190) with annotated guide and passenger strands were included in this analysis.

**Figure S3**: Predictions of miRNA strand selection. (A) Example let-7 duplexes illustrating nucleotides considered for thermodynamic stability calculations. The terminal two nucleotides (left) or four terminal nucleotides (right) were considered. Ranked-order plots of predicted *C. elegans* miRNA strand asymmetry for (B) MFE duplex structures using unwinding energy, (C) hairpin-derived duplex structures using unwinding energy, (D) End-constrained duplexes using unwinding energy and (E) nearest neighbor energy values. (B-E) Predicted strand ratios were determined using a formula from the twin-drive model [$\ln(5p/3p) = k\Delta\Delta G_{5p\text{-}3p} + N_{5p}\text{-}N_{3p}$], using previously described values for constants (Suzuki et al. 2015) . miRNAs are plotted in order of lowest to highest predicted ln(5p/3p) values. miRNAs above dashed line are predicted 5p dominant and miRNAs below dashed line are predicted 3p dominant. Correct predictions (based on guide strands reported in miRBase) are indicated as black dots, and incorrect predictions are indicated as red dots.

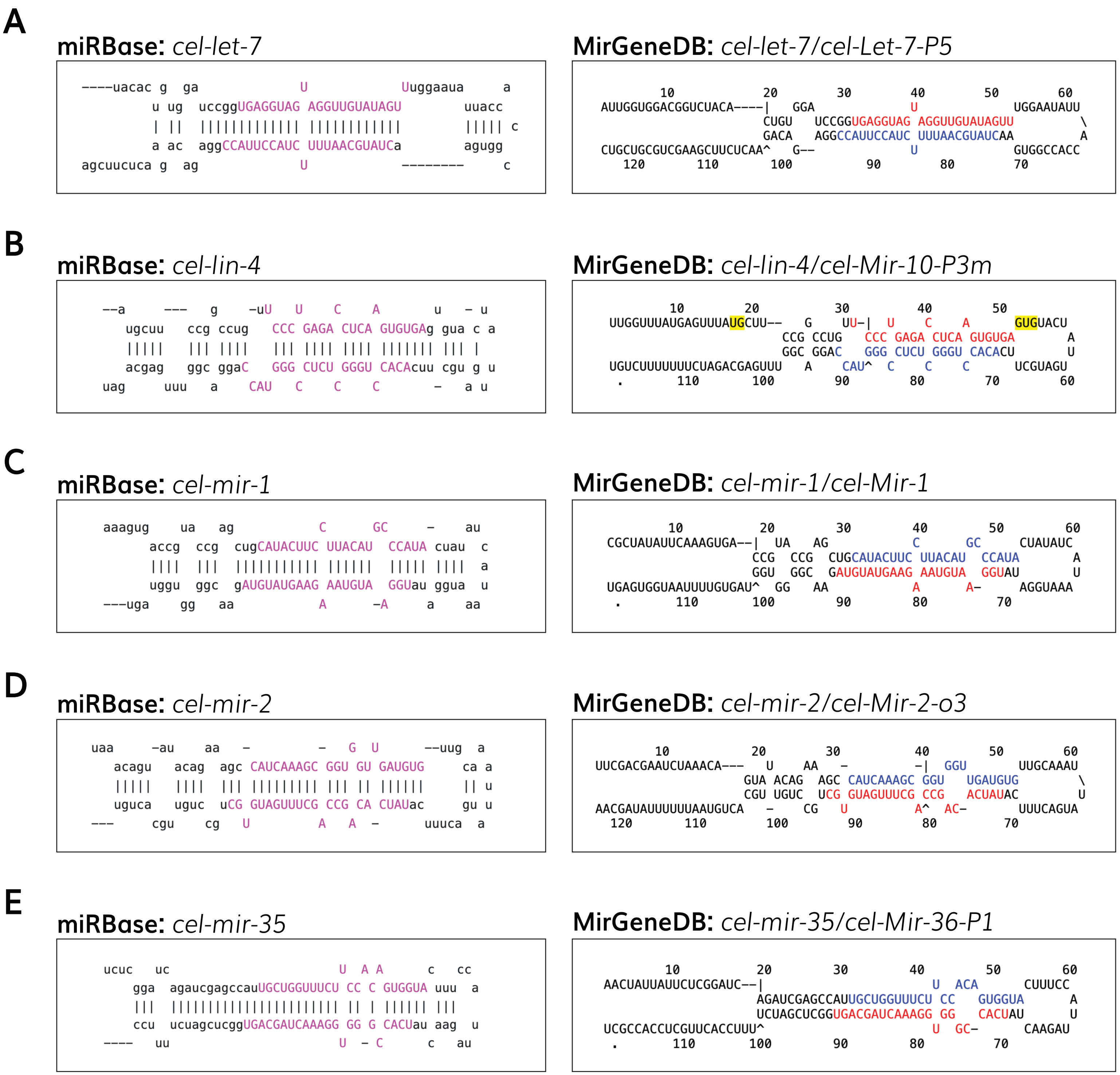
A
miRBase: cel-let-7
----uacac g ga U Uuggaaua a
u ug uccggUGAGGUAG AGGUUGUAUAGU uuacc
| || ||||||||||||| ||||||||||||| ||||| c
a ac aggCCAUUCCAUC UUUAACGUAUCa agugg
agcuucuca g ag U -------- c
MirGeneDB: cel-let-7/cel-Let-7-P5
10 20 30 40 50 60
AUUGGUGGACGGUCUACA----| GGA U UGGAAUAUU
CUGU UCCGGUGAGGUAG AGGUUGUAUAGUU \
GACA AGGCCAUUCCAUC UUUAACGUAUCAA A
CUGCUGCGUCGAAGCUUCUCAA^ G-- U GUGGCCACC
120 110 100 90 80 70
B
miRBase: cel-lin-4
--a --- g -uU U C A u - u
ugcuu ccg ccug CCC GAGA CUCA GUGUGAg gua c a
||||| ||| |||| ||| |||| |||| ||||||| ||| |
acgag ggc ggaC GGG CUCU GGGU CACAcuu cgu g u
uag uuu a CAU C C C - a u
MirGeneDB: cel-lin-4/cel-Mir-10-P3m
10 20 30 40 50
UUGGUUUAUGAGUUUAUGCUU-- G UU-| U C A GUGUACU
CCG CCUG CCC GAGA CUCA GUGUGA A
GGC GGAC GGG CUCU GGGU CACACU U
UGUCUUUUUUUCUAGACGAGUUU A CAU^ C C C UCGUAGU
. 110 100 90 80 70 60
C
miRBase: cel-mir-1
aaagug ua ag C GC - au
accg ccg cugCAUACUUC UUACAU CCAUA cuau c
|||| ||| ||||||||||| |||||| ||||| |||| a
uggu ggc gAUGUAUGAAG AAUGUA GGUau ggua u
---uga gg aa A -A a aa
MirGeneDB: cel-mir-1/cel-Mir-1
10 20 30 40 50 60
CGCUAUAUUCAAAGUGA--| UA AG C GC CUAUAUC
CCG CCG CUGCAUACUUC UUACAU CCAUA A
GGU GGC GAUGUAUGAAG AAUGUA GGUAU U
UGAGUGGUAAUUUUGUGAU^ GG AA A A- AGGUAAA
. 110 100 90 80 70
D
miRBase: cel-mir-2
uaa -au aa - - G U --uug a
acagu acag agc CAUCAAAGC GGU GU GAUGUG ca a
||||| |||| ||| ||||||||| ||| || |||||| || u
uguca uguc uCG GUAGUUUCG CCG CA CUAUac gu u
--- cgu cg U A A - uuuca a
MirGeneDB: cel-mir-2/cel-Mir-2-o3
10 20 30 40 50 60
UUCGACGAAUCUAAACA--- U AA - -| GGU UUGCAAAU
GUA ACAG AGC CAUCAAAGC GGU UGAUGUG \
CGU UGUC UCG GUAGUUUCG CCG ACUAUAC U
AACGAUAUUUUUUAAUGUCA - CG U A^ AC- UUUCAGUA
120 110 100 90 80 70
E
miRBase: cel-mir-35
ucuc uc U A A c cc
gga agaucgagccauUGCUGGUUUCU CC C GUGGUA uuu a
||| |||||||||||||||||||||||| || | |||||| |||
ccu ucuagcucggUGACGAUCAAAGG GG G CACUau aag u
---- uu U - C c au
MirGeneDB: cel-mir-35/cel-Mir-36-P1
10 20 30 40 50 60
AACUAUUAUUCUCGGAUC--| U ACA CUUUCC
AGAUCGAGCCAUUGCUGGUUUCU CC GUGGUA A
UCUAGCUCGGUGACGAUCAAAGG GG CACUAU U
UCGCCACCUCGUUCACCUUU^ U GC- CAAGAU
. 110 100 90 80 70

**Figure S1**

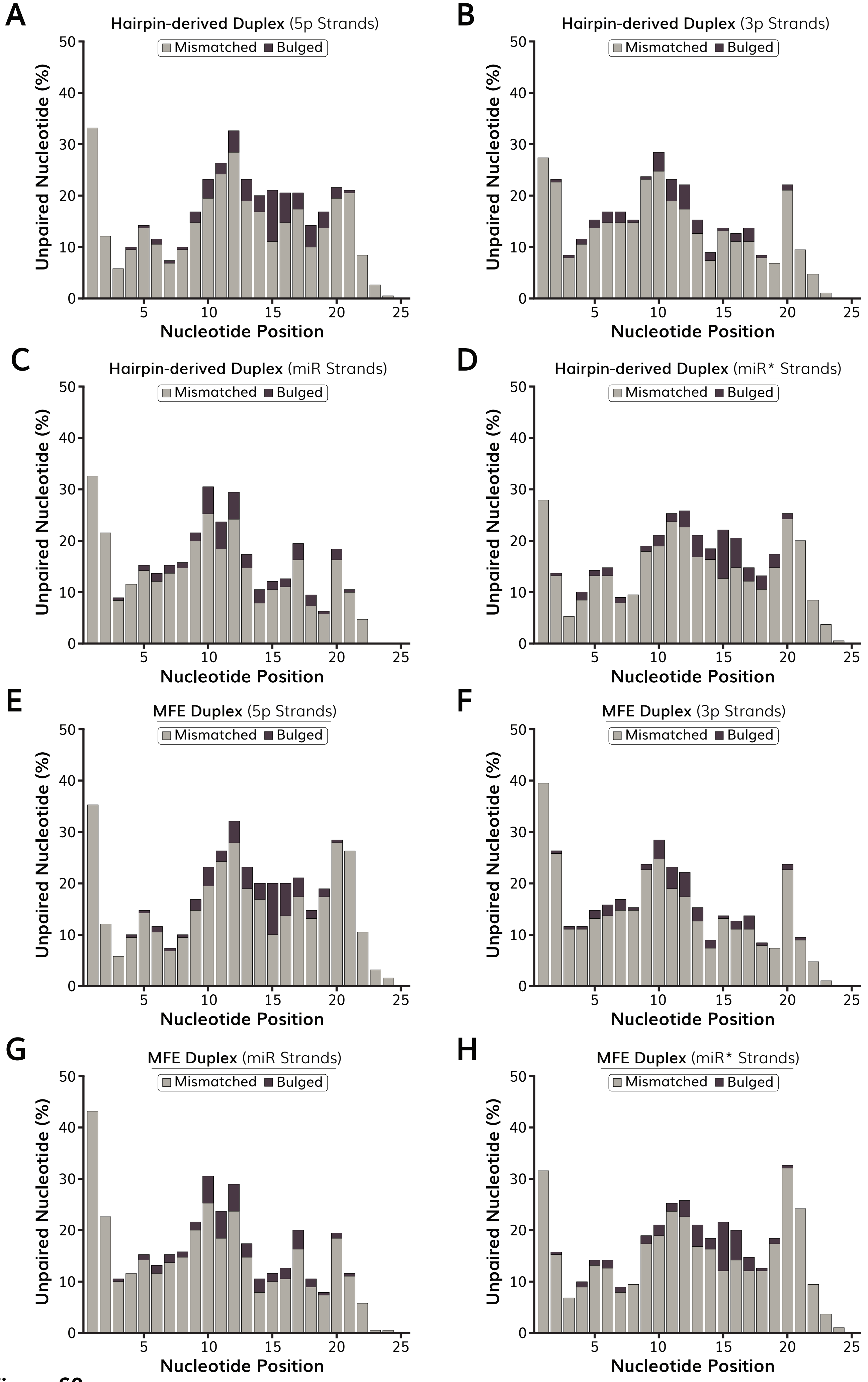

A
Hairpin-derived Duplex (5p Strands)
B
Hairpin-derived Duplex (3p Strands)
C
Hairpin-derived Duplex (miR Strands)
D
Hairpin-derived Duplex (miR* Strands)
E
MFE Duplex (5p Strands)
F
MFE Duplex (3p Strands)
G
MFE Duplex (miR Strands)
H
MFE Duplex (miR* Strands)
Mismatched
Bulged
Unpaired Nucleotide (%)
Nucleotide Position


Figure S2

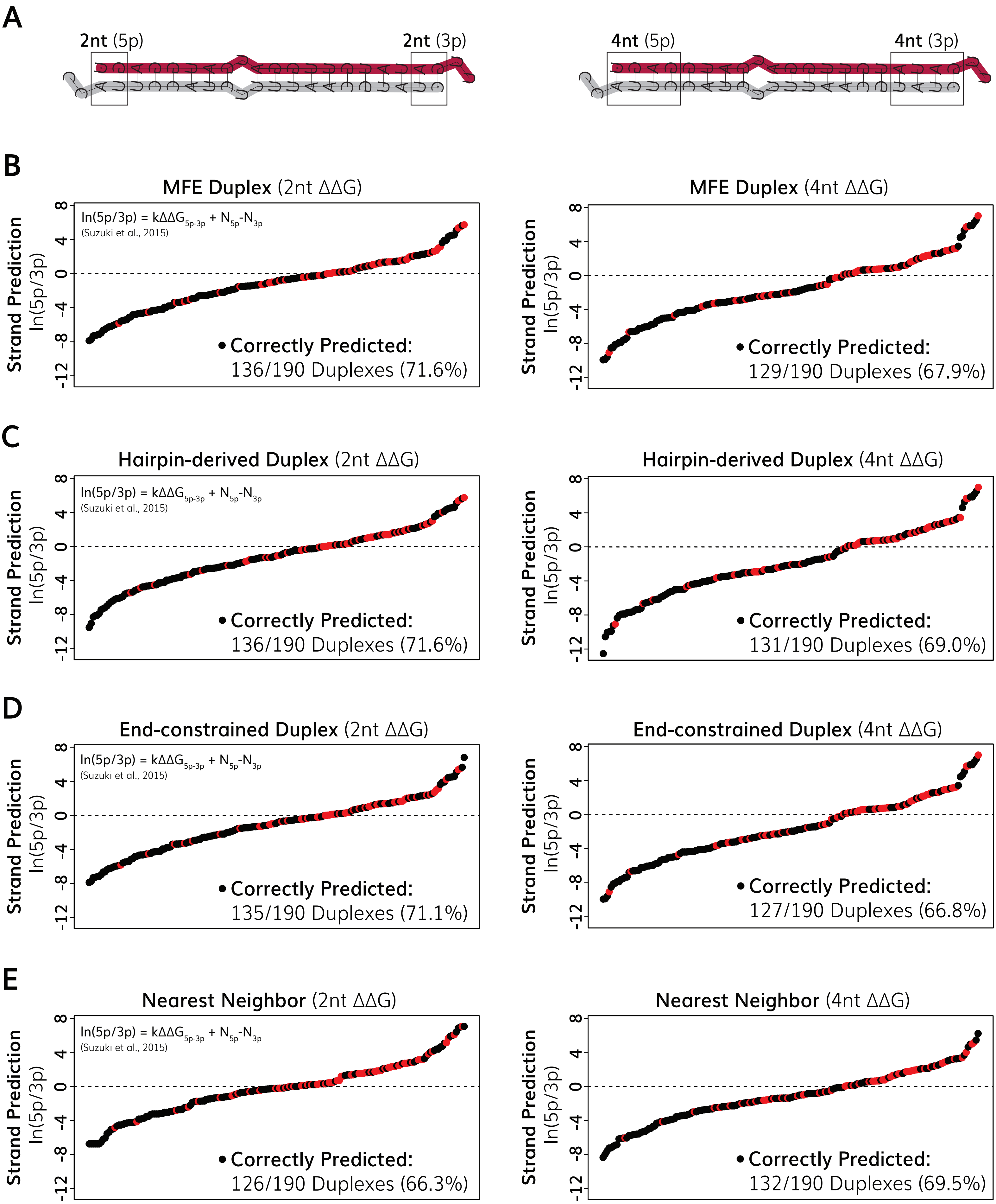
A
2nt (5p)
2nt (3p)
4nt (5p)
4nt (3p)
B
MFE Duplex (2nt ΔΔG)
ln(5p/3p) = kΔΔG5p-3p + N5p-N3p
(Suzuki et al., 2015)
Strand Prediction
ln(5p/3p)
• Correctly Predicted:
136/190 Duplexes (71.6%)
MFE Duplex (4nt ΔΔG)
• Correctly Predicted:
129/190 Duplexes (67.9%)
C
Hairpin-derived Duplex (2nt ΔΔG)
• Correctly Predicted:
136/190 Duplexes (71.6%)
Hairpin-derived Duplex (4nt ΔΔG)
• Correctly Predicted:
131/190 Duplexes (69.0%)
D
End-constrained Duplex (2nt ΔΔG)
• Correctly Predicted:
135/190 Duplexes (71.1%)
End-constrained Duplex (4nt ΔΔG)
• Correctly Predicted:
127/190 Duplexes (66.8%)
E
Nearest Neighbor (2nt ΔΔG)
• Correctly Predicted:
126/190 Duplexes (66.3%)
Nearest Neighbor (4nt ΔΔG)
• Correctly Predicted:
132/190 Duplexes (69.5%)

**Figure S3**

**Table S1:** *C. elegans* miRNA hairpins folded at different temperatures

| miRNA | Hairpin (15°C) | Hairpin (20°C) | Hairpin (25°C) | Hairpin (37°C) |
|---|---|---|---|---|
| *let-7* | 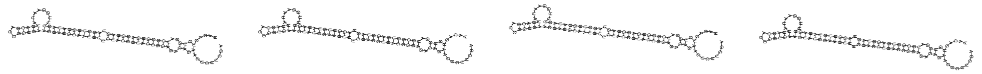 | | | |
| *lin-4* | 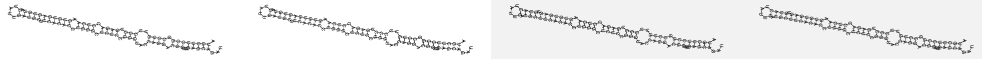 | | | |
| *lsy-6* | 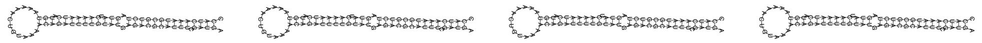 | | | |
| *mir-1018* | 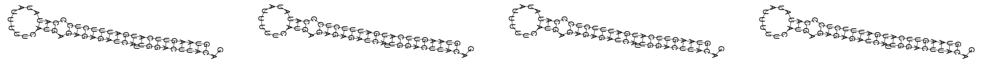 | | | |
| *mir-1019* | 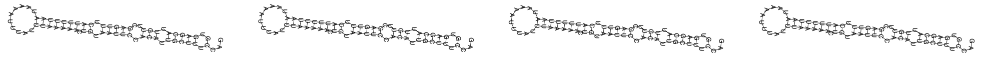 | | | |
| *mir-1020* | 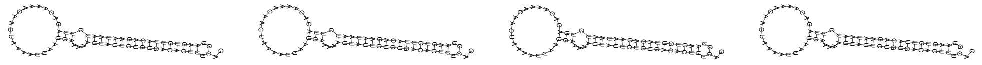 | | | |
| *mir-1021* | 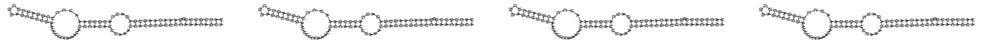 | | | |
| *mir-1022* | 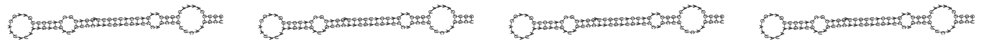 | | | |
| *mir-12134* | 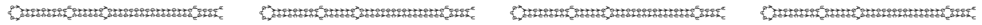 | | | |
| *mir-124* | 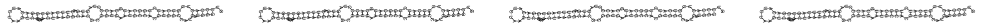 | | | |
| *mir-1817* | 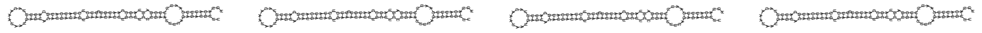 | | | |
| *mir-1818* | 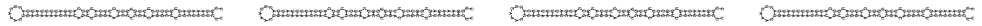 | | | |
| *mir-1819* | 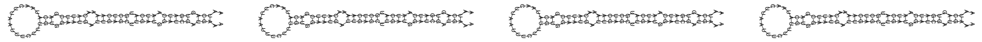 | | | |
| *mir-1820* | 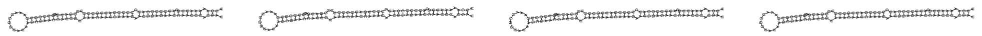 | | | |
| *mir-1821* | 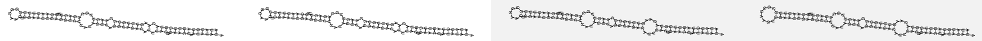 | | | |
| *mir-1822* | 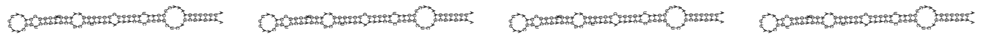 | | | |
| *mir-1823* | 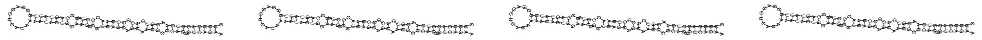 | | | |
| *mir-1824* | 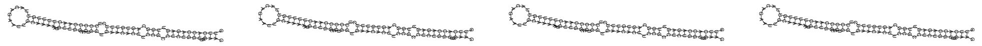 | | | |

| | |
|---|---|
| *mir-1828* | 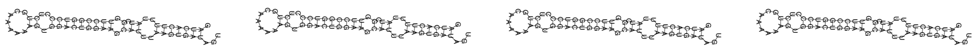 |
| *mir-1829a* | 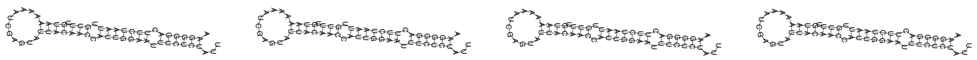 |
| *mir-1829b* | 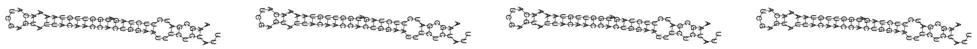 |
| *mir-1829c* | 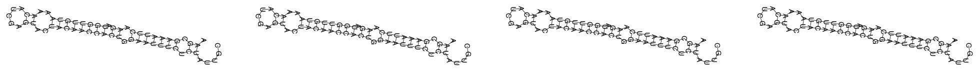 |
| *mir-1830* | 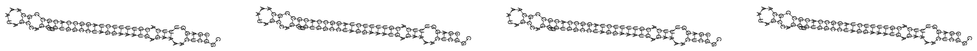 |
| *mir-1832a* | 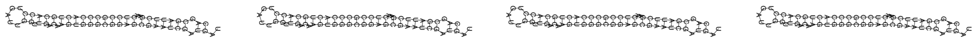 |
| *mir-1832b* | 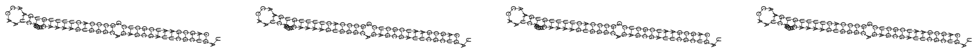 |
| *mir-1833* | 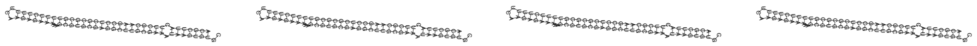 |
| *mir-1* | 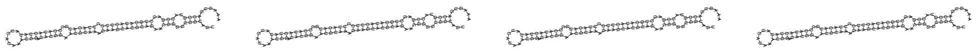 |
| *mir-2207* | 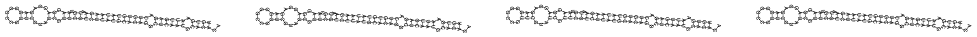 |
| *mir-2208a* | 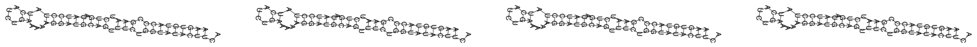 |
| *mir-2208b* | 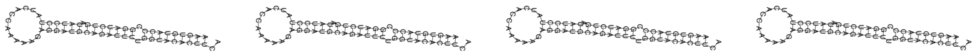 |
| *mir-2209a* | 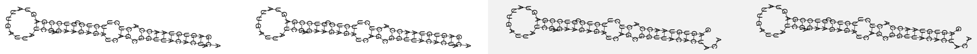 |
| *mir-2209b* | 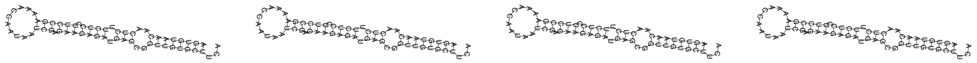 |
| *mir-2209c* | 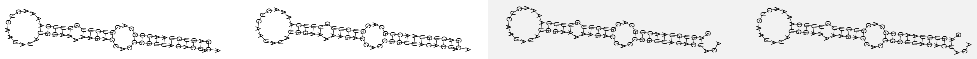 |
| *mir-2210* | 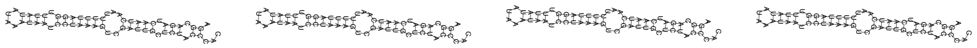 |
| *mir-2211* | 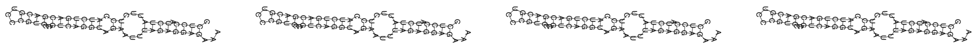 |
| *mir-2212* | 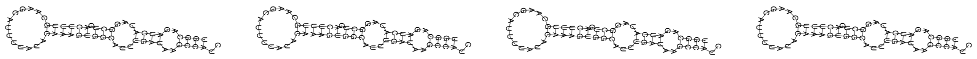 |
| *mir-2213* | 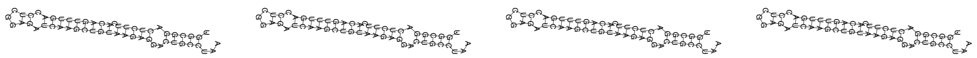 |

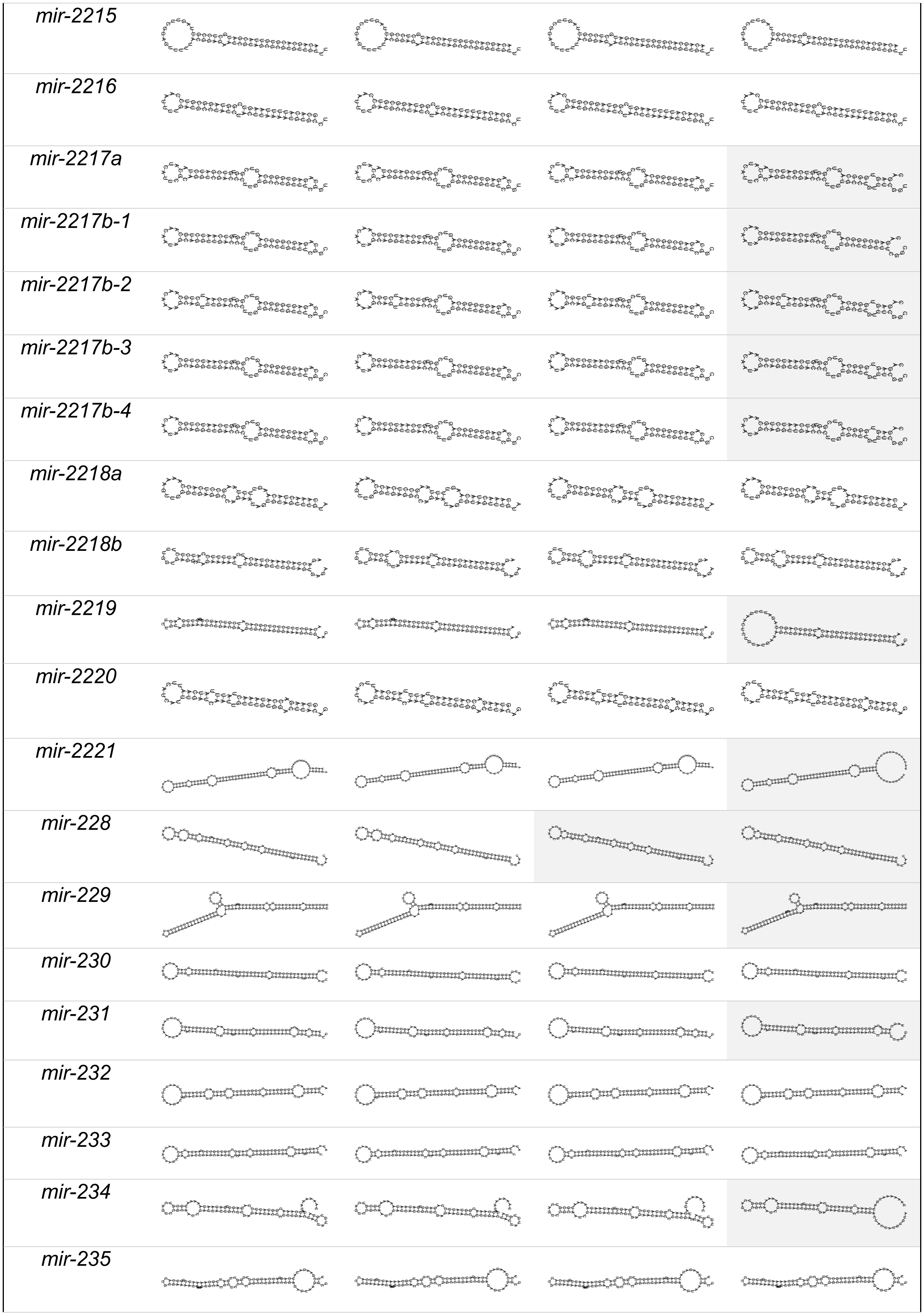
mir-2215
mir-2216
mir-2217a
mir-2217b-1
mir-2217b-2
mir-2217b-3
mir-2217b-4
mir-2218a
mir-2218b
mir-2219
mir-2220
mir-2221
mir-228
mir-229
mir-230
mir-231
mir-232
mir-233
mir-234
mir-235

| | |
|---|---|
| *mir-236* | 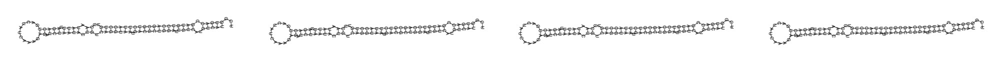 |
| *mir-237* | 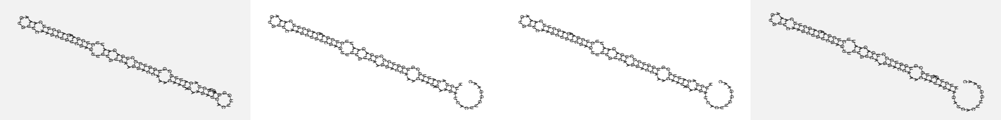 |
| *mir-238* | 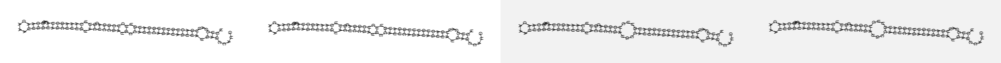 |
| *mir-239a* | 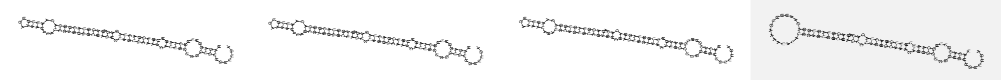 |
| *mir-239b* | 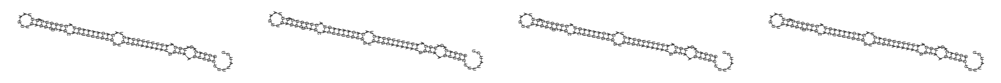 |
| *mir-240* | 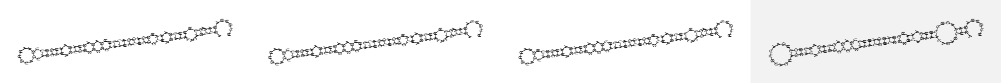 |
| *mir-241* | 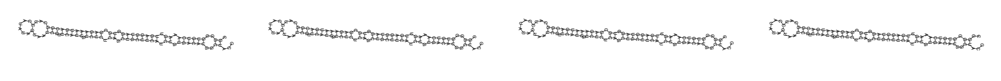 |
| *mir-242* | 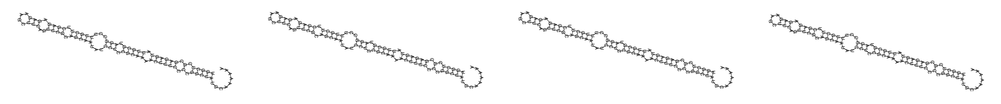 |
| *mir-243* | 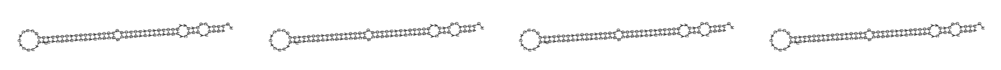 |
| *mir-244* | 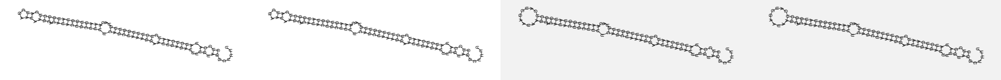 |
| *mir-245* | 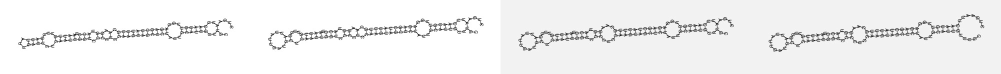 |
| *mir-246* | 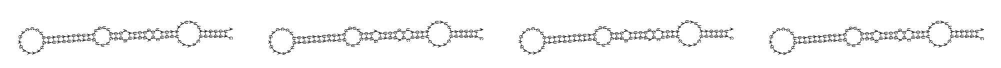 |
| *mir-247* | 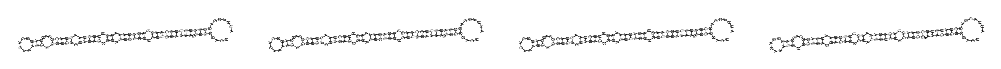 |
| *mir-248* | 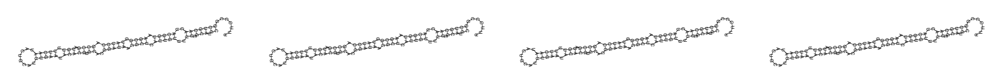 |
| *mir-249* | 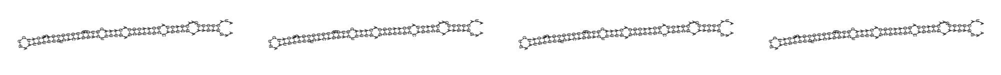 |
| *mir-250* | 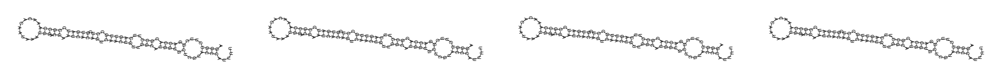 |
| *mir-251* | 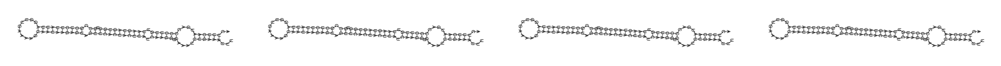 |
| *mir-252* | 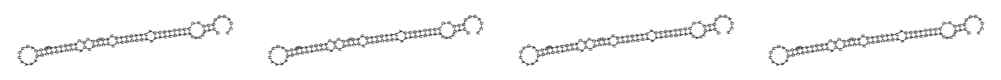 |
| *mir-253* | 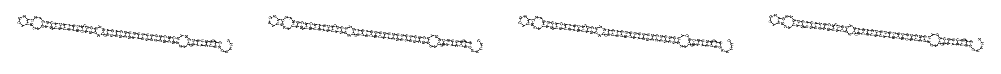 |
| *mir-254* | 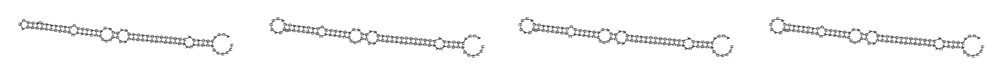 |

| | |
|---|---|
| *mir-255* | 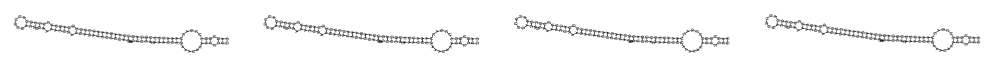 |
| *mir-256* | 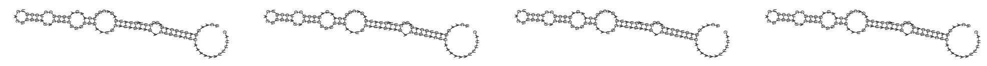 |
| *mir-259* | 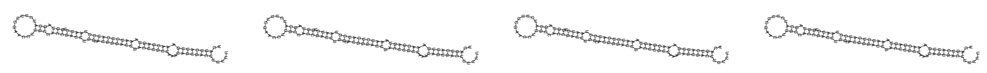 |
| *mir-261* | 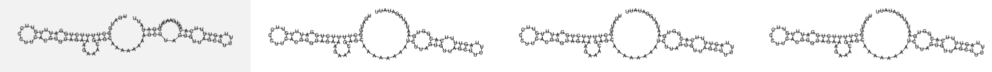 |
| *mir-264* | 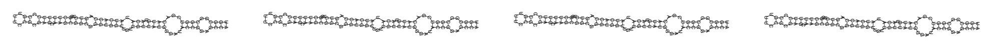 |
| *mir-265* | 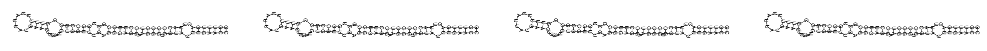 |
| *mir-266* | 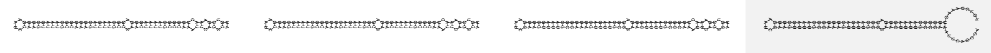 |
| *mir-267* | 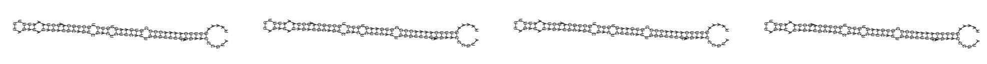 |
| *mir-268* | 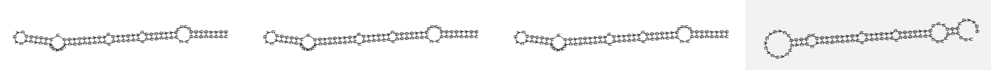 |
| *mir-269* | 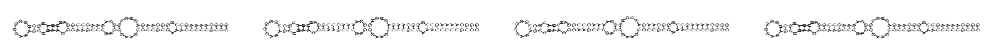 |
| *mir-270* | 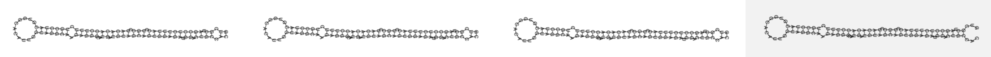 |
| *mir-271* | 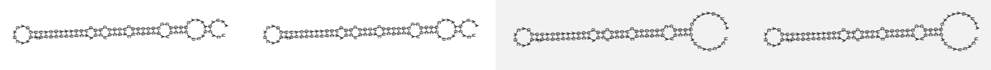 |
| *mir-272* | 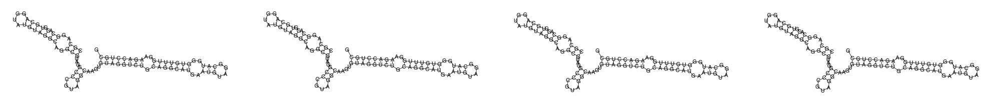 |
| *mir-273* | 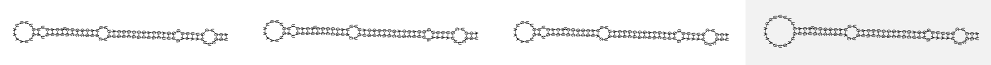 |
| *mir-2953* | 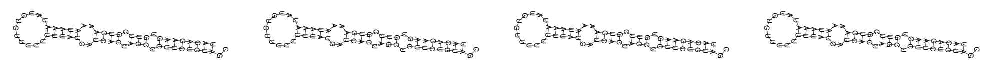 |
| *mir-2* | 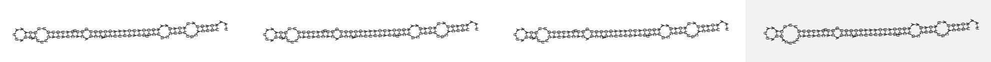 |
| *mir-34* | 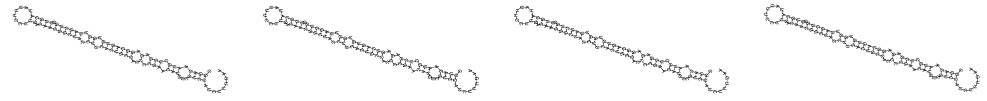 |
| *mir-354* | 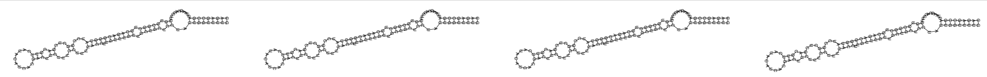 |
| *mir-355* | 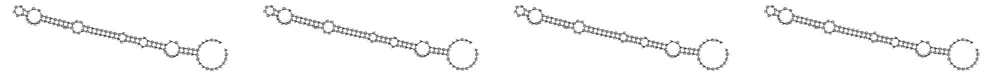 |
| *mir-356a* | 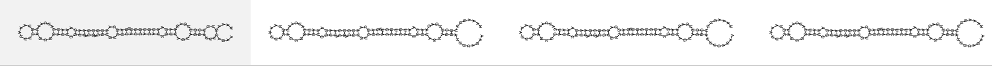 |
| *mir-356b* | 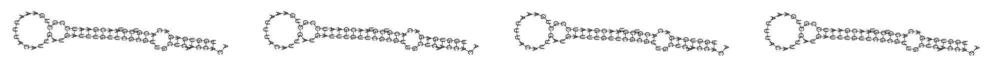 |
| *mir-357* | 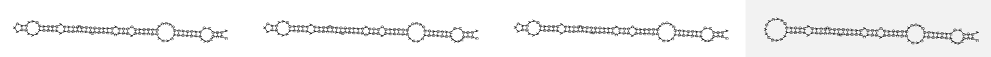 |

| | |
|---|---|
| *mir-358* | 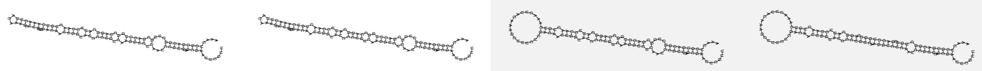 |
| *mir-359* | 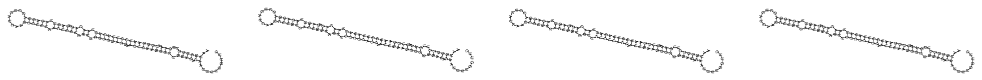 |
| *mir-35* | 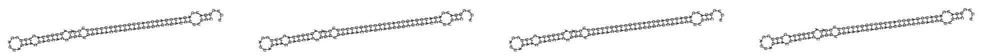 |
| *mir-360* | 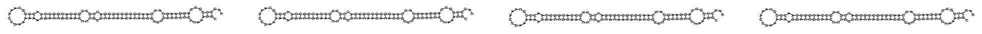 |
| *mir-36* | 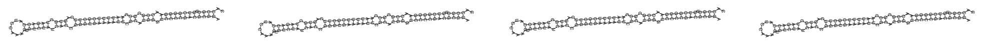 |
| *mir-37* | 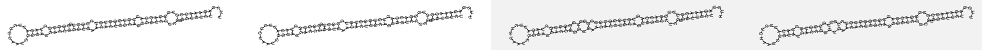 |
| *mir-38* | 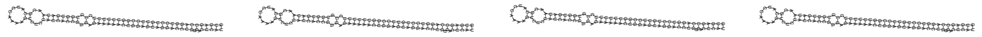 |
| *mir-392* | 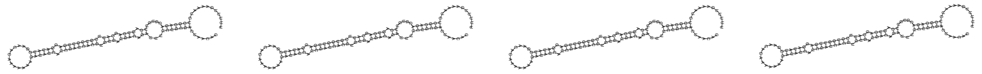 |
| *mir-39* | 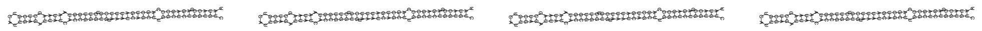 |
| *mir-40* | 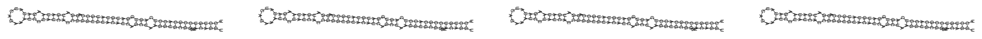 |
| *mir-41* | 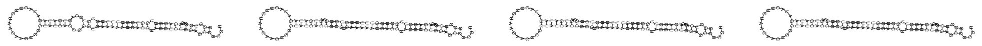 |
| *mir-42* | 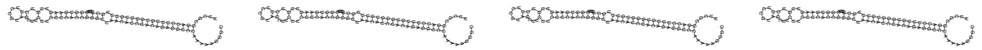 |
| *mir-43* | 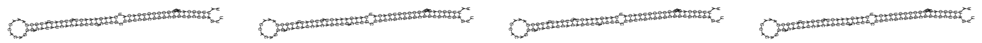 |
| *mir-44* | 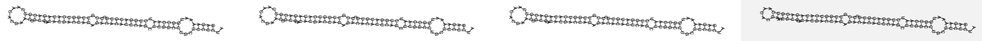 |
| *mir-45* | 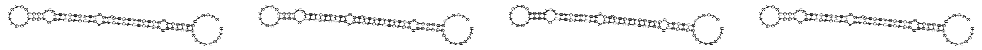 |
| *mir-46* | 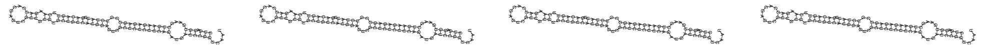 |
| *mir-47* | 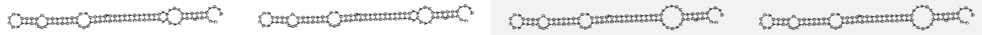 |
| *mir-4805* | 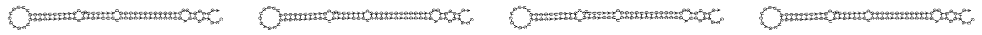 |
| *mir-4806* | 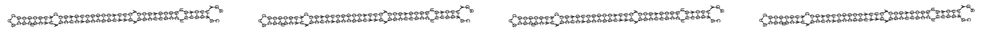 |
| *mir-4807* | 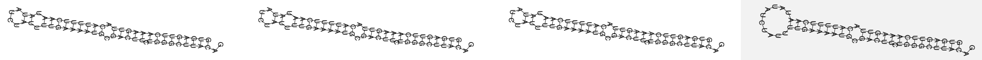 |
| *mir-4808* | 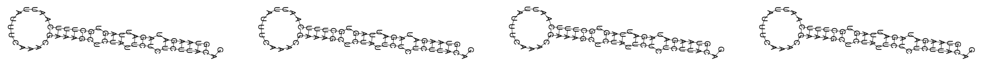 |
| *mir-4809* | 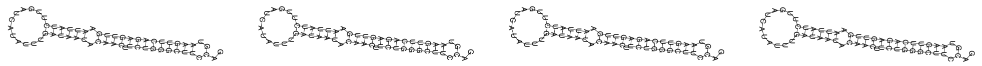 |
| *mir-4810a* | 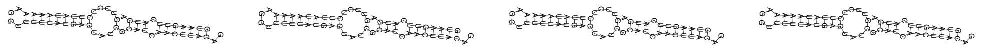 |
| *mir-4810b* | 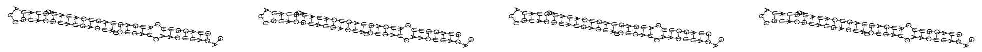 |

| | |
|---|---|
| *mir-4811* | 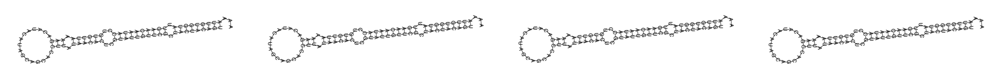 |
| *mir-4812* | 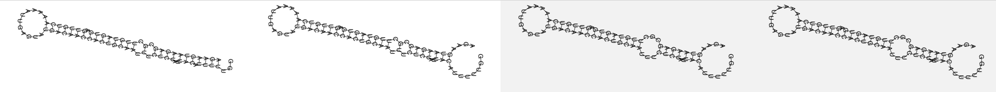 |
| *mir-4813* | 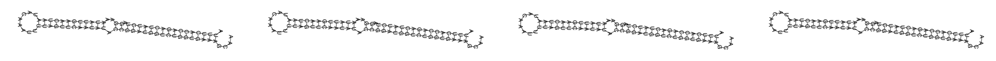 |
| *mir-4814* | 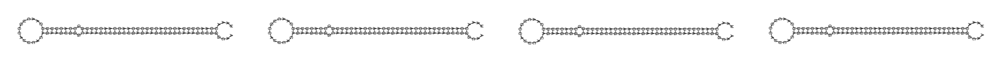 |
| *mir-4815* | 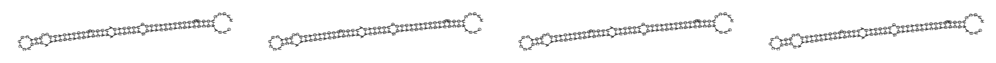 |
| *mir-4816* | 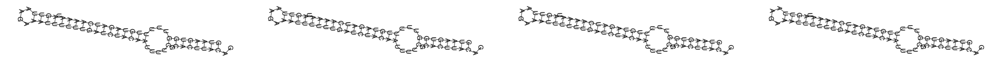 |
| *mir-48* | 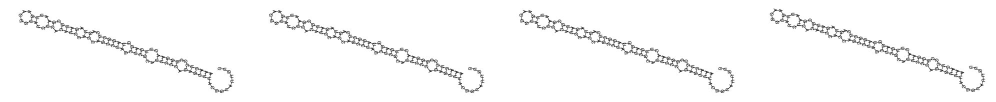 |
| *mir-4920* | 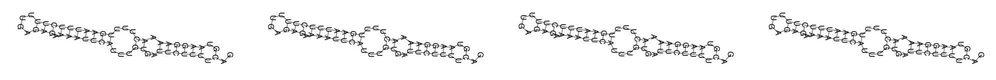 |
| *mir-4921* | 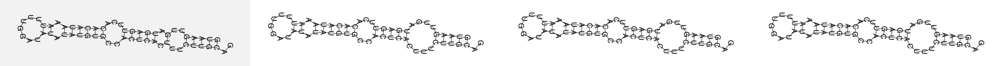 |
| *mir-4922-1* | 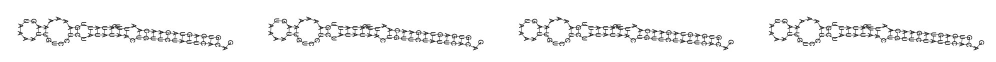 |
| *mir-4922-2* | 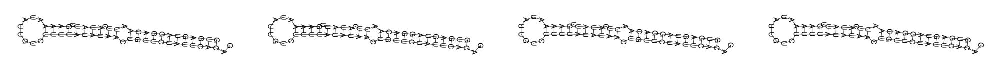 |
| *mir-4923a* | 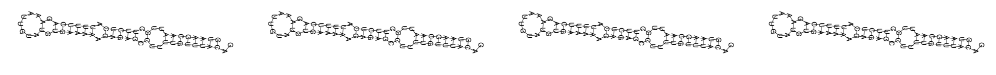 |
| *mir-4923b* | 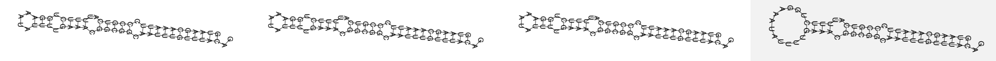 |
| *mir-4924* | 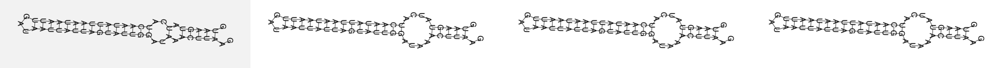 |
| *mir-4925* | 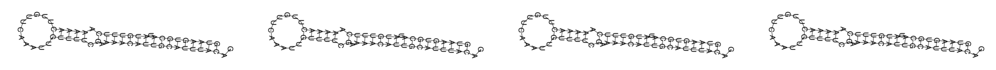 |
| *mir-4926* | 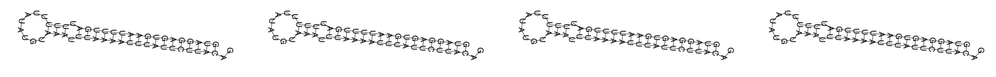 |
| *mir-4927* | 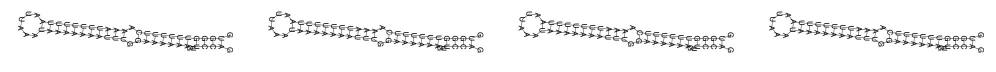 |
| *mir-4929* | 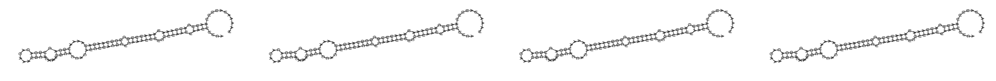 |
| *mir-4930* | 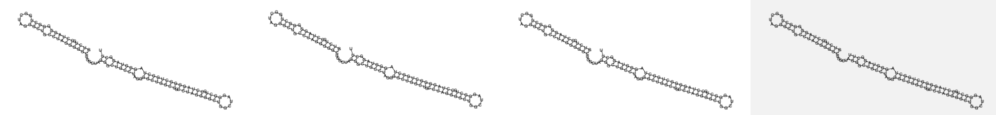 |
| *mir-4931* | 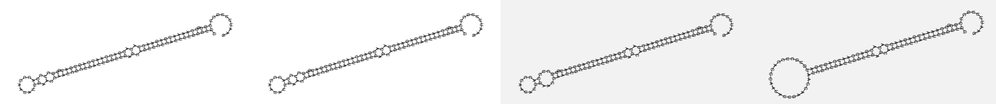 |
| *mir-4932* | 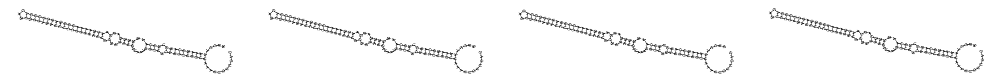 |

| | |
|---|---|
| *mir-4933* | 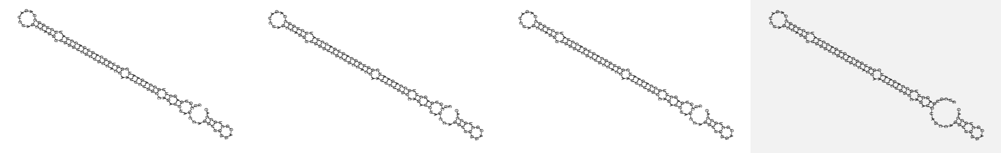 |
| *mir-4934* | 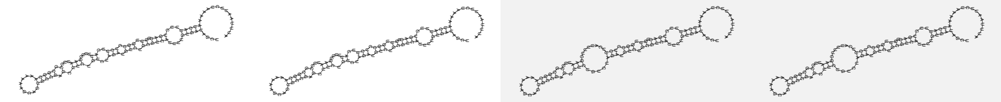 |
| *mir-4935* | 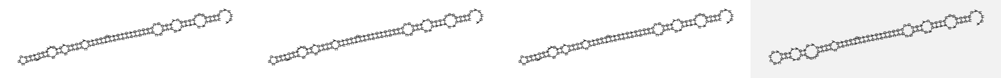 |
| *mir-4936* | 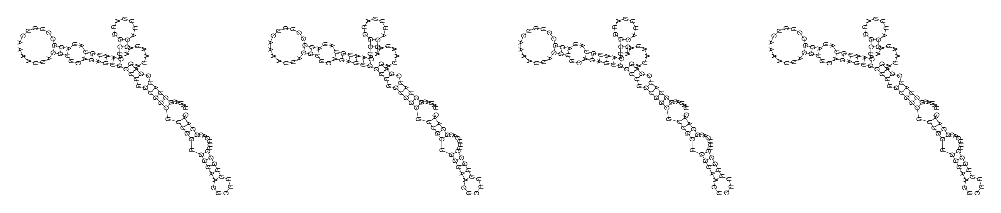 |
| *mir-4937* | 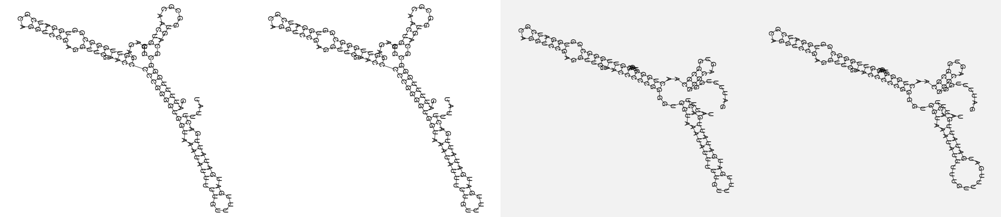 |
| *mir-4938* | 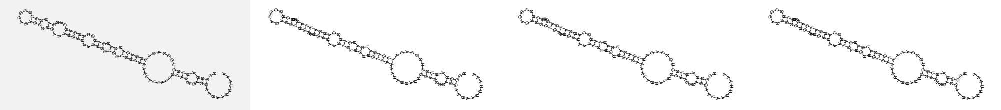 |
| *mir-49* | 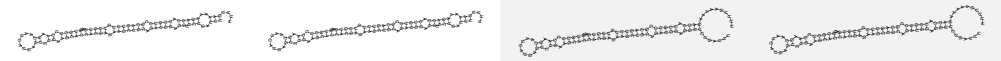 |
| *mir-50* | 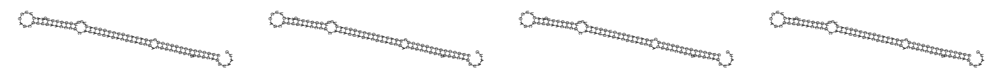 |
| *mir-51* | 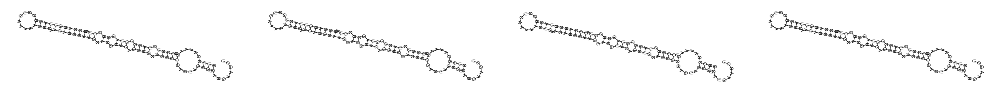 |
| *mir-52* | 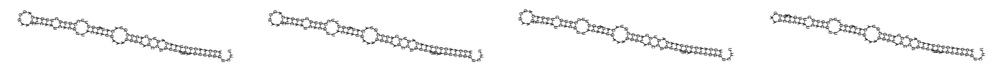 |
| *mir-53* | 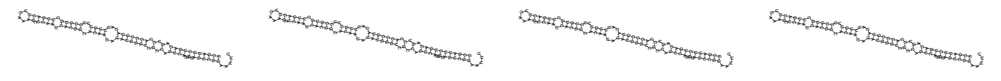 |
| *mir-54* | 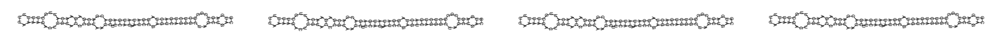 |
| *mir-5545* | 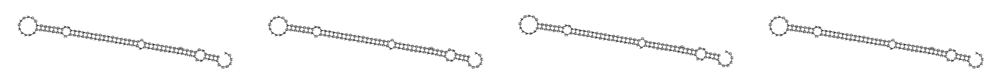 |
| *mir-5546* | 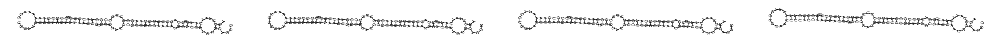 |
| *mir-5547* | 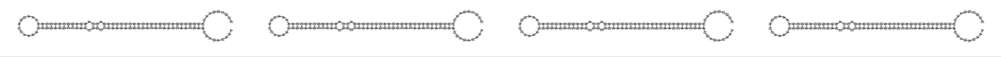 |

| | |
|---|---|
| *mir-5548* | 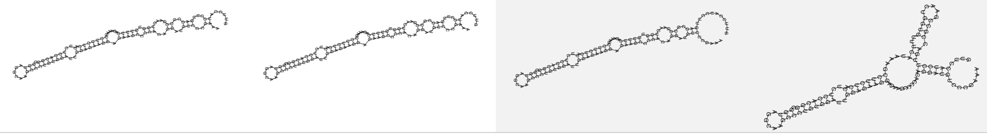 |
| *mir-5549* | 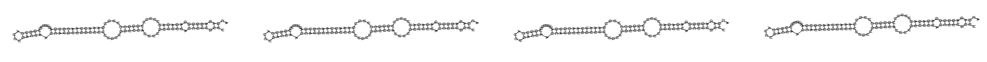 |
| *mir-5550* | 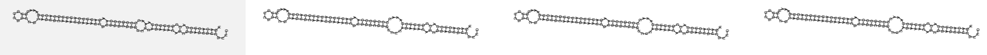 |
| *mir-5551* | 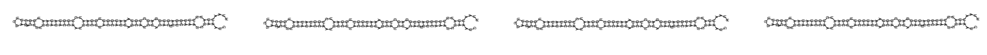 |
| *mir-5552* | 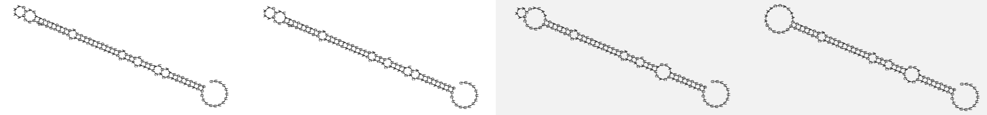 |
| *mir-5553* | 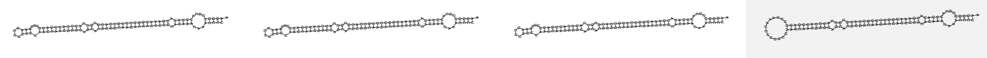 |
| *mir-5592-1* | 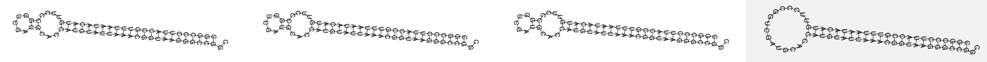 |
| *mir-5592-2* | 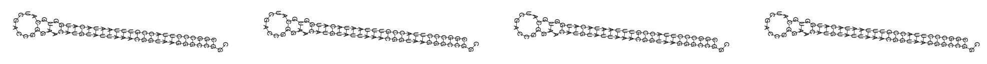 |
| *mir-5593-1* | 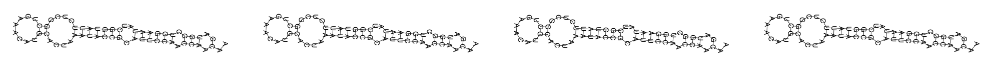 |
| *mir-5593-2* | 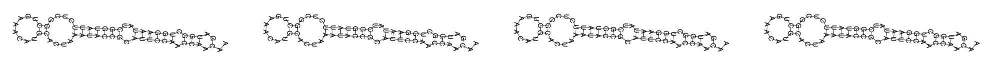 |
| *mir-5594* | 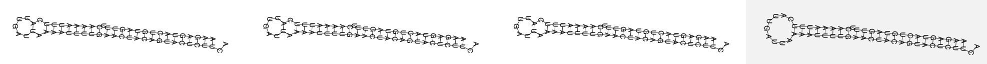 |
| *mir-5595* | 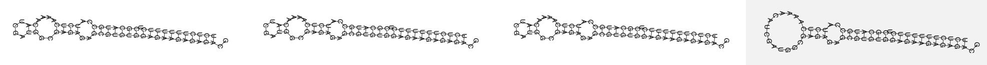 |
| *mir-55* | 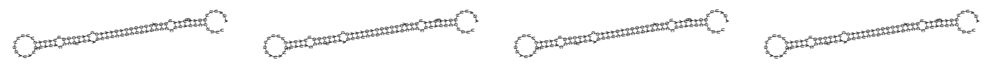 |
| *mir-55b* | 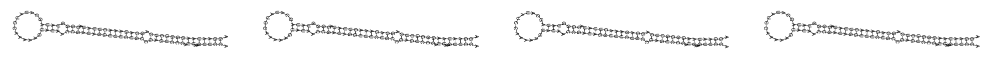 |
| *mir-56* | 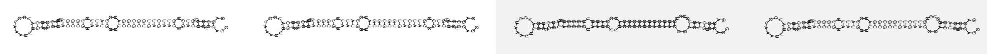 |
| *mir-57* | 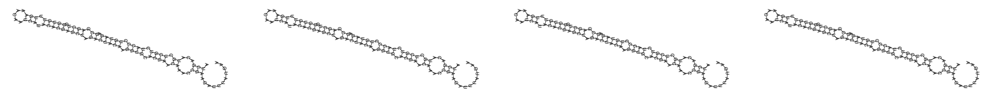 |
| *mir-58a* | 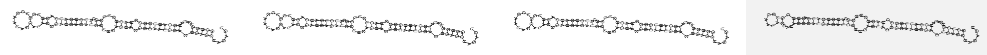 |
| *mir-58b* | 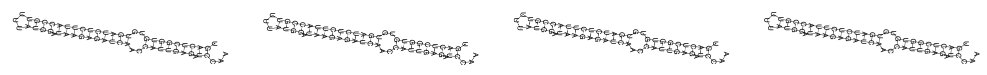 |
| *mir-58c* | 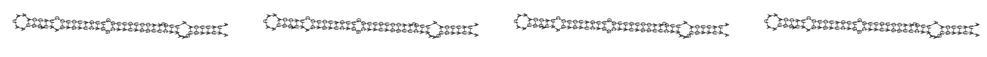 |
| *mir-59* | 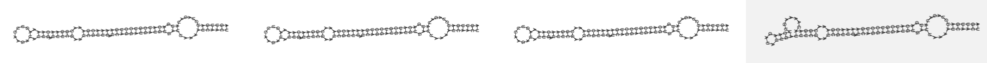 |
| *mir-60* | 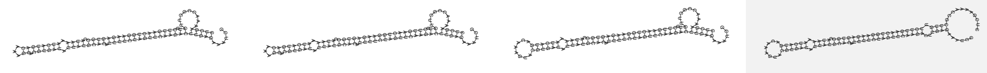 |
| *mir-61* | 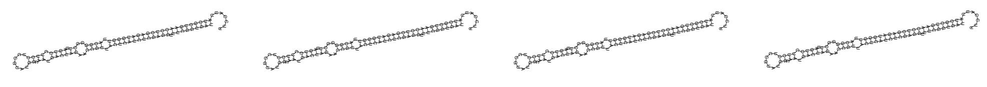 |

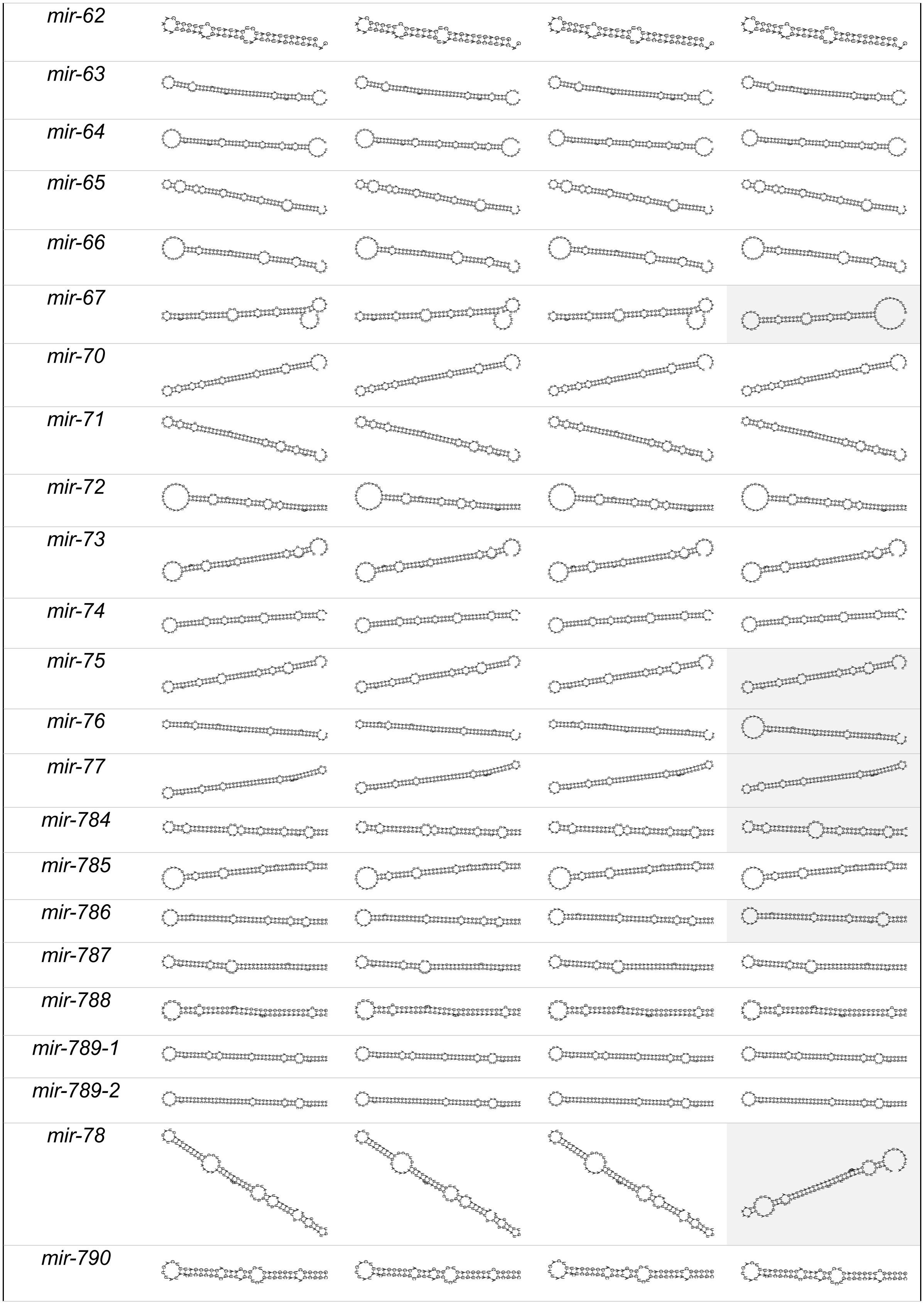
mir-62
mir-63
mir-64
mir-65
mir-66
mir-67
mir-70
mir-71
mir-72
mir-73
mir-74
mir-75
mir-76
mir-77
mir-784
mir-785
mir-786
mir-787
mir-788
mir-789-1
mir-789-2
mir-78
mir-790

| | |
|---|---|
| *mir-791* | 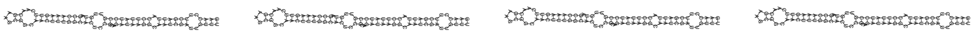 |
| *mir-792* | 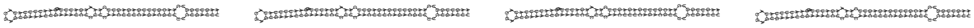 |
| *mir-793* | 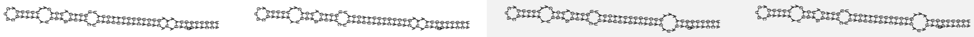 |
| *mir-794* | 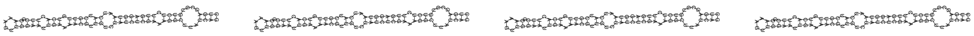 |
| *mir-795* | 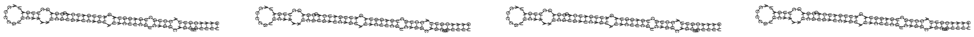 |
| *mir-796* | 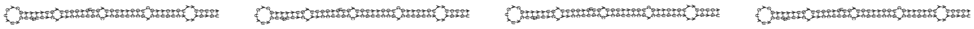 |
| *mir-797* | 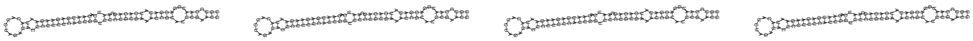 |
| *mir-798* | 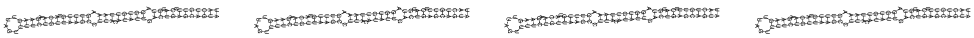 |
| *mir-799* | 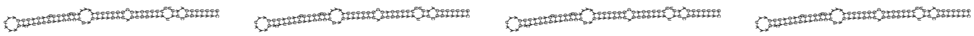 |
| *mir-79* | 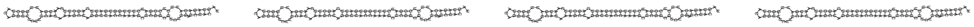 |
| *mir-800* | 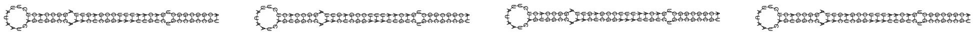 |
| *mir-80* | 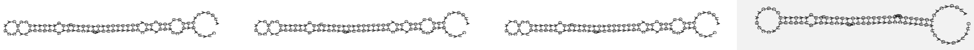 |
| *mir-8186-1* | 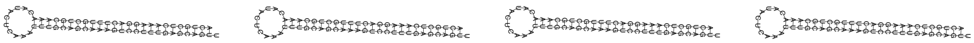 |
| *mir-8186-2* | 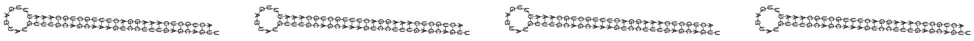 |
| *mir-8187* | 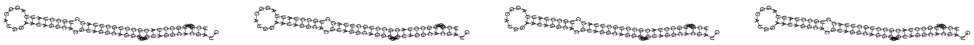 |
| *mir-8188* | 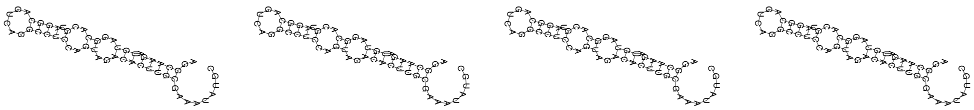 |
| *mir-8189* | 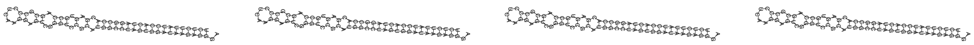 |
| *mir-8190* | 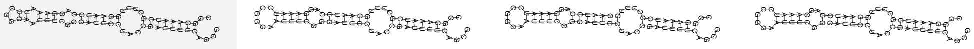 |
| *mir-8191* | 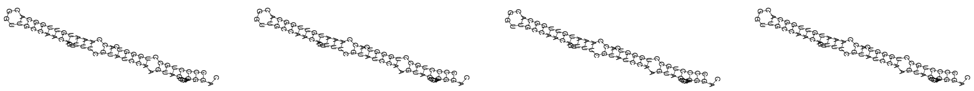 |
| *mir-8192* | 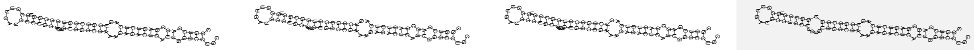 |
| *mir-8193* | 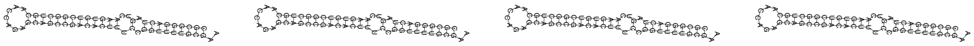 |
| *mir-8194* | 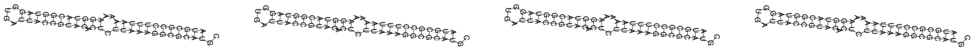 |
| *mir-8195* | 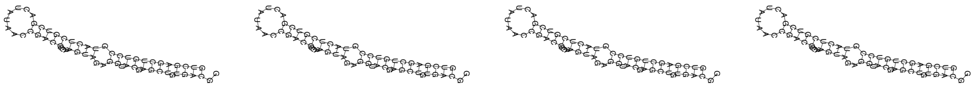 |

| | |
|---|---|
| *mir-8196a* | 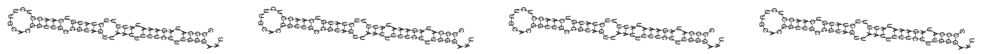 |
| *mir-8196b* | 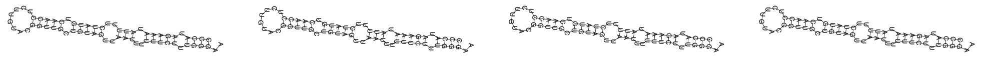 |
| *mir-8197* | 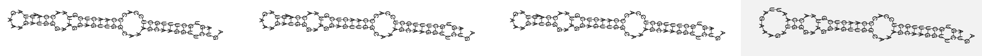 |
| *mir-8198* | 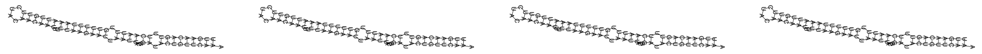 |
| *mir-8199* | 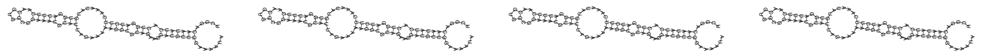 |
| *mir-81* | 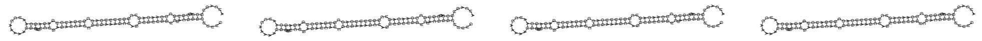 |
| *mir-8200* | 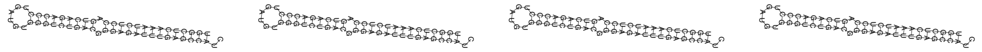 |
| *mir-8201* | 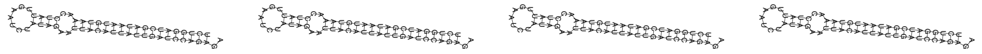 |
| *mir-8202* | 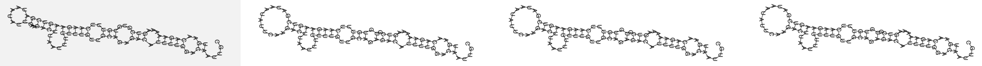 |
| *mir-8203* | 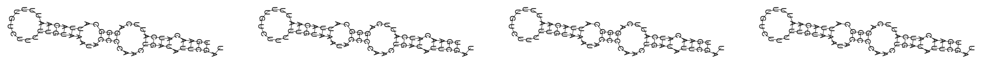 |
| *mir-8204* | 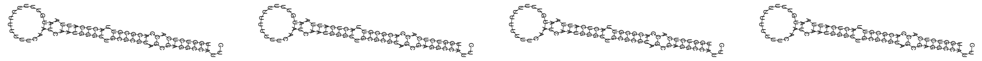 |
| *mir-8205* | 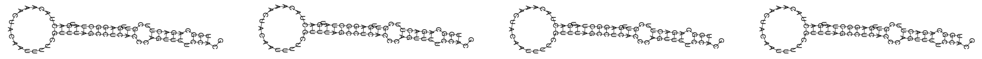 |
| *mir-8206* | 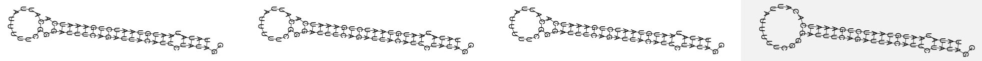 |
| *mir-8207* | 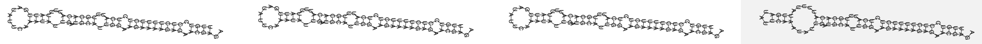 |
| *mir-8208* | 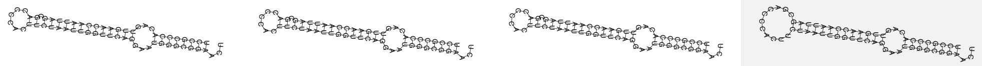 |
| *mir-8209* | 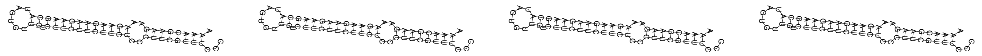 |
| *mir-8210* | 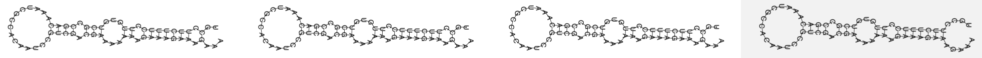 |
| *mir-8211* | 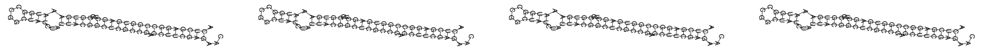 |
| *mir-8212* | 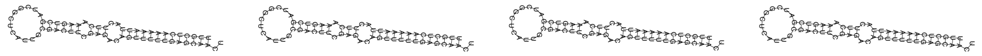 |
| *mir-82* | 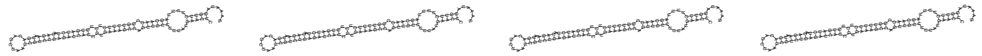 |
| *mir-83* | 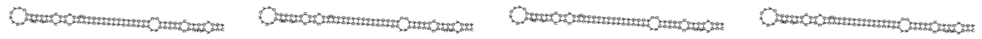 |
| *mir-84* | 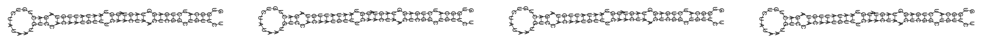 |
| *mir-85* | 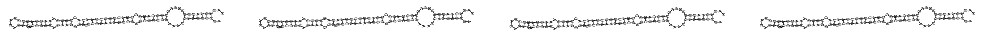 |

| | | | | |
|---|---|---|---|---|
| *mir-86* | 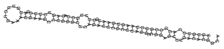 | 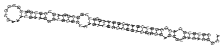 | 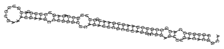 | 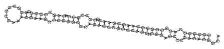 |
| *mir-87* | 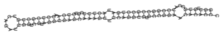 | 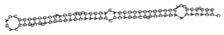 | 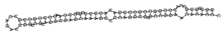 | 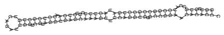 |
| *mir-90* | 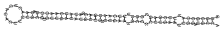 | 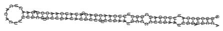 | 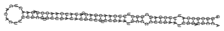 | 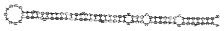 |

**Table S2:** *C. elegans* miRNA duplexes folded at different temperatures

| miRNA | Duplex (15°C) | Duplex (20°C) | Duplex (25°C) | Duplex (37°C) |
|---|---|---|---|---|
| *let-7* | 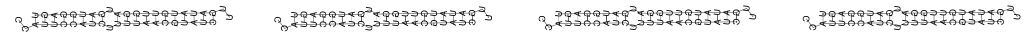 | | | |
| *lin-4* | 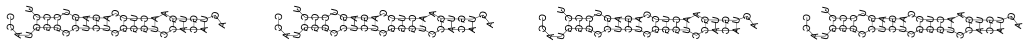 | | | |
| *lsy-6* | 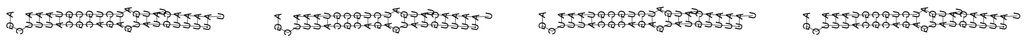 | | | |
| *mir-1019* | 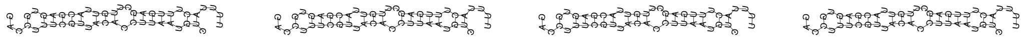 | | | |
| *mir-1020* | 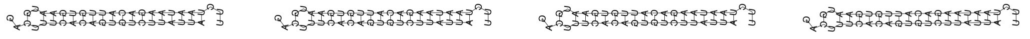 | | | |
| *mir-1022* | 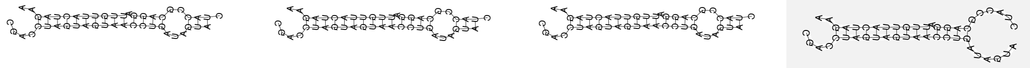 | | | |
| *mir-124* | 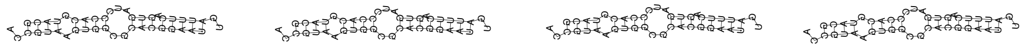 | | | |
| *mir-1819* | 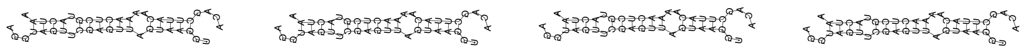 | | | |
| *mir-1820* | 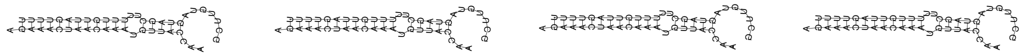 | | | |
| *mir-1821* | 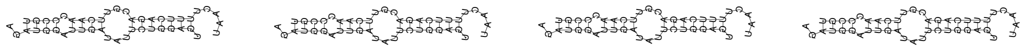 | | | |
| *mir-1822* | 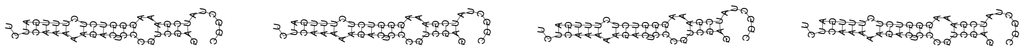 | | | |
| *mir-1823* | 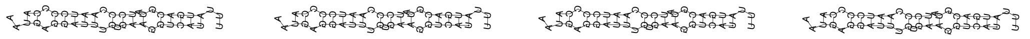 | | | |
| *mir-1824* | 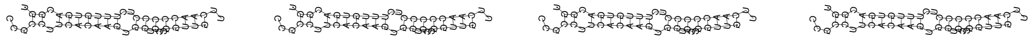 | | | |
| *mir-1829a* | 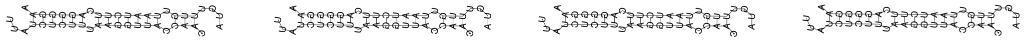 | | | |
| *mir-1829b* | 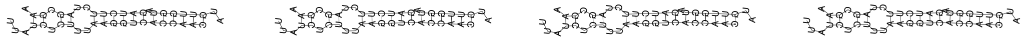 | | | |
| *mir-1829c* | 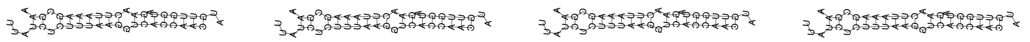 | | | |
| *mir-1830* | 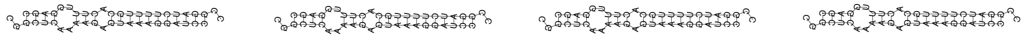 | | | |
| *mir-1832a* | 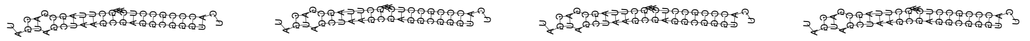 | | | |
| *mir-1832b* | 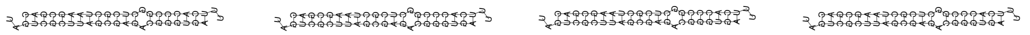 | | | |
| *mir-1* | 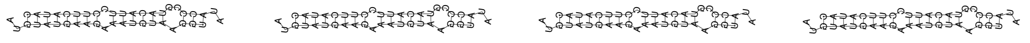 | | | |
| *mir-2207* | 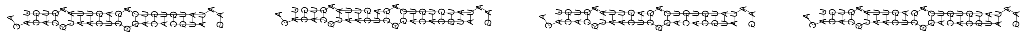 | | | |
| *mir-2208a* | 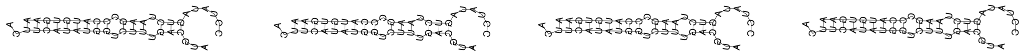 | | | |
| *mir-2208b* | 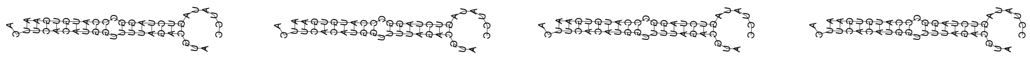 | | | |

| | |
|---|---|
| *mir-2209a* | 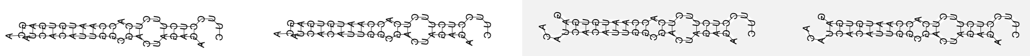 |
| *mir-2209b* | 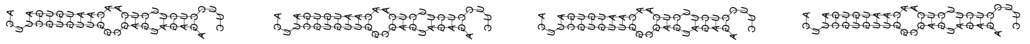 |
| *mir-2209c* | 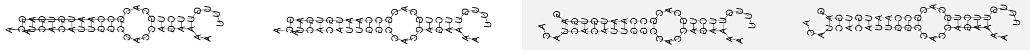 |
| *mir-2210* | 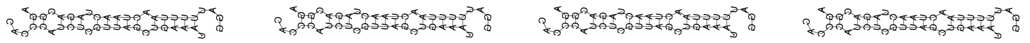 |
| *mir-2211* | 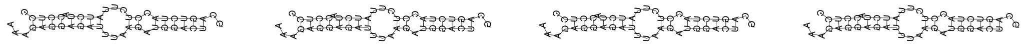 |
| *mir-2212* | 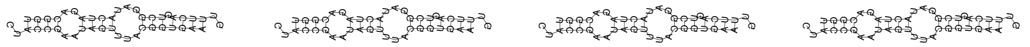 |
| *mir-2213* | 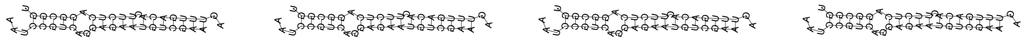 |
| *mir-2215* | 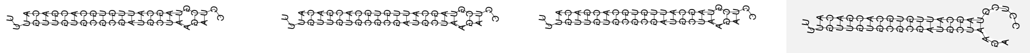 |
| *mir-2216* | 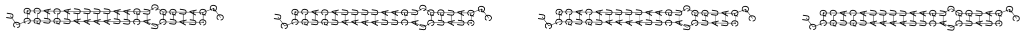 |
| *mir-2217a* | 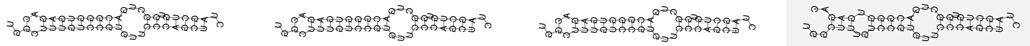 |
| *mir-2217b* | 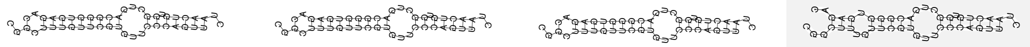 |
| *mir-2218a* | 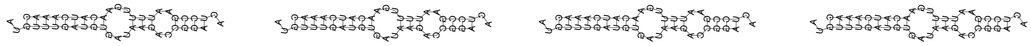 |
| *mir-2218b* | 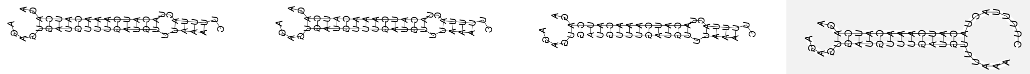 |
| *mir-2219* | 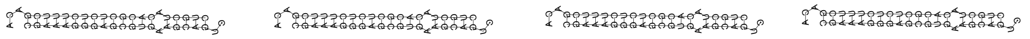 |
| *mir-2220* | 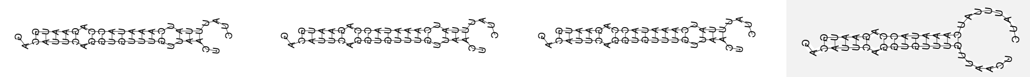 |
| *mir-228* | 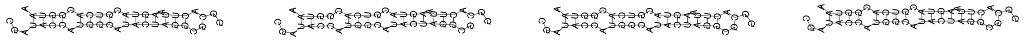 |
| *mir-229* | 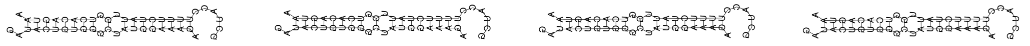 |
| *mir-230* | 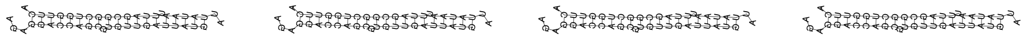 |
| *mir-231* | 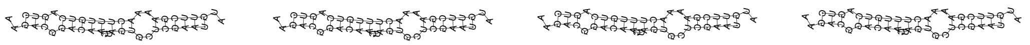 |
| *mir-232* | 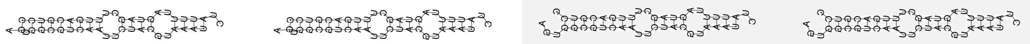 |
| *mir-233* | 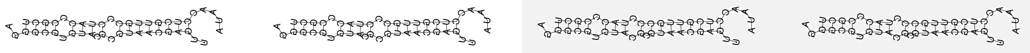 |
| *mir-234* | 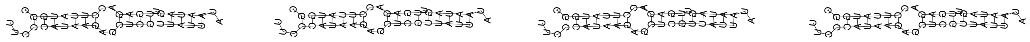 |
| *mir-235* | 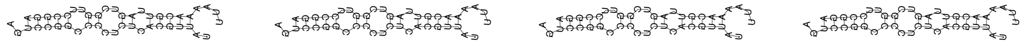 |
| *mir-236* | 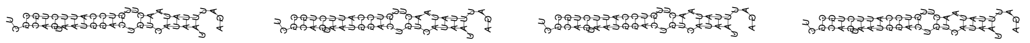 |
| *mir-237* | 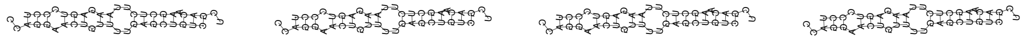 |

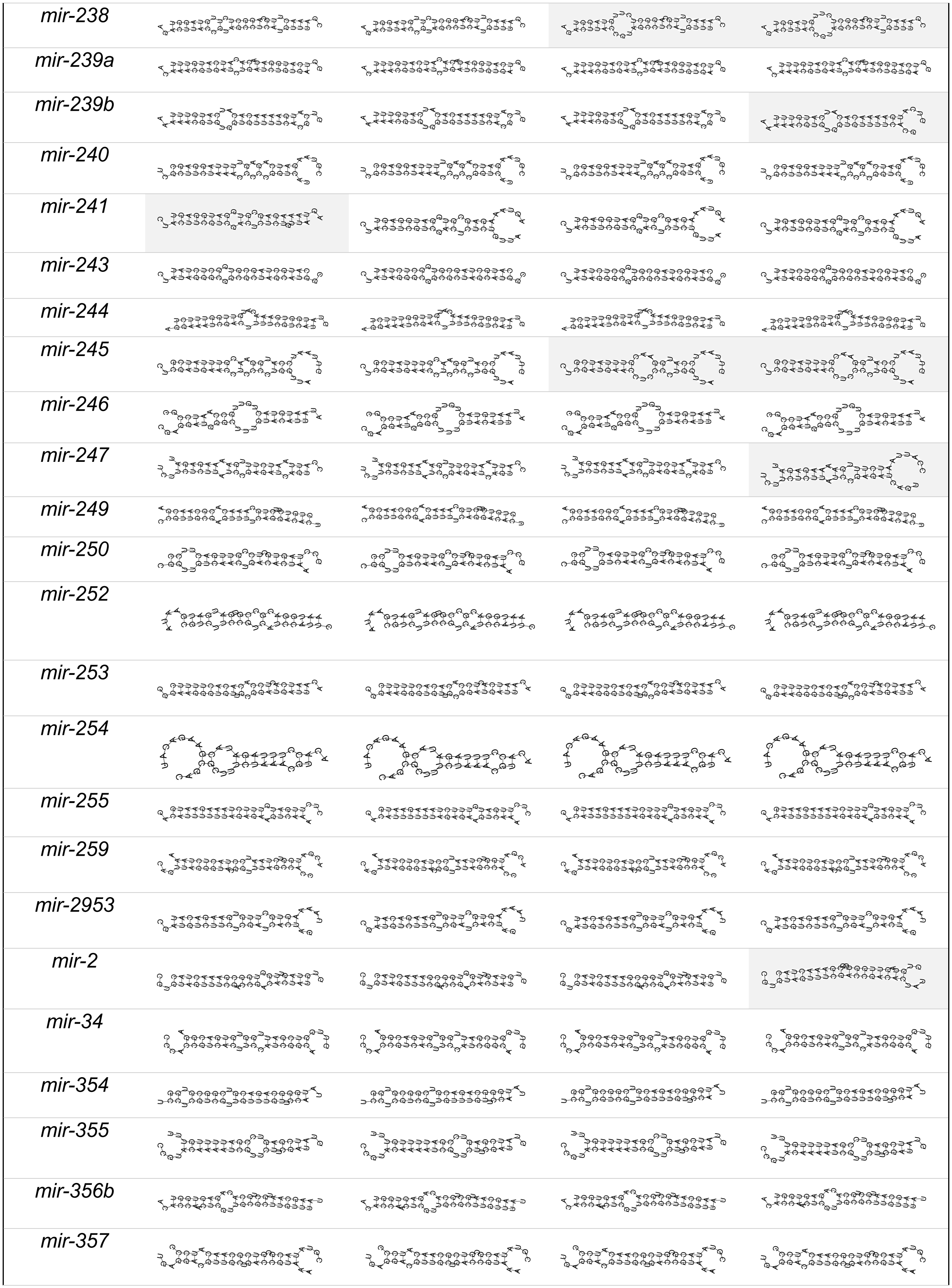
mir-238
mir-239a
mir-239b
mir-240
mir-241
mir-243
mir-244
mir-245
mir-246
mir-247
mir-249
mir-250
mir-252
mir-253
mir-254
mir-255
mir-259
mir-2953
mir-2
mir-34
mir-354
mir-355
mir-356b
mir-357

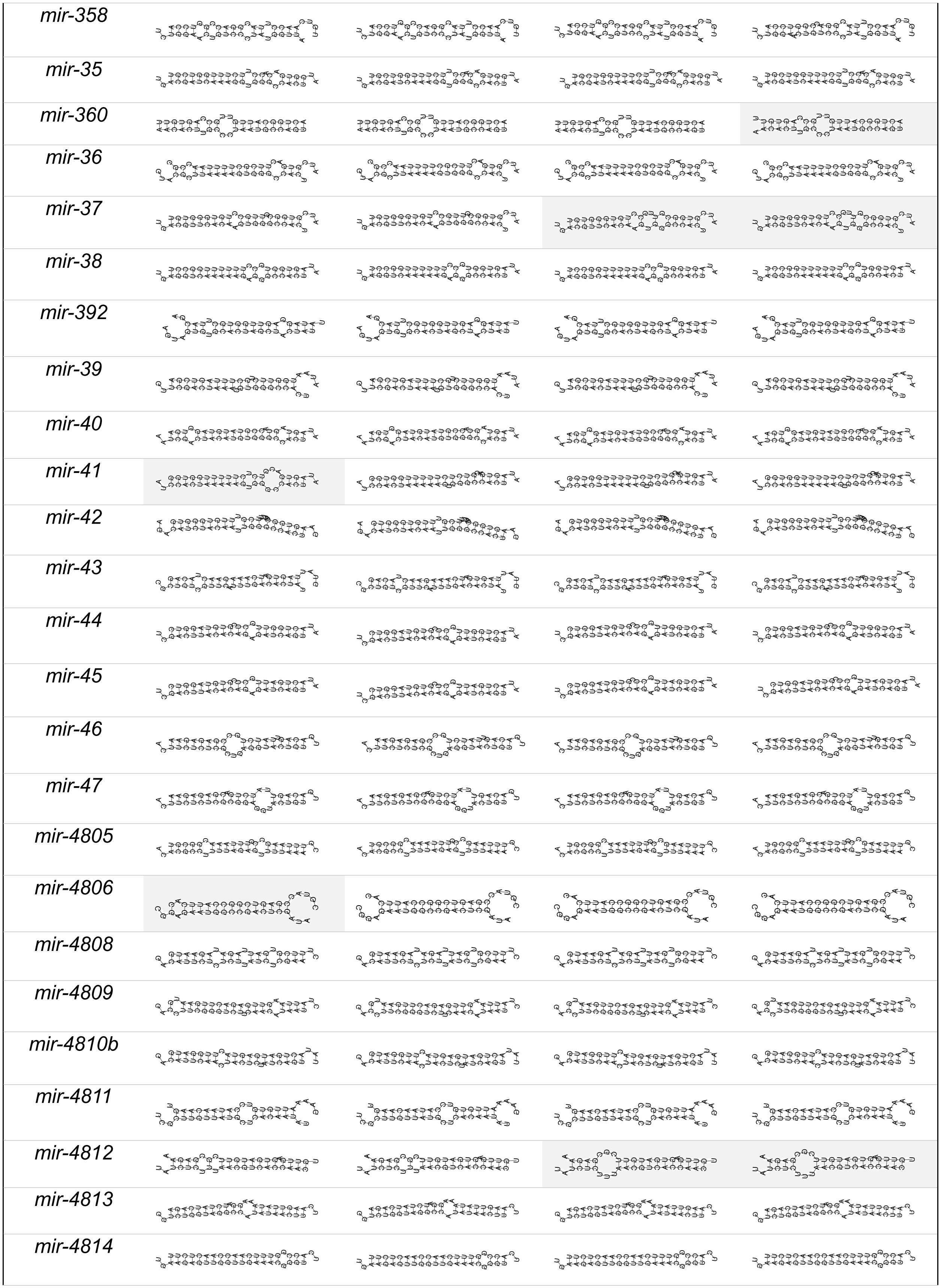

mir-358
mir-35
mir-360
mir-36
mir-37
mir-38
mir-392
mir-39
mir-40
mir-41
mir-42
mir-43
mir-44
mir-45
mir-46
mir-47
mir-4805
mir-4806
mir-4808
mir-4809
mir-4810b
mir-4811
mir-4812
mir-4813
mir-4814

| | |
|---|---|
| *mir-4816* | 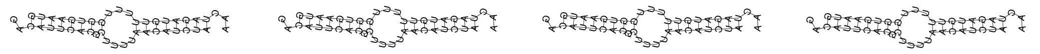 |
| *mir-48* | 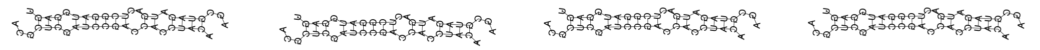 |
| *mir-49* | 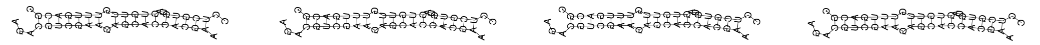 |
| *mir-50* | 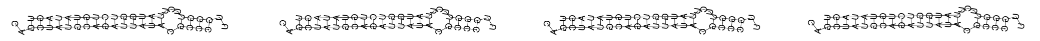 |
| *mir-51* | 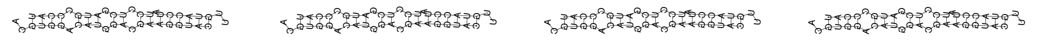 |
| *mir-52* | 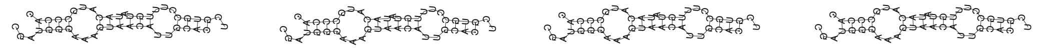 |
| *mir-53* | 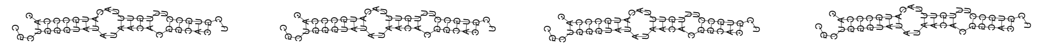 |
| *mir-54* | 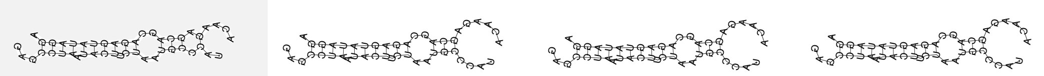 |
| *mir-5545* | 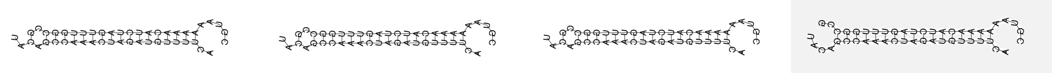 |
| *mir-5546* | 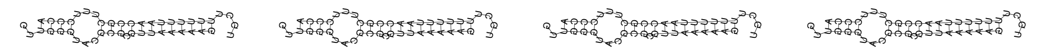 |
| *mir-5547* | 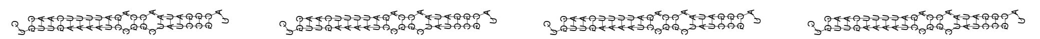 |
| *mir-5548* | 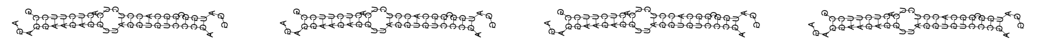 |
| *mir-5549* | 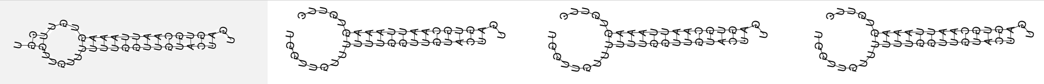 |
| *mir-5550* | 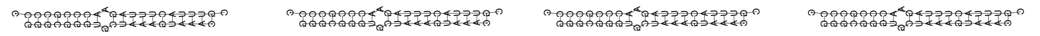 |
| *mir-5551* | 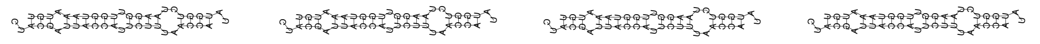 |
| *mir-5552* | 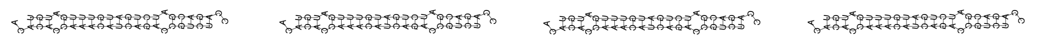 |
| *mir-5553* | 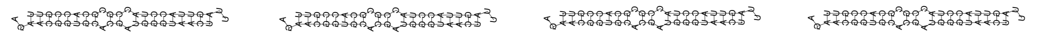 |
| *mir-5592* | 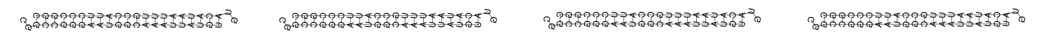 |
| *mir-5593* | 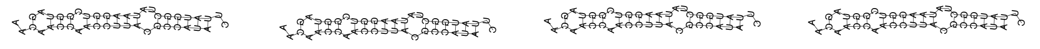 |
| *mir-5594* | 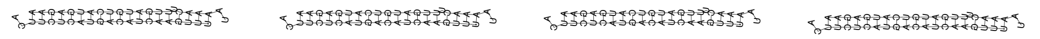 |
| *mir-5595* | 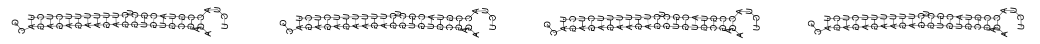 |
| *mir-55* | 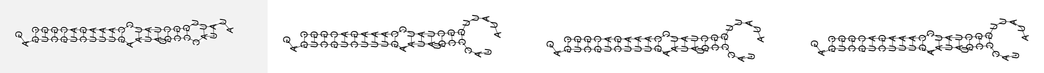 |
| *mir-56* | 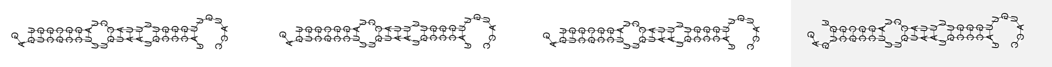 |
| *mir-57* | 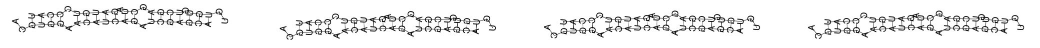 |

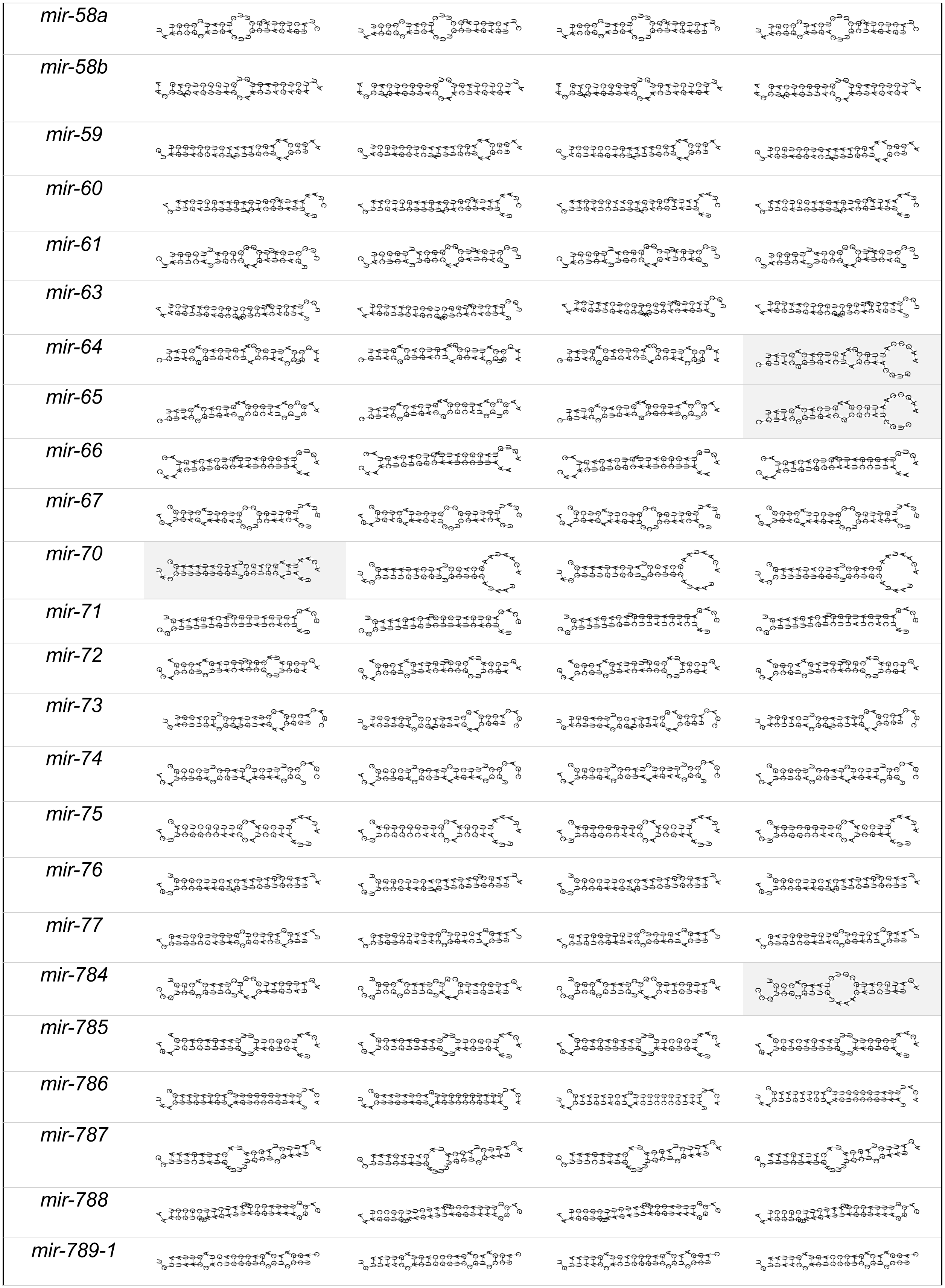
mir-58a
mir-58b
mir-59
mir-60
mir-61
mir-63
mir-64
mir-65
mir-66
mir-67
mir-70
mir-71
mir-72
mir-73
mir-74
mir-75
mir-76
mir-77
mir-784
mir-785
mir-786
mir-787
mir-788
mir-789-1

| | |
|---|---|
| *mir-790* | 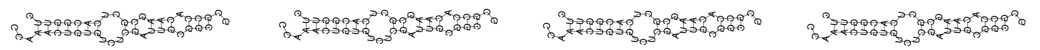 |
| *mir-791* | 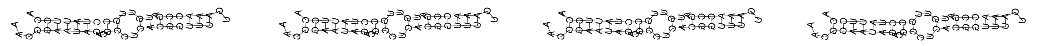 |
| *mir-792* | 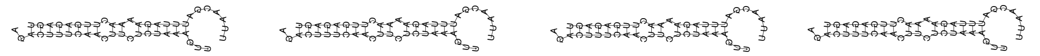 |
| *mir-794* | 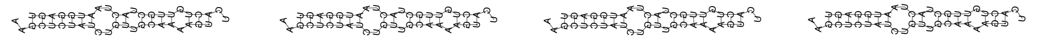 |
| *mir-795* | 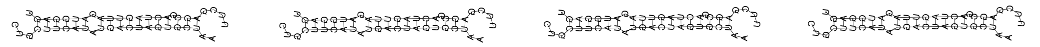 |
| *mir-797* | 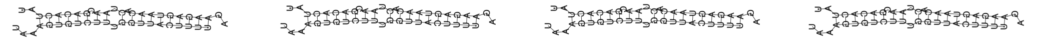 |
| *mir-79* | 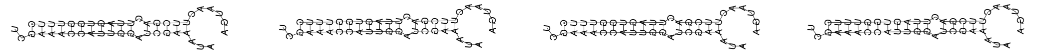 |
| *mir-800* | 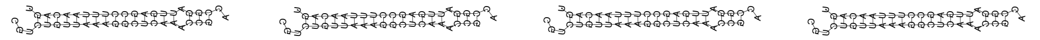 |
| *mir-80* | 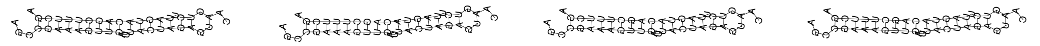 |
| *mir-8186* | 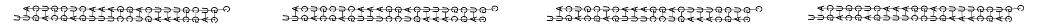 |
| *mir-8187* | 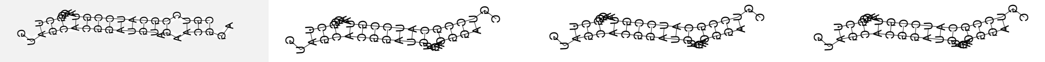 |
| *mir-8188* | 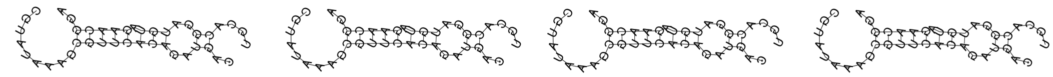 |
| *mir-8189* | 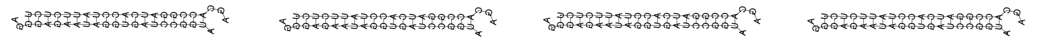 |
| *mir-8190* | 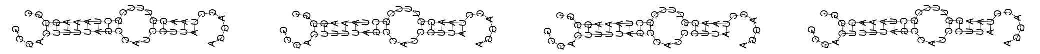 |
| *mir-8191* | 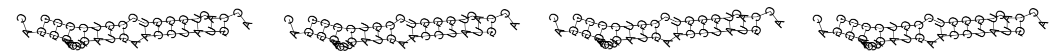 |
| *mir-8192* | 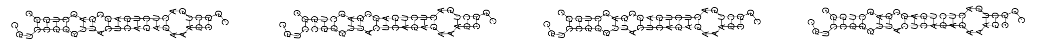 |
| *mir-8193* | 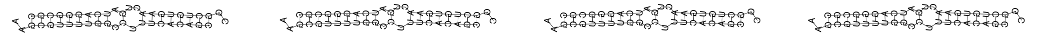 |
| *mir-8194* | 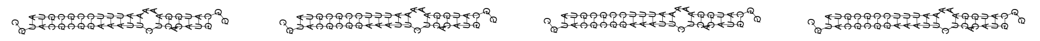 |
| *mir-8195* | 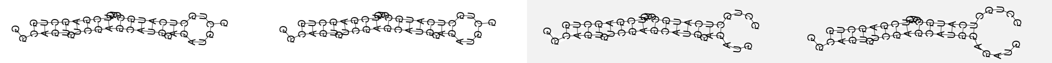 |
| *mir-8196a* | 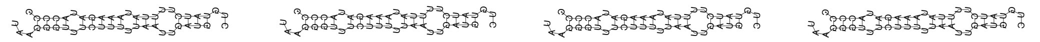 |
| *mir-8196b* | 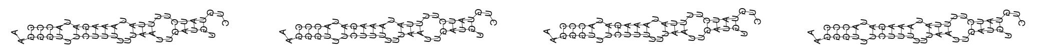 |
| *mir-8197* | 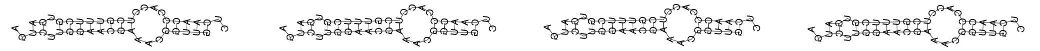 |
| *mir-8198* | 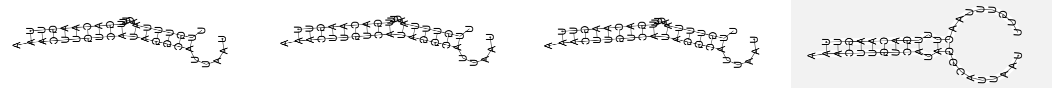 |

| | |
|---|---|
| *mir-8199* | 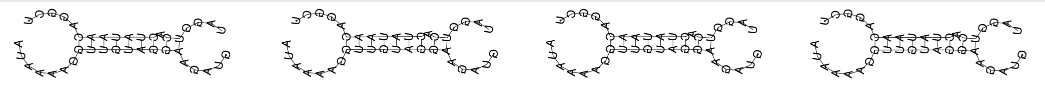 |
| *mir-81* | 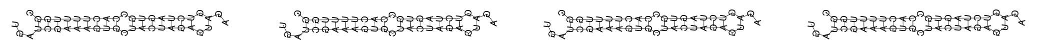 |
| *mir-8200* | 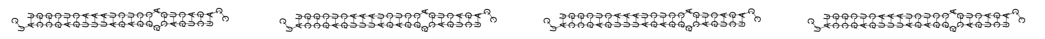 |
| *mir-8201* | 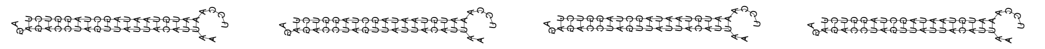 |
| *mir-8202* | 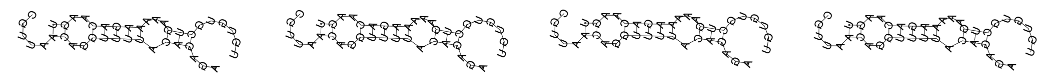 |
| *mir-8203* | 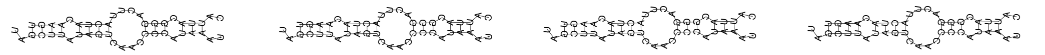 |
| *mir-8204* | 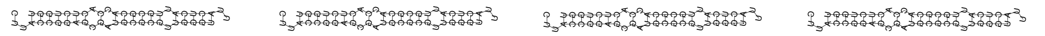 |
| *mir-8205* | 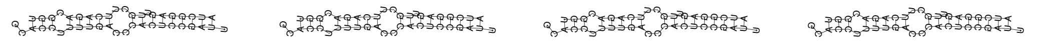 |
| *mir-8206* | 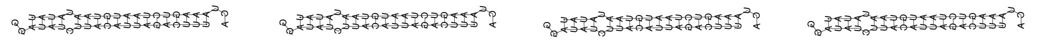 |
| *mir-8207* | 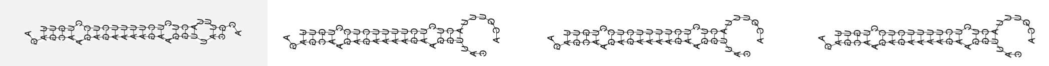 |
| *mir-8208* | 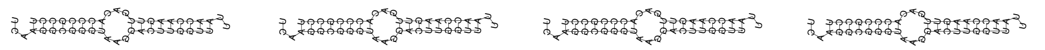 |
| *mir-8209* | 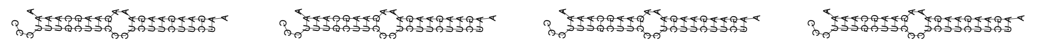 |
| *mir-8210* | 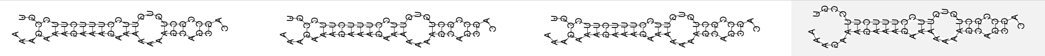 |
| *mir-8211* | 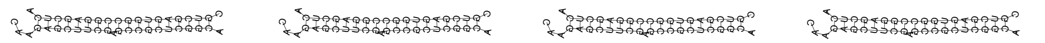 |
| *mir-8212* | 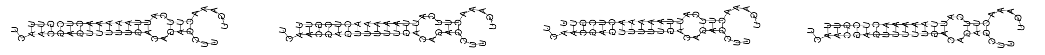 |
| *mir-82* | 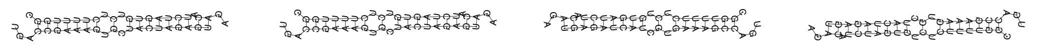 |
| *mir-83* | 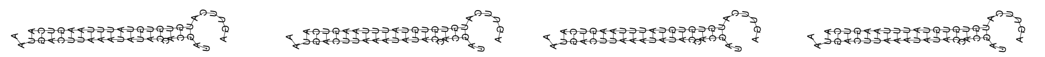 |
| *mir-84* | 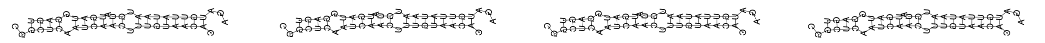 |
| *mir-85* | 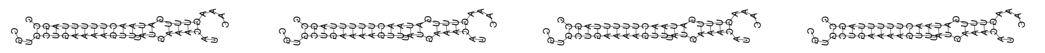 |
| *mir-86* | 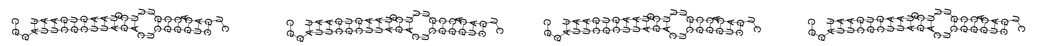 |
| *mir-87* | 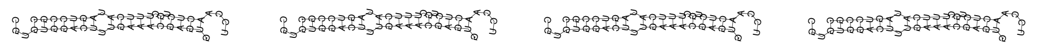 |
| *mir-90* | 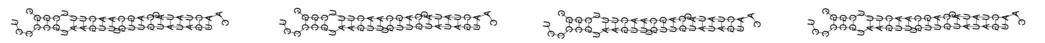 |

**Table S3:** *C. elegans* miRNA duplexes arising from precursor hairpins (constrained) compared to minimal free energy (MFE) duplex structures at 20°C

| miRNA | Constrained Duplex (20°C) | MFE Duplex (20°C) |
|---|---|---|
| *let-7* | U UU<br>UGAGGUAG AGGUUGUAUAG<br>AUUCCAUC UUUAACGUAUC<br>CC U | U UU<br>UGAGGUAG AGGUUGUAUAG<br>AUUCCAUC UUUAACGUAUC<br>CC U |
| *lin-4* | U U C A GA<br>CCC GAGA CUCA GUGU<br>GGG CUCU GGGU CACA<br>CCAU C C C | U U C A GA<br>CCC GAGA CUCA GUGU<br>GGG CUCU GGGU CACA<br>CCAU C C C |
| *lsy-6* | A U U<br>AAAUGCGUCU GUA CAAAA<br>UUUACGCAGA UAU GUUUU<br>AGC G – | A U U<br>AAAUGCGUCU GUA CAAAA<br>UUUACGCAGA UAU GUUUU<br>AGC G – |
| *mir-1019* | U U UC U UUUU<br>G GAGCA UGU GAGUU CA<br>C UUCGU ACA CUUAA GU<br>GAC U U C– U C | U U UC U UUUU<br>G GAGCA UGU GAGUU CA<br>C UUCGU ACA CUUAA GU<br>GAC U U C– U C |
| *mir-1020* | U CUU<br>G AAGUGUUACAGAAUAAU<br>C UUCACAGUGUCUUAUUA<br>GA U | U CUU<br>G AAGUGUUACAGAAUAAU<br>C UUCACAGUGUCUUAUUA<br>GA U |
| *mir-1022* | AA A GC– C<br>GAUCAUUGUU GGAC CAU<br>CUAGUAGUAA CCUG GUA<br>CGAC – AUA | AA A GC– C<br>GAUCAUUGUU GGAC CAU<br>CUAGUAGUAA CCUG GUA<br>CGAC – AUA |
| *mir-124* | C CUA A GU<br>GCAU CACC GUG CUUUA<br>CGUA GUGG CAC GGAAU<br>AC A CG– – | C CUA A GU<br>GCAU CACC GUG CUUUA<br>CGUA GUGG CAC GGAAU<br>AC A CG– – |
| *mir-1819* | A U AA G CA<br>AUCA GCUCAA CAUUC A<br>UAGU CGAGUU GUAAG U<br>AGG U A– G | A U AA GACA<br>AUCA GCUCAA CAUUC<br>UAGU CGAGUU GUAAG<br>AGG U A– GU |
| *mir-1820* | UU AU CG<br>UUUUGAUUGUUU CGAUG GUU<br>GAAACUAACAAA GUUAC CAA<br>A U– –– | UU AUGUUCG<br>UUUUGAUUGUUU CGAUG<br>GAAACUAACAAA GUUAC<br>A U– CAA |
| *mir-1821* | C UGC UU AU<br>UGCC AACU AGACU UCA<br>AUGG UUGA UCUGG AGU<br>AG A UAU –– | C UGC UCAAU<br>UGCC AACU AGACUUU<br>AUGG UUGA UCUGGAG<br>AG A UAU U |
| *mir-1822* | C – AA UA GGC<br>AGUUU UCUG GG AGC UC<br>UCAAA AGAC CC UCG AG<br>UC A U CG –– | C – AA AUCGGC<br>AGUUU UCUG GG AGCU<br>UCAAA AGAC CC UCGA<br>UC A U CG G |
| *mir-1823* | C C – A G UUU<br>AC CCUAA C CU U CAGUA<br>UG GGAUU G GA A GUCAU<br>AA A U U – G | C C – A G UUU<br>AC CCUAA C CU U CAGUA<br>UG GGAUU G GA A GUCAU<br>AA A U U – G |
| *mir-1824* | U C CU –– UU<br>GG AGUGUUU CCC CCAAC<br>CC UCACAAG GGG GGUUG<br>CCG U U– CC | U C CU –– UU<br>GG AGUGUUU CCC CCAAC<br>CC UCACAAG GGG GGUUG<br>CCG U U– CC |
| *mir-1829a* | A C U U UA<br>AGGGGA UUCUAAU GUU G<br>UCUCUU AAGGUUA CAA C<br>UUA U C – | A C U UGUA<br>AGGGGA UUCUAAU GUU<br>UCUCUU AAGGUUA CAA<br>UUA U C C |
| *mir-1829b* | A C UC A UA<br>AG GA UUCUAG UGGUUG<br>UC CU AAGGUC ACCAAC<br>UUA U UU – | A C UC A UA<br>AG GA UUCUAG UGGUUG<br>UC CU AAGGUC ACCAAC<br>UUA U UU – |
| *mir-1829c* | A C A A UA<br>AG GAAAUUC AG UGGUUG<br>UC CUUUAAG UC ACCAAC<br>UUA U G – | A C A A UA<br>AG GAAAUUC AG UGGUUG<br>UC CUUUAAG UC ACCAAC<br>UUA U G – |
| *mir-1830* | GU A CC<br>CGAG UUC CGUUUUCUAGG<br>GCUC AAG GUAAAGGAUCC<br>CG AA A | GU A CC<br>CGAG UUC CGUUUUCUAGG<br>GCUC AAG GUAAAGGAUCC<br>CG AA A |
| *mir-1832a* | G AA CU<br>CA CGAUUCG CUCCGCCCA<br>GU GCUAAGC GAGGCGGGU<br>UA A –– | G AA CU<br>CA CGAUUCG CUCCGCCCA<br>GU GCUAAGC GAGGCGGGU<br>UA A –– |
| *mir-1832b* | G UU<br>CAGCGAAUCGCUC GCCCACU<br>GUCGCUUAGCGAG CGGGUGA<br>UA A | G UU<br>CAGCGAAUCGCUC GCCCACU<br>GUCGCUUAGCGAG CGGGUGA<br>UA A |
| *mir-1* | C GC UA<br>CAUACUUC UUACAU CCA<br>GUAUGAAG AAUGUA GGU<br>AU A A– | C GC UA<br>CAUACUUC UUACAU CCA<br>GUAUGAAG AAUGUA GGU<br>AU A A– |
| *mir-2207* | A A AAG<br>UGUG AUUGAG CUGUGUAU<br>ACAC UAACUC GACACGUA<br>AC G G | A A AAG<br>UGUG AUUGAG CUGUGUAU<br>ACAC UAACUC GACACGUA<br>AC G G |
| *mir-2208a* | C A CC<br>AAGUGUACC GAAUCUG UAU<br>UUCAUAUGG CUUUGAC GUA<br>AC U – | C AUAUCC<br>AAGUGUACC GAAUCUG<br>UUCAUAUGG CUUUGAC<br>AC U GUA |
| *mir-2208b* | C A CC<br>AAGUGUACC GGAUCUG UAU<br>UUCACAUGG UUUAGAC GUA<br>AC U – | C AUAUCC<br>AAGUGUACC GGAUCUG<br>UUCACAUGG UUUAGAC<br>AC U GUA |

| | | |
|---|---|---|
| *mir-2209a* | – A CU C UC<br> G AGUGUAACC CU UCUC U<br> C UCACAUUGG GA AGAG A<br>A A C CU – | – A CU CUUC<br> G AGUGUAACC CU UCUC<br> C UCACAUUGG GA AGAG<br>A A C CU A |
| *mir-2209b* | AA U C UC<br>AGUGUAAC CUC UCUC U<br>UCGUGUUG GAG AGAG A<br>ACU GC U – | AA U CUUC<br>AGUGUAAC CUC UCUC<br>UCGUGUUG GAG AGAG<br>ACU GC U A |
| *mir-2209c* | – CAC G UU<br> G AGUGUAACCG GUCUU U<br> C UCACAUUGGC CAGAA A<br>A A CAC A | – CAC GUUU<br> G AGUGUAACCG GUCUU<br> C UCACAUUGGC CAGAA<br>A A CAC AA |
| *mir-2210* | A C U AAU GG<br>GG AGA CAAUC UUUUA<br>CC UCU GUUAG GAAAU<br>CAC A C CU– | A C U A UAGG<br>GG AGA CAAUC AUUUU<br>CC UCU GUUAG UGAAA<br>CAC A C C U |
| *mir-2211* | C A UUC C CG<br>CUCC UCUA UC AUCUGA<br>GAGG AGAU AG UGGACU<br>AAA – UUA A | C A UUC C CG<br>CUCC UCUA UC AUCUGA<br>GAGG AGAU AG UGGACU<br>AAA – UUA A |
| *mir-2212* | AG UAG U UG<br>UGGC AUCA GCU ACUU<br>ACCG UAGU CGG UGAA<br>CU AA UUA – | AG UAG U UG<br>UGGC AUCA GCU ACUU<br>ACCG UAGU CGG UGAA<br>CU AA UUA – |
| *mir-2213* | U A– C GA<br>GGCGG CUCUU ACAGUUU<br>CCGUC GAGAA UGUCGAA<br>AAU AG – | U A– C GA<br>GGCGG CUCUU ACAGUUU<br>CCGUC GAGAA UGUCGAA<br>AAU AG – |
| *mir-2215* | G CC<br>ACAGCACGUGUUACGAU CU<br>UGUUGUGCGCGAUGCUA GA<br>UU A | G CC<br>ACAGCACGUGUUACGAU CU<br>UGUUGUGCGCGAUGCUA GA<br>UU A |
| *mir-2216* | C GC<br>GCACAUUUUAAGU GGUAG<br>CGUGUAAAAUUCA CUAUC<br>UC U | C GC<br>GCACAUUUUAAGU GGUAG<br>CGUGUAAAAUUCA CUAUC<br>UC U |
| *mir-2217a* | A GUC U UC<br>C GAGUGGGCA GG GUCGA<br>G UUUGUCCGU CC CAGCU<br>UG C GUU – | A GUC U UC<br>C GAGUGGGCA GG GUCGA<br>G UUUGUCCGU CC CAGCU<br>UG C GUU – |
| *mir-2217b* | A GUC U UC<br>C GAGUGGGCA GG GUCAA<br>G UUUGUUCGU CC CAGUU<br>CG C GUU – | A GUC U UC<br>C GAGUGGGCA GG GUCAA<br>G UUUGUUCGU CC CAGUU<br>CG C GUU – |
| *mir-2218a* | AGU AA CA<br>CAAACUACA UUU GCCU<br>GUUUGAUGU AAG CGGA<br>AU GAU AC | AGU AA CA<br>CAAACUACA UUU GCCU<br>GUUUGAUGU AAG CGGA<br>AU GAU AC |
| *mir-2218b* | AG UC UC<br>ACUACAAACUACA AUUU<br>UGAUGUUUGAUGU UAAA<br>AGAG U– | AG UC UC<br>ACUACAAACUACA AUUU<br>UGAUGUUUGAUGU UAAA<br>AGAG U– |
| *mir-2219* | ACA A<br>GCUUUCUCUCGCAC UCGUC<br>CGAAAGGGAGCGUG AGCAG<br>A UC | ACA A<br>GCUUUCUCUCGCAC UCGUC<br>CGAAAGGGAGCGUG AGCAG<br>A UC |
| *mir-2220* | A U U UC<br>GUAAG CCAUAAAC AUU A<br>CAUUC GGUGUUUG UAA U<br>GA A U C | A U UAUC<br>GUAAG CCAUAAAC AUU<br>CAUUC GGUGUUUG UAA<br>GA A U CU |
| *mir-228* | A C C A A GG<br>AUGG ACUG AUGA UUC C<br>UACC UGGC UACU AGG G<br>CGC A A – C | A C C A A GG<br>AUGG ACUG AUGA UUC C<br>UACC UGGC UACU AGG G<br>CGC A A – C |
| *mir-229* | A G U CCA CG<br>AUGACACU G UAUCUUUU U<br>UACUGUGG C AUGGAAAG A<br>AG G U ––– | A G U CCAUCG<br>AUGACACU G UAUCUUUU<br>UACUGUGG C AUGGAAAG<br>AG G U A |
| *mir-230* | A – U UA<br>CUUGGUCG GCGAUU AAUAU<br>GGACCAGC UGUUGA UUAUG<br>AGA G – | A – U UA<br>CUUGGUCG GCGAUU AAUAU<br>GGACCAGC UGUUGA UUAUG<br>AGA G – |
| *mir-231* | A –– AC UA<br>CUG CUGUU UCA AGCUUG<br>GAC GACAA AGU UCGAAU<br>AA G CU GC | A –– AC UA<br>CUG CUGUU UCA AGCUUG<br>GAC GACAA AGU UCGAAU<br>AA G CU GC |
| *mir-232* | C UC AU UC<br>CUGCAGUU GAUG UUUA<br>GGCGUCAA CUAC AAAU<br>AGU UU GU | – UC AU UC<br>C CUGCAGUU GAUG UUUA<br>G GGCGUCAA CUAC AAAU<br>A U UU GU |
| *mir-233* | C – C C UA<br>UCGC CAU C CGUUGCUC AA<br>GGCG GUA G GUAACGAG UU<br>AG U C C – | C – C CAAUA<br>UCGC CAU C CGUUGCUC<br>GGCG GUA G GUAACGAG<br>AG U C C UU |
| *mir-234* | C CA U UA<br>GGUAUUC GAGU GAUAA<br>CCAUAAG CUCG UUAUU<br>UUC AG – | C CA U UA<br>GGUAUUC GAGU GAUAA<br>CCAUAAG CUCG UUAUU<br>UUC AG – |
| *mir-235* | UU CU U AAUU<br>AGGCC GG GA UGCAA<br>UCCGG CC CU ACGUU<br>AG C– CU C AU | UU CU U AAUU<br>AGGCC GG GA UGCAA<br>UCCGG CC CU ACGUU<br>AG C– CU C AU |
| *mir-236* | – UU A UAGA<br>CGUC UUACCUG CA UAUU<br>GCAG AAUGGAC GU AUAA<br>UC U U– C U | – UU A UAGA<br>CGUC UUACCUG CA UAUU<br>GCAG AAUGGAC GU AUAA<br>UC U U– C U |
| *mir-237* | C G UU A CU<br>UCC UGA AA CUCGA CAG<br>AGG ACU UU GAGCU GUC<br>CC A G UU – | C G UU A CU<br>UCC UGA AA CUCGA CAG<br>AGG ACU UU GAGCU GUC<br>CC A G UU – |

*mir-238*

```
        U C     C  U    GC                U C     C  U    GC
  UGGAUG U UCGGA GU CAAA            UGGAUG U UCGGA GU CAAA
  ACUUAC G AGCCU CA GUUU            ACUUAC G AGCCU CA GUUU
AG      C U     -  U              AG      C U     -  U
```

*mir-239a*

```
           C  A        GG                    C  A        GG
  UUUGUACUA AC UAGGUACU             UUUGUACUA AC UAGGUACU
  AAACGUGAU UG AUCUGUGA             AAACGUGAU UG AUCUGUGA
AC         C  -                   AC         C  -
```

*mir-239b*

```
         UA         A UG                   UA         A UG
  UUUGUAC  CACAAAAGU C              UUUGUAC  CACAAAAGU C
  AAACGUG  GUGUUUUCA G              AAACGUG  GUGUUUUCA G
AA       UG         C             AA       UG         C
```

*mir-240*

```
           U A A    A UGC                    U A A    AAUGC
  CGAGGAUUU G G CUAG A              CGAGGAUUU G G CUAG
  GCUUCUAAA C C GGUC U              GCUUCUAAA C C GGUC
UC         C C C    A             UC         C C C    AU
```

*mir-241*

```
          G  C    -   GA                    G  C    AAUGA
  UGAGGUAG UG GAGA AAU              UGAGGUAG UG GAGA
  ACUUCGUC AC CUCU UUA              ACUUCGUC AC CUCU
CU        G  U    G               CU        G  U    GUUA
```

*mir-243*

```
         G                                 G
  UAUCUCG UGCGAUCGUAC               UAUCUCG UGCGAUCGUAC
  AUAGGGC GCGCUAGCAUG               AUAGGGC GCGCUAGCAUG
CU       G           GC           CU       G           GC
```

*mir-244*

```
           UAC         UG                    UAC         UG
 UCUUUGGUUG   AAAGUGGUA          UCUUUGGUUG   AAAGUGGUA
 GGAAAUCGAC   UUUCGUCAU          GGAAAUCGAC   UUUCGUCAU
A          U--                    A          U--
```

*mir-245*

```
          C A  U   U   UG                   C A  U   UAAUUG
  GCUAUUUG A GG ACC AAU             GCUAUUUG A GG ACC
  CGAUGAAC U CC UGG UUA             CGAUGAAC U CC UGG
CU        C C  C   -              CU        C C  C   UUA
```

*mir-246*

```
  G    A   UUGU       UA           CG    A   UUGU       UA
 C CCUA CCG    CAUGUAA             CCUA CCG    CAUGUAA
 G GGAU GGC    GUACAUU             GGAU GGC    GUACAUU
C A    G   UUU-                   CGA    G   UUU-
```

*mir-247*

```
  U      A  U     A   CC            U      A  U     A   CC
   AGAGAA AG UUCUA UUA               AGAGAA AG UUCUA UUA
   UCUCUU UC GAGAU AGU               UCUCUU UC GAGAU AGU
UCU      A  C     C               UCU      A  C     C
```

*mir-249*

```
A       A    C   UU               A       A    C   UU
 GCAACGC CAAA GUC  CUGUG           GCAACGC CAAA GUC  CUGUG
 CGUUGCG GUUU CAG  GACAC           CGUUGCG GUUU CAG  GACAC
C       A    U   --     U         C       A    U   --     U
```

*mir-250*

```
   UU      C  C     CCG              UU      C  C     CCG
 CC  CAGUUG CU GUGAU               CC  CAGUUG CU GUGAU
 GG  GUCAAC GA CACUA               GG  GUCAAC GA CACUA
C  UU      U  -     A             C  UU      U  -     A
```

*mir-252*

```
AUAA    U  U  C C                 AUAA    U  U  C C
    GUAG AG GC G AGGUAA               GUAG AG GC G AGGUAA
    CGUC UC CG C UCCAUU               CGUC UC CG C UCCAUU
        U  -  U A      C                  U  -  U A      C
```

*mir-253*

```
           - A   C     CA                    - A   C     CA
  CUUUUCACA C CCU ACUAA             CUUUUCACA C CCU ACUAA
  GAAGGGUGU G GGA UGAUU             GAAGGGUGU G GGA UGAUU
GG         U C   -                GG         U C   -
```

*mir-254*

```
UACA AA  AU-      C  CA           UACAGAA  AU-      C  CA
    G  GC   AGAUUU CA                    GC   AGAUUU CA
    C  CG   UCUAAA GU                    CG   UCUAAA GU
     AG  CUU      C                   CAG  CUU      C
```

*mir-255*

```
               G     CUC                         G     CUC
  GUAAGAAAUCUUU UAGUU               GUAAGAAAUCUUU UAGUU
  CAUUUUUUAGAGA GUCAA               CAUUUUUUAGAGA GUCAA
GA             A     A            GA             A     A
```

*mir-259*

```
  A        -  U    U   A CA         A        -  U    U   AGCA
   AAUCUCAU CC AAUC GGU G            AAUCUCAU CC AAUC GGU
   UUAGGGUA GG UUAG CCA C            UUAGGGUA GG UUAG CCA
CAG        C  U    -   C          CAG        C  U    -   CC
```

*mir-2953*

```
          U   C    AAAU                     U   C    AAAU
  UACAGAAG GUU GUGA                 UACAGAAG GUU GUGA
  AUGUCUUC CGA CACU                 AUGUCUUC CGA CACU
CG        U   U    AG             CG        U   U    AG
```

*mir-2*

```
            -   G  U    UG                    -   G  U    UG
   CAUCAAAGC GGU GU GAUG             CAUCAAAGC GGU GU GAUG
   GUAGUUUCG CCG CA CUAU             GUAGUUUCG CCG CA CUAU
CGU         A   A  -              CGU         A   A  -
```

*mir-34*

```
   A       U  U      G UG            A       U  U      GUUG
    GGCAGUG GG UAGCUG U               GGCAGUG GG UAGCUG
    CCGUCAC UC AUCGGC A               CCGUCAC UC AUCGGC
CCCA       U  C      -            CCCA       U  C      A
```

*mir-354*

```
   U     U       -   AU              U     U       -   AU
 GG GCGGC GCAGACG GGU              GG GCGGC GCAGACG GGU
 CC CGUCG UGUUUGU CCA              CC CGUCG UGUUUGU CCA
U  U     U       U                U  U     U       U
```

*mir-355*

```
   U         CU  -    UG            UU         CU  -    UG
  U UGUUUUAGC  GA GCUA                UGUUUUAGC  GA GCUA
  G ACAAAAUCG  CU CGAU                ACAAAAUCG  CU CGAU
CC U         UU  U                CCGU         UU  U
```

*mir-356b*

```
      -   AC   U  U      U                -   AC   U  U      U
  UGGU GAG  ACG CG AACGAA           UGGU GAG  ACG CG AACGAA
  ACCA CUC  UGC GC UUGUUU           ACCA CUC  UGC GC UUGUUU
AC    A   GU   -  -               AC    A   GU   -  -
```

*mir-357*

```
  C   A     -   C    A GC           C   A     -   C    AUGC
   CCU CAACG CUG GCAU U              CCU CAACG CUG GCAU
   GGA GUUGC GAC CGUA A              GGA GUUGC GAC CGUA
UGA   C     U   -    A            UGA   C     U   -    AA
```

*mir-358*

```
      G C    C  U    CUGU                 G C    C  U    CUGU
  ACCU G CAGG AU CCAA               ACCU G CAGG AU CCAA
  UGGA C GUCC UA GGUU               UGGA C GUCC UA GGUU
UC    A U    C  U    A            UC    A U    C  U    A
```

**mir-35**

```
             U  A A    UA
  UGCUGGUUUCU CC C GUGG
  ACGAUCAAAGG GG G CACU
UG           U  - C

             U  A A    UA
  UGCUGGUUUCU CC C GUGG
  ACGAUCAAAGG GG G CACU
UG           U  - C
```

**mir-360**

```
      C  UU
UUGUGA CG  GUUACGGUCA
AACACU GC  UAAUGCCAGU
      U  CC

      C  UU
UUGUGA CG  GUUACGGUCA
AACACU GC  UAAUGCCAGU
      U  CC
```

**mir-36**

```
  C  C           CA   CUA
   GC AAUUUUCGCUU  GUG
   CG UUAAAAGUGGG  CAC
GUA  C           C-   U

  C  C           CA   CUA
   GC AAUUUUCGCUU  GUG
   CG UUAAAAGUGGG  CAC
GUA  C           C-   U
```

**mir-37**

```
           C    G     CUA
  UGUGGGUGU CGUU CGGUG
  ACGUUCACA GUGG GCCAC
UG         A    -     U

           C    G     CUA
  UGUGGGUGU CGUU CGGUG
  ACGUUCACA GUGG GCCAC
UG         A    -     U
```

**mir-38**

```
             C G      UA
  UCCGGUUUUUU C UGGUGA
  AGGUCAAAAAG G GCCACU
UG           A G

             C G      UA
  UCCGGUUUUUU C UGGUGA
  AGGUCAAAAAG G GCCACU
UG           A G
```

**mir-392**

```
   G   U         G    U
  A CAU CGUGGUUGA GAUA
  U GUG GCACUAGCU CUAU
AG A   U         A

  AG   U         G    U
    CAU CGUGGUUGA GAUA
    GUG GCACUAGCU CUAU
AGUA   U         A
```

**mir-39**

```
           -  U      A UA
  AGCUGAUUU CG CUUGGU A
  UCGACUAAA GU GGGCCA U
GU         U  -      C

           -  U      AAUA
  AGCUGAUUU CG CUUGGU
  UCGACUAAA GU GGGCCA
GU         U  -      CU
```

**mir-40**

```
     G          A  A   UA
  AGU GAUGUAUGCC UG UGA
  UCG CUACAUGUGG GC ACU
AA   A          -  C

     G          A  A   UA
  AGU GAUGUAUGCC UG UGA
  UCG CUACAUGUGG GC ACU
AA   A          -  C
```

**mir-41**

```
             U  GCA    UA
  GGUGGUUUUUC CU   GUGA
  CCACUAAAAAG GG   CACU
AU           U  GC-

             -    CA    UA
  GGUGGUUUUUC UCUG  GUGA
  CCACUAAAAAG GGGC  CACU
AU           U    --
```

**mir-42**

```
            U   UUUU      AG
   GUGGGUGUU GCU    CGGUGA
   CAUCUACAA UGG    GCCACU
AGA         U   ----

            U   UUUU      AG
   GUGGGUGUU GCU    CGGUGA
   CAUCUACAA UGG    GCCACU
AGA         U   ----
```

**mir-43**

```
      U    -     A     UAUG
  GACA CAAG AAACU GUGAU
  CUGU GUUC UUUGA CACUA
CG    C    A     -     U

      U    -     A     UAUG
  GACA CAAG AAACU GUGAU
  CUGU GUUC UUUGA CACUA
CG    C    A     -     U
```

**mir-44**

```
           C  G       UA
  CUGGAUGUG UC UUGGUCA
  GACUUACAC AG GAUCAGU
UC         -  A

           C  G       UA
  CUGGAUGUG UC UUGGUCA
  GACUUACAC AG GAUCAGU
UC         -  A
```

**mir-45**

```
           C  G       UA
  CUGGAUGUG UC UUAGUCA
  GACUUACAC AG GAUCAGU
UC         -  A

           C  G       UA
  CUGGAUGUG UC UUAGUCA
  GACUUACAC AG GAUCAGU
UC         -  A
```

**mir-46**

```
          CG-     U    GU
  AAGAGAGC   UCUAU GACA
  UUCUCUCG   AGGUA CUGU
AC        CUG     -

          CG-     U    GU
  AAGAGAGC   UCUAU GACA
  UUCUCUCG   AGGUA CUGU
AC        CUG     -
```

**mir-47**

```
          A    AU-     GU
  AAGAGAGC GUCU   UGACA
  UUCUCUCG CGGA   ACUGU
AC        -    GGU

          A    AU-     GU
  AAGAGAGC GUCU   UGACA
  UUCUCUCG CGGA   ACUGU
AC        -    GGU
```

**mir-4805**

```
       C      G C      GC
  UGCGG AAAUUU C GAAUUU
  ACGCC UUUAGA G UUUAAA
AC     U      - U

       C      G C      GC
  UGCGG AAAUUU C GAAUUU
  ACGCC UUUAGA G UUUAAA
AC     U      - U
```

**mir-4806**

```
   A              C  GC
  C CUUACCGGCUGAGC AU
  G GAAUGGCCGACUCG UA
CG A              A

  CA              CAUGC
    CUUACCGGCUGAGC
    GAAUGGCCGACUCG
CGGA              AUA
```

**mir-4808**

```
        U   U   U    UC
  GUAAGA AGA UAG GCUU
  CAUUCU UCU AUC CGAA
GA      C   U   U

        U   U   U    UC
  GUAAGA AGA UAG GCUU
  CAUUCU UCU AUC CGAA
GA      C   U   U
```

**mir-4809**

```
   U         -    A    UC
  G AAGUUCAGA GUUG AUUA
  C UUCGGGUCU CAAC UAAU
GA C         U    A

   U         -    A    UC
  G AAGUUCAGA GUUG AUUA
  C UUCGGGUCU CAAC UAAU
GA C         U    A
```

**mir-4810b**

```
         C     -      UAU
  GUAGGUU AUGAG UAGUCA
  CAUUCAA UACUC AUCAGU
GA       C     U

         C     -      UAU
  GUAGGUU AUGAG UAGUCA
  CAUUCAA UACUC AUCAGU
GA       C     U
```

**mir-4811**

```
            CU      A AG
  UGAACAAUAC  GUGUUA A
  GCUUGUUAUG  CACAAU U
UC          CU      A

  U         CU      AAAG
   GAACAAUAC  GUGUUA
   CUUGUUAUG  CACAAU
UCG         CU      AU
```

**mir-4812**

```
 A    C C        A    U
  AGAG G UUGUAGUG GUUG
  UCUC U AACGUCAC CAAC
UA    U U        -

 A    C C        A    U
  AGAG G UUGUAGUG GUUG
  UCUC U AACGUCAC CAAC
UA    U U        -
```

**mir-4813**

```
           A  AA        CU
  AGACUAUCU GG  AUAUUGAA
  UCUGGUAGG CC  UAUAACUU
GG         -  A-

           A  AA        CU
  AGACUAUCU GG  AUAUUGAA
  UCUGGUAGG CC  UAUAACUU
GG         -  A-
```

**mir-4814**

```
                  G   CU
  UUCUCAACCAACUUUG CCA
  AAGAGUUGGUUGAAAC GGU
UG                G

                  G   CU
  UUCUCAACCAACUUUG CCA
  AAGAGUUGGUUGAAAC GGU
UG                G
```

**mir-4816**

```
         - UUU-        CAA
  GUAAGUG G    UUGUAGAU
  CAUUCAC C    AACAUCUA
GA       G UUUU

         - UUU-        CAA
  GUAAGUG G    UUGUAGAU
  CAUUCAC C    AACAUCUA
GA       G UUUU
```

**mir-48**

```
      G      CA       CGA
  UGAG UAGGCU  GUAGAUG
  GCUC AUCCGA  CACCUAC
AC    G      A-       A

  U   G      CA       CGA
   GAG UAGGCU  GUAGAUG
   CUC AUCCGA  CACCUAC
ACG   G      A-       A
```

**mir-49**

```
  C       G     AU      CC
   GCAGUUU UUGUG  GUGCU
   CGUCGAA AGCAC  CACGA
AGA       G     --      A

  C       G     AU      CC
   GCAGUUU UUGUG  GUGCU
   CGUCGAA AGCAC  CACGA
AGA       G     --      A
```

**mir-50**

```
                 UCU    UU
  UGAUAUGUCUGGUAU   UGGG
  GCUAUGCAGAUUAUA   GCCC
CA               -C-

                 UCU    UU
  UGAUAUGUCUGGUAU   UGGG
  GCUAUGCAGAUUAUA   GCCC
CA               -C-
```

**mir-51**

```
      C   G  C  A      UU
  UACC GUA CU CU UCCAUG
  GUGG CAU GA GA AGGUAC
AC    A   G  C  -

      C   G  C  A      UU
  UACC GUA CU CU UCCAUG
  GUGG CAU GA GA AGGUAC
AC    A   G  C  -
```

**mir-52**

```
  C    GUA   A   UUC    CU
   ACCC   CAU UGU   CGUG
   UGGG   GUA ACA   GCAC
CGA    AAA   -   UU-

  C    GUA   A   UUC    CU
   ACCC   CAU UGU   CGUG
   UGGG   GUA ACA   GCAC
CGA    AAA   -   UU-
```

**mir-53**

```
  C       CAU    UU     CU
   ACCCGUA   UUGU  CCGUG
   UGGGUAU   AACA  GGCAC
CGC       AU-    C-

  C       CAU    UU     CU
   ACCCGUA   UUGU  CCGUG
   UGGGUAU   AACA  GGCAC
CGC       AU-    C-
```

**mir-54**

```
  A    -    -  CG   A A CA
   GGAU AUGA GA  ACG G A
   CCUA UACU CU  UGC C U
GAG    A    U  AA   C A

  A    -    -  CG   AGAACA
   GGAU AUGA GA  ACG
   CCUA UACU CU  UGC
GAG    A    U  AA   CCAU
```

**mir-5545**

```
   C                 AA GC
  G CGGUUUGAUCUACAAAA  U
  C GCCAAACUAGAUGUUUU  A
UA A                 C-

   C                 AAUGC
  G CGGUUUGAUCUACAAAA
  C GCCAAACUAGAUGUUUU
UA A                 CA
```

**mir-5546**

```
      UUU   -         UCGU
  ACCC   CGC CAAUUUUUU
  UGGG   GCG GUUAAAAAG
GU    UAC   C

      UUU   -         UCGU
  ACCC   CGC CAAUUUUUU
  UGGG   GCG GUUAAAAAG
GU    UAC   C
```

**mir-5547**

```
            A  A      AU
  CAACUUUUAG CC AUAGGC
  GUUGAAAAUC GG UAUCCG
CU          C  C

            A  A      AU
  CAACUUUUAG CC AUAGGC
  GUUGAAAAUC GG UAUCCG
CU          C  C
```

**mir-5548**

```
  G       A UC       C    GG
   CCUUCUC C  UCCACGG GGUA
   GGAAGAG G  AGGUGUC CCGA
AGA       - UU       -

  G       A UC       C    GG
   CCUUCUC C  UCCACGG GGUA
   GGAAGAG G  AGGUGUC CCGA
AGA       - UU       -
```

**mir-5549**

```
   -UUGU             GU
  C     GAAAUUAACGUGA
  G     UUUUGGUUGUACU
UG UUGUU

   CUUGU             GU
        GAAAUUAACGUGA
        UUUUGGUUGUACU
UGGUUGUU
```

**mir-5550**

```
C        A           C
 CCCGCCCA GAUUUCAUUUG
 GGGCGGGU CUAAAGUAAAC
         G

C        A           C
 CCCGCCCA GAUUUCAUUUG
 GGGCGGGU CUAAAGUAAAC
         G
```

**mir-5551**

```
     A      U    UC    AU
  UGU AAUGGU GGAA  UGGU
  ACG UUACCA UCUU  ACCA
CU   A      U    UA

     A      U    UC    AU
  UGU AAUGGU GGAA  UGGU
  ACG UUACCA UCUU  ACCA
CU   A      U    UA
```

**mir-5552**

```
     A           A     CC
  UGU GUUUGUAGUCU GCAGA
  ACA CAAACAUCAGA CGUCU
AC   C           C

      A           A     CC
  UGU GUUUGUAGUCU GCAGA
  ACA CAAACAUCAGA CGUCU
AC   C           C
```

**mir-5553**

```
          C  C         UU
  UUGCCACG GC AUCCAUUGA
  AACGGUGC CG UGGGUAACU
AG        A  A

          C  C         UU
  UUGCCACG GC AUCCAUUGA
  AACGGUGC CG UGGGUAACU
AG        A  A
```

**mir-5592**

```
                       UG
  CGGCCCUUACCGUUUAAUACA
  GCCGGGAAUGGCAAAUUAUGU
CG

                       UG
  CGGCCCUUACCGUUUAAUACA
  GCCGGGAAUGGCAAAUUAUGU
CG
```

**mir-5593**

```
   A   C      AU      UC
  G UGG UGGAAU  CGGUAU
  C ACC ACCUUA  GCCAUA
AA A   A      C-

   A   C      AU      UC
  G UGG UGGAAU  CGGUAU
  C ACC ACCUUA  GCCAUA
AA A   A      C-
```

**mir-5594**

```
                 U    AU
  AAGAGUACUGUAGUU CAAA
  UUCUCAUGACAUCAA GUUU
AC               -

                 U    AU
  AAGAGUACUGUAGUU CAAA
  UUCUCAUGACAUCAA GUUU
AC               -
```

**mir-5595**

```
             U       CA CU
  UCUCUUUUUUC CGCAUGC  U
  AGAGAGAAGAG GUGUGCG  A
GC           -       AG

             U       - AUCU
  UCUCUUUUUUC CGCAUGC C
  AGAGAGAAGAG GUGUGCG G
GC           -       A A
```

**mir-55**

```
            C   -   U  UA
  CGGCAGAAAC UAU CGG  UA
  GUCGUCUUUG AUA GCC  AU
GA          A   U   C

            C   -   UUAUA
  CGGCAGAAAC UAU CGG
  GUCGUCUUUG AUA GCC
GA          A   U   CAU
```

**mir-56**

```
         UC    U     UGUACC
  UGGCGGA  CAUU UGGGU
  GUCGCCU  GUAA GCCCA
GA       UU    U     U

         UC    U     UGUACC
  UGGCGGA  CAUU UGGGU
  GUCGCCU  GUAA GCCCA
GA       UU    U     U
```

**mir-57**

```
      C     A  G    U   GU
  UACC UGUAG UC AGCU UGU
  GUGG ACAUC AG UCGA GCA
AC    A     -  A    -

      C     A  G    U   GU
  UACC UGUAG UC AGCU UGU
  GUGG ACAUC AG UCGA GCA
AC    A     -  A    -
```

**mir-58a**

```
      C    CUU  U      UC
  UGCC UACU   CG AUCUCA
  ACGG AUGA   GC UAGAGU
UA    C    CUU  -

      C    CUU  U      UC
  UGCC UACU   CG AUCUCA
  ACGG AUGA   GC UAGAGU
UA    C    CUU  -
```

**mir-58b**

```
     -       UG        UA
   GA CUCGGUG  UGAUCUCU
   CU GAGUUAC  ACUAGAGA
AAC  A       CA

     -       UG        UA
   GA CUCGGUG  UGAUCUCU
   CU GAGUUAC  ACUAGAGA
AAC  A       CA
```

**mir-59**

```
           -      AA   AA
  UCGUCCUGA AAACGA  CGG
  AGUAGGACU UUUGCU  GCU
GU         A      AA

           -      AA   AA
  UCGUCCUGA AAACGA  CGG
  AGUAGGACU UUUGCU  GCU
GU         A      AA
```

**mir-60**

```
            -    C    A UC
  AACUGGAAGA GUGC AUAA A
  UUGAUCUUUU CACG UAUU U
AC          A    -    A

            -    C    AAUC
  AACUGGAAGA GUGC AUAA
  UUGAUCUUUU CACG UAUU
AC          A    -    AU
```

**mir-61**

```
       U    GG  U    CUU
  UGGGU ACGG  CU AGUC
  ACUCA UGCC  GA UCAG
CU     U    AA  -    U

       U    GG  U    CUU
  UGGGU ACGG  CU AGUC
  ACUCA UGCC  GA UCAG
CU     U    AA  -    U
```

**mir-63**

```
            --    A     CGU
  UCUAACUCGU  CGGU GUCAU
  AGGUUGAGCG  GUCA CAGUA
AA          AA    -     U

            --    A     CGU
  UCUAACUCGU  CGGU GUCAU
  AGGUUGAGCG  GUCA CAGUA
AA          AA    -     U
```

**mir-64**

```
     A      AG   --   CGAA
 UAUG CACUGA  CGU  UAC
 GUAC GUGACU  GCA  GUG
C    G      A-   AC

     A      AG    A - GAA
 UAUG CACUGA  CGUU C C
 GUAC GUGACU  GCAA G G
C    G      A-    C U
```

**mir-65**

```
     A     AA      A C AA
 UAUG CACUG  GCGUA C G
 GUAC GUGAC  CGCAU G C
C    C     G-      C U

     A     AA      A C AA
 UAUG CACUG  GCGUA C G
 GUAC GUGAC  CGCAU G C
C    C     G-      C U
```

**mir-66**

```
 CA        A        G GA
   UGACACUG UUAGGGAU U
   ACUGUGGC AAUCCUUA A
CAA        -        A

 CA        A        GUGA
   UGACACUG UUAGGGAU
   ACUGUGGC AAUCCUUA
CAA        -        AA
```

**mir-67**

```
  C    A     CC-      U UG
   GCUC UUCUG   GGUUGU A
   UGAG AAGAU   CCAACA U
AGA    A     CCU      C

  C    A     CC-      UAUG
   GCUC UUCUG   GGUUGU
   UGAG AAGAU   CCAACA
AGA    A     CCU      CU
```

**mir-70**

```
  C         U     A    CA
   GAAAUACUA CGACG AUAA
   CUUUGUGGU GCUGC UAAU
UAC         U     A

  C         U     AAUAACA
   GAAAUACUA CGACG
   CUUUGUGGU GCUGC
UAC         U     AUAAU
```

**mir-71**

```
          U         G CG
 UGAAAGACA GGGUAGUGA A
 GCUUUUUGU CUUAUCACU U
C         -         A

 U        U         GACG
  GAAAGACA GGGUAGUGA
  CUUUUUGU CUUAUCACU
CG        -         AU
```

**mir-72**

```
  A    A     U   AU    GA
   GGCA GAUGU GGC  AGCU
   CCGU UUACA CCG  UCGA
GCA    C     -   CU

  A    A     U   AU    GA
   GGCA GAUGU GGC  AGCU
   CCGU UUACA CCG  UCGA
GCA    C     -   CU
```

**mir-73**

```
        U  -     GA    CAGC
  UGGACU CC AUAUC  GCCA
  ACUUGA GG UGUAG  CGGU
UG      C  A     AA

        U  -     GA    CAGC
  UGGACU CC AUAUC  GCCA
  ACUUGA GG UGUAG  CGGU
UG      C  A     AA
```

**mir-74**

```
  C     U    C    UC   GC
   GGGCU CCAU UCUU  CCA
   UCUGA GGUA AGAA  GGU
ACA     C    A    C-

  C     U    C    U  CAGC
   GGGCU CCAU UCUU CC
   UCUGA GGUA AGAA GG
ACA     C    A    C  U
```

**mir-75**

```
  C         CA     A  UA
   AGUCGGUUG  AGCUU AA
   UCGGCCAAC  UCGAA UU
ACU         CA     A

  C         CA     AAAUA
   AGUCGGUUG  AGCUU
   UCGGCCAAC  UCGAA
ACU         CA     AUU
```

**mir-76**

```
  U        -      U    UA
   GGGCUUCA CAAUAG CGAA
   UCCGAAGU GUUGUU GCUU
AGU        A      -

  U        -      U    UA
   GGGCUUCA CAAUAG CGAA
   UCCGAAGU GUUGUU GCUU
AGU        A      -
```

**mir-77**

```
            C     G   AU
  GAUGGUUGUG UCUGA GAA
  CUGUCGAUAC GGACU CUU
AC          C     A

            C     G   AU
  GAUGGUUGUG UCUGA GAA
  CUGUCGAUAC GGACU CUU
AC          C     A
```

**mir-784**

```
      A    C GC        GA
  UGGC CAAU U  GUACGUA
  GUCG GUUG A  CAUGUAU
CC    C    U AA

  U   A    C GC        GA
   GGC CAAU U  GUACGUA
   UCG GUUG A  CAUGUAU
CCG   C    U AA
```

**mir-785**

```
  A         UU      A CA
   GCACAGAAU  UUCGCU A
   UGUGUUUUG  AAGUGA U
AGA         UU      A

  A         UU      AACA
   GCACAGAAU  UUCGCU
   UGUGUUUUG  AAGUGA
AGA         UU      AU
```

**mir-786**

```
  C        G          U CA
   GAAUAUCA UUGGGGUAUU A
   CUUGUAGU AGUCCCGUAA U
UAA        A          -

  C        G          UACA
   GAAUAUCA UUGGGGUAUU
   CUUGUAGU AGUCCCGUAA
UAA        A          U
```

**mir-787**

```
          -AU-    U    CA
  AAAGAUAC    ACGA CUUA
  UUUCUAUG    UGCU GAAU
GC        AUUU    C

          -AU-    U    CA
  AAAGAUAC    ACGA CUUA
  UUUCUAUG    UGCU GAAU
GC        AUUU    C
```

**mir-788**

```
       --     CU       G AG
  UCCGC  UUCUAA  UCCAUUU C
  AGGUG  AAGAUU  AGGUAAA G
AA     GC     --       G

       --     CU       G AG
  UCCGC  UUCUAA  UCCAUUU C
  AGGUG  AAGAUU  AGGUAAA G
AA     GC     --       G
```

**mir-789-1**

```
       A           A   C
  AAUUG UGACCCAGACA GGA
  UUAAC ACUGGGUCCGU CCU
UG     C           C

       A           A   C
  AAUUG UGACCCAGACA GGA
  UUAAC ACUGGGUCCGU CCU
UG     C           C
```

**mir-790**

```
  C       UC-  G   A   CG
   UUGGCAC   GC AAC CCG
   AACUGUG   CG UUG GGC
CCA       UCU  A   C

  C       UC-  G   A   CG
   UUGGCAC   GC AAC CCG
   AACUGUG   CG UUG GGC
CCA       UCU  A   C
```

**mir-791**

```
  A       --  UU  A      GU
   CCUUAUC  CG  GU GCCAAA
   GGAAUAG  GC  CA CGGUUU
AAC       AC  CU  -

  A       --  UU  A      GU
   CCUUAUC  CG  GU GCCAAA
   GGAAUAG  GC  CA CGGUUU
AAC       AC  CU  -
```

**mir-792**

```
          C  A      AG    UU
  UGAGAGUU AA AGAUUU  CAA
  ACUUUCAA UU UCUAAA  GUU
AG        C  C      --

          C  A      AGCAAUU
  UGAGAGUU AA AGAUUU
  ACUUUCAA UU UCUAAA
AG        C  C      GUU
```

**mir-794**
```
         AU  U    G   CU                   AU  U    G   CU
  UGAGGUA  CA CGUU UCA              UGAGGUA  CA CGUU UCA
  GCUCUAU  GU GCAA AGU              GCUCUAU  GU GCAA AGU
AA       CU  U    A               AA       CU  U    A
```

**mir-795**
```
         G          G GCUU          U      G           AGCUU
  UGAGGUA AUUGAUCAGC A               GAGGUA AUUGAUCAGCG
  GCUUCAU UGACUAGUGC U               CUUCAU UGACUAGUGCU
CU       A          – AA          CUG      A           AA
```

**mir-797**
```
  A      C  U A         GA         UA      C  U A         GA
 U UCACAG AA C CAAUGAGAA             UCACAG AA C CAAUGAGAA
 A AGUGUC UU G GUUACUUUU             AGUGUC UU G GUUACUUUU
U A      –  U –                   UAA      –  U –
```

**mir-79**
```
             C     CA  GA                      C     CAAUGA
  CUUUGGUGAUU AGCUU  AU             CUUUGGUGAUU AGCUU
  GAAACCAUUGG UCGAA  UA             GAAACCAUUGG UCGAA
UC           A     A–             UC           A     AUA
```

**mir-800**
```
  U               A   CA            U               A   CA
   GACAAUUUCCGAGUU GGC               GACAAUUUCCGAGUU GGC
   CUGUUAAAGGCUCAA CCG               CUGUUAAAGGCUCAA CCG
CGU               A               CGU               A
```

**mir-80**
```
  A         ––      U  G AC         A         ––      U  G AC
   GCUUUCGAC  AUGAU CU A             GCUUUCGAC  AUGAU CU A
   CGAAAGUUG  UACUA GA U             CGAAAGUUG  UACUA GA U
AGC         AU      –  G          AGC         AU      –  G
```

**mir-8186**
```
                      C                                 C
 ACUGCUCAAAGGACUUUGCUG             ACUGCUCAAAGGACUUUGCUG
 UGACGAGUUUCCUGAAACGAC             UGACGAGUUUCCUGAAACGAC
U                                 U
```

**mir-8187**
```
     GGAA       –––––    GC            GGAA       –––––    GC
  UCG    UGCCUAC     GCCU           UCG    UGCCUAC     GCCU
  AGC    ACGGAUG     CGGA           AGC    ACGGAUG     CGGA
GU   ––––       UAGAA             GU   ––––       UAGAA
```

**mir-8188**
```
       AG     A   AG  ACGU               AG     A   AG  ACGU
         GCAAG UGU  GC                     GCAAG UGU  GC
         CGUUC ACA  UG                     CGUUC ACA  UG
CGUAUAAAG     –   GA  AC          CGUAUAAAG     –   GA  AC
```

**mir-8189**
```
                     CGA                               CGA
  UCUCUUAUCCACUAGGCCA               UCUCUUAUCCACUAGGCCA
  GGAGAAUAGGUGAUCCGGU               GGAGAAUAGGUGAUCCGGU
AG                   A            AG                   A
```

**mir-8190**
```
  CG        CUUU     CCAGGA         CG        CUUU     CCAGGA
    GGAAAUCG    GGAAU                 GGAAAUCG    GGAAU
    CUUUUAGC    CCUUA                 CUUUUAGC    CCUUA
GCGA        CAU–                  GCGA        CAU–
```

**mir-8191**
```
     –––––    C     C  CA              –––––    C     C  CA
  CCC     CUGC UGGGU AC             CCC     CUGC UGGGU AC
  GGG     GAUG ACCUA UG             GGG     GAUG ACCUA UG
CA   AACCU    A     –             CA   AACCU    A     –
```

**mir-8192**
```
  C    G  C       AG   GC           C    G  C       AG   GC
   GGUC AG GAGUCUC  UCG              GGUC AG GAGUCUC  UCG
   CCGG UU CUCAGAG  AGC              CCGG UU CUCAGAG  AGC
CGU    G  A       AA              CGU    G  A       AA
```

**mir-8193**
```
           A UC         GC                   A UC         GC
  CGCGGGACU G  AAGUGUCG             CGCGGGACU G  AAGUGUCG
  GCGUUUUGG C  UUCACAGC             GCGUUUUGG C  UUCACAGC
AA         C U–                   AA         C U–
```

**mir-8194**
```
              AA  –    GG                   AA  –    GG
  AUGCGCCUUUAA  AG GUAC             AUGCGCCUUUAA  AG GUAC
  UACGCGGAAAUU  UC CAUG             UACGCGGAAAUU  UC CAUG
CG            C–  A               CG            C–  A
```

**mir-8195**
```
      –    –  –  G–   UCGUCG            –    GUC     –  GU G
  GUCG AGCU GU CC  UAC              GUCG AGCU   CGUAC  UC C
  CAGU UCGA CA GG  AUG              CAGU UCGA   GCAUG  AG G
GG    G    G  U  AG               GG    G    –––     G  AU
```

**mir-8196a**
```
  C    U     U   UU    GUC          C    U     U   UU    GUC
   CCCA AGAAA AUU  CUAU              CCCA AGAAA AUU  CUAU
   GGGU UCUUU UAA  GAUG              GGGU UCUUU UAA  GAUG
UAA    U     U   UU               UAA    U     U   UU
```

**mir-8196b**
```
      U     U–   UU     UC              U     U–   UU    GUC
  CCCA AGAAA  AUU  CUAUG            CCCA AGAAA  AUU  CUAU
  GGGU UCUUU  UAA  GAUGU            GGGU UCUUU  UAA  GAUG
AA    U     UU   UU               AA    U     UU   UU    U
```

**mir-8197**
```
    U        CCAC     UC              U        CCAC     UC
  AG GCUUUGCU    CCAAC              AG GCUUUGCU    CCAAC
  UC UGGAACGA    GGUUG              UC UGGAACGA    GGUUG
AG  U        AAC–                 AG  U        AAC–
```

**mir-8198**
```
          U  –––    UGUU                    UUCA      U
 UUGAACAGU UC   AAUU               UUGAACAGU    AUUUGU
 AACUUGUCA AG   UUAA               AACUUGUCA    UAGGCA
A         U  GCA    U             A         ––––      UUAAU
```

**mir-8199**
```
  UCGGA       –A–   GAU             UCGGA       A  GGAU
       CAAUAUC   CUG                     CAAUAUC CU
       GUUGUAG   GAU                     GUUGUAG GA
AUAAAAG       GAA   G             AUAAAAG       –  AGAUG
```

**mir-81**
```
  C         C       G GA            C         C       G GA
   GGUUUUCAC GUGAUCU A               GGUUUUCAC GUGAUCU A
   UCGAAAGUG UACUAGA U               UCGAAAGUG UACUAGA U
UGA         C       G             UGA         C       G
```

**mir-8200**
```
                A      CC                         A      CC
  UGGCUCAAAUCUCC GUCAGA             UGGCUCAAAUCUCC GUCAGA
  ACCGAGUUUAGAGG CAGUCU             ACCGAGUUUAGAGG CAGUCU
CU              G                 CU              G
```

**mir-8201**
```
                    ACCU                              ACCU
  UCUGGAUCGAUAAUGUAA                UCUGGAUCGAUAAUGUAA
  AGACCUAGUUAUUACAUU                AGACCUAGUUAUUACAUU
AG                  AA            AG                  AA
```

**mir-8202**
```
       AA     AAA   GUG GU               AA     AAA   GUGUGU
     UG  CAGAA   GUC   U               UG  CAGAA   GUC
     AC  GUUUU   CAG   A               AC  GUUUU   CAG
CGUUA  AG     –A–   AG–           CGUUA  AG     –A–   AGA
```

| | | |
|---|---|---|
| *mir-8203* | C UUCA CA– C<br>UGAA AUCA GGG UUA<br>GCUU UAGU CCC AAU<br>UA A CAAC AUA | C UUCA C AC<br>UGAA AUCA GGG AUU<br>GCUU UAGU CCC UAA<br>UA A CAAC A AU |
| *mir-8204* | A C U UU<br>UGGUCUC C ACGCGU ACUCA<br>ACCGGAG G UGCGCG UGGGU<br>CUU C A U | A C U UU<br>UGGUCUC C ACGCGU ACUCA<br>ACCGGAG G UGCGCG UGGGU<br>CUU C A U |
| *mir-8205* | C UC U<br>UGG AGACU GU GAGGCUA<br>ACC UUUGA CA CUCCGAU<br>GC U CC – U | C UC U<br>UGG AGACU GU GAGGCUA<br>ACC UUUGA CA CUCCGAU<br>GC U CC – U |
| *mir-8206* | U UCA<br>UAUA AAUGUAAUCUGAAA<br>AUAU UUACAUUAGACUUU<br>GG C | U UCA<br>UAUA AAUGUAAUCUGAAA<br>AUAU UUACAUUAGACUUU<br>GG C |
| *mir-8207* | C C UU CA<br>UUGU CUCUUUUCU UCA UG<br>AGCA GAGAAAAGA GGU AC<br>AG A A U– | C C UUUGCA<br>UUGU CUCUUUUCU UCA<br>AGCA GAGAAAAGA GGU<br>AG A A UAC |
| *mir-8208* | CAG UU<br>UCCGCCCA UUGAACCAA<br>AGGCGGGU GACUUGGUU<br>UCA AAG | CAG UU<br>UCCGCCCA UUGAACCAA<br>AGGCGGGU GACUUGGUU<br>UCA AAG |
| *mir-8209* | A AA A<br>AAACGAAG AGAAGAAGA<br>UUUGCUUC UCUUCUUCU<br>CCC CC | A AA A<br>AAACGAAG AGAAGAAGA<br>UUUGCUUC UCUUCUUCU<br>CCC CC |
| *mir-8210* | UG C C GUG C AC<br>C UUCUUUC UU UCG CG<br>G AAGAAAG AA AGC GC<br>AAA A A AAA A | UG C C GUG C AC<br>C UUCUUUC UU UCG CG<br>G AAGAAAG AA AGC GC<br>AAA A A AAA A |
| *mir-8211* | A – UG<br>CUCGAGGC CGGUGAGC C<br>GAGCUUCG GCCGCUCG G<br>CAA A –– CA | A – C<br>CUCGAGGC CGGUGAGCUG<br>GAGCUUCG GCCGCUCGGC<br>CAA A A |
| *mir-8212* | AC A GU<br>UUGCUCAAAAAUU UUC AA<br>AACGAGUUUUUGA GAG UU<br>UC CA C | AC AAAGU<br>UUGCUCAAAAAUU UUC<br>AACGAGUUUUUGA GAG<br>UC CA CUU |
| *mir-82* | C U U A GA<br>GGUUUUC C GUGAUCU CA<br>CCGAAAG G UACUAGA GU<br>UGA U C – | C U U A GA<br>GGUUUUC C GUGAUCU CA<br>CCGAAAG G UACUAGA GU<br>UGA U C – |
| *mir-83* | U A UGA<br>ACUGAAUUUAUGUG GU CU<br>UGACUUAAAUAUAC CA GA<br>AA – C U | – ACUUGA<br>ACUGAAUUUAUGUG UGU<br>UGACUUAAAUAUAC ACG<br>AA C AU |
| *mir-84* | G A U AGA<br>UGAG UAGU UG AAUAUUGU<br>GCUC AUCA AC UUGUAACA<br>CG A – U C | G A U AGA<br>UGAG UAGU UG AAUAUUGU<br>GCUC AUCA AC UUGUAACA<br>CG A – U C |
| *mir-85* | C – G A AC<br>CGAUUUUUCAA UA UUUG A<br>GCUGAAAAGUU AU AAAC U<br>CGU U G A | C – G AAAC<br>CGAUUUUUCAA UA UUUG<br>GCUGAAAAGUU AU AAAC<br>CGU U G AU |
| *mir-86* | C UU A UC<br>UAAGUGAAU CU GCC CAG<br>AUUCGCUUA GA CGG GUC<br>CGG – CU – | C UU A UC<br>UAAGUGAAU CU GCC CAG<br>AUUCGCUUA GA CGG GUC<br>CGG – CU – |
| *mir-87* | U C U A CU<br>CGCCUGA ACUUU G CUCA C<br>GUGGACU UGAAA C GAGU G<br>CGU U – – – | U C U ACCU<br>CGCCUGA ACUUU G CUCA<br>GUGGACU UGAAA C GAGU<br>CGU U – – G |
| *mir-90* | C U – G AC<br>GGC UUCAA CGAC AUAUCA<br>CCG AAGUU GUUG UAUAGU<br>UCC U U – | C U – G AC<br>GGC UUCAA CGAC AUAUCA<br>CCG AAGUU GUUG UAUAGU<br>UCC U U – |
